# Structural evolution of the MTCH family of mitochondrial insertases

**DOI:** 10.64898/2026.02.17.705849

**Authors:** Taylor A. Stevens, Zhilin Luo, Camryn Lee, Masami Hazu, Erini G. Galatis, Alison J. Inglis, Alina Guna, Rebecca M. Voorhees

## Abstract

We demonstrate that MTCH2 is the defining member of a large family of mitochondrial outer membrane (OM) insertases. MTCH insertases are conserved across holozoa and have diverged from the solute carrier 25 transporters. The cryoelectron microscopy structure of the 33 kDa human MTCH2 revealed that evolution of its insertase activity required loss of a transmembrane helix, which created a lipid-accessible hydrophilic groove stabilized by its unique, structured C-terminus. Mutational analyses showed that MTCH insertase activity is attenuated, while experimental structures and reconstitution of hyperactive mutants demonstrated that the hydrophobicity, charge, and size of the residues that line its groove regulated MTCH function. Leveraging the MTCH2 structure, we identified the plant OM insertase, and proposed a universal mechanism for OM insertion across all kingdoms of life.

**Teaser:** Structure of human MTCH2 reveals conserved features necessary to maintain mitochondrial proteome integrity across eukaryotes.

## Background and Introduction

A critical step in the biogenesis of all integral membrane proteins is their integration into the lipid bilayer. Successful insertion requires the simultaneous shepherding of a hydrophobic element, often a transmembrane helix (TM), into the membrane coupled with the translocation of an associated soluble domain across the bilayer. The latter of which typically requires catalysis by a membrane protein insertase. To ensure the efficient and robust insertion of the full diversity of membrane proteins, eukaryotic cells have evolved multiple families of insertases that function at each membrane and specialize in specific classes of substrates (*1–3*).

An important site of membrane protein biogenesis is the mitochondrial outer membrane (OM). The OM is essential for mediating communication between the organelle and the cytosol (*4*), which relies on the accurate localization and insertion of a suite of α-helical membrane proteins. Because of their roles in regulating both mitochondrial dynamics and cellular processes such as apoptosis and the innate immune system, defects in OM protein biogenesis are associated with numerous human diseases including Parkinson’s disease, metabolic disorders, and many cancers (*5*, *6*). In mammals, the OM contains ∼150 α-helical membrane proteins, all of which are encoded by the nuclear genome, translated on cytosolic ribosomes, and then integrated directly into the OM (*7*).

We recently demonstrated that the human Mitochondrial Carrier Homolog 2 (MTCH2) functions as an OM insertase for many of these α-helical proteins in human cells (*8*), providing a biochemical explanation for its central role in mitochondrial biology and human disease (*9–11*). MTCH2 depletion affects diverse α-helical OM substrates including tail-anchored, signal-anchored, and multipass proteins (*8*). MTCH2 and its close paralog, MTCH1 (*12*), are diverged members of the solute carrier 25 (SLC25) family, which are transporters typically localized to the inner mitochondrial membrane (*13–15*). Many aspects of MTCH2 function are not understood, including disagreement about its topology (*12*), fundamental questions regarding the molecular basis of its insertase activity, and limited information about how broadly it is conserved. Moreover, how MTCH2 evolved from a family of transporters that move small molecules across the membrane to one that mediates insertion of proteins into the bilayer is not clear. Finally, the relationship between MTCH2 and its counterparts in fungus (Mim1/2; (*16*, *17*)) and protists (pATOM36; (*18*, *19*)), and identification of a putative OM insertase in plants, would provide insight into the evolution of OM protein biogenesis across eukaryotic kingdoms. Here we set out to use a combination of structure, biochemistry, and molecular biology to characterize how MTCH2 evolved to co-opt a solute carrier fold for insertion of α-helical proteins into the bilayer.

### The MTCH family of mitochondrial membrane protein insertases

Bioinformatic analysis identified a distinct SLC25 subclass, which includes both human MTCH1 and MTCH2, as well as thousands of other sequences widely distributed across the holozoan clade (Fig. 1A). Sequence comparison suggested that these MTCH2 homologs retain many features of the SLC25 family including highly conserved prolines within TM1, TM3, and TM5 (fig. S1A,B) (*13*). However, they have also lost several transporter elements, such as the characteristic salt bridges between helices, important for the alternating access mechanism used for solute transport. Further, the MTCH2 homologs all contain a highly diverged and shortened C-terminus compared to canonical SLC25s (fig. S1C,D). While all metazoans contain at least one MTCH2 homolog predicted to localize to their respective OM, vertebrates express two paralogs, MTCH1 and MTCH2, which have previously been shown to be synthetic lethal in human cells (*20*).

**Fig. 1.**
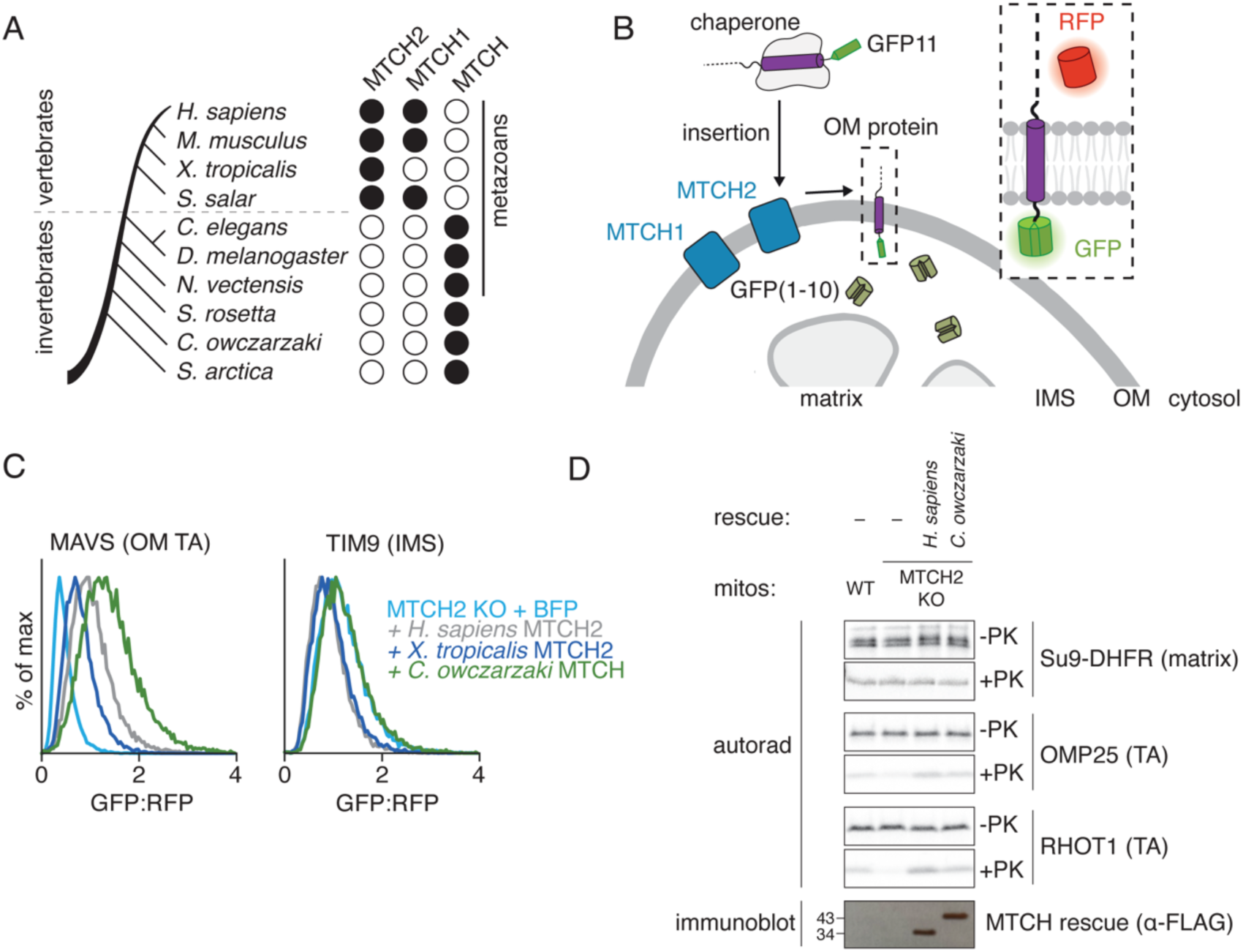
MTCH2 is the defining member of the MTCH family of insertases conserved across holozoa. **A)** To identify MTCH2 homologs across the holozoan clade, we queried Uniprot for sequences annotated with the panther family PTHR10780. Vertebrates expressed homologs of both MTCH1 and MTCH2, while invertebrates contained a single MTCH homolog, which cannot be unambiguously assigned as related to MTCH1 vs 2. The presence and type of the predicted homolog is indicated with a black circle, and a summary of the query results are in table S4. **B)** Schematic of the split-GFP fluorescent reporter system previously established to assess insertion into the mitochondrial outer membrane (OM) (*8*, *21*, *22*). Here, GFP(1-10) is localized to the inner membrane space (IMS) while GFP11 is appended to the protein of interest. Insertion into the OM in the correct orientation results in complementation and thereby GFP fluorescence. GFP11 containing reporters are encoded on the same open reading frame as a normalization marker, RFP, separated by a P2A ribosomal skipping sequence to specifically allow measurement of post-translational effects on reporter biogenesis. Using this system, we demonstrated that both MTCH1 and MTCH2 facilitated insertion of α-helical containing OM proteins in human cells (see also fig. S2). **C)** Insertion of the OM tail-anchored (TA) reporter MAVS-GFP11 and an IMS localized control, TIM9-GFP11 as described in (B) was assessed by flow cytometry in K562 MTCH2 KO cells expressing either a BFP control, human MTCH2 (*H. sapiens*), *X. tropicalis* MTCH2, or *C. owczarzaki* MTCH. GFP fluorescence, a proxy for reporter insertion, relative to our normalization control (RFP) was calculated and is displayed as a histogram (additional substrates and relevant controls are displayed in fig S3A). **D)** To directly assess insertion activity of the indicated MTCH homologs, we performed an *in vitro* insertion experiment using isolated mitochondria (fig. S3C). The indicated ^35^S-methionine-labelled substrates were translated in rabbit reticulocyte lysate and released from the ribosome by treatment with puromycin. These included the MTCH2 independent Su9-DHFR control (which contains the canonical TOM targeting sequence, Su9), and two MTCH2 dependent OM proteins, OMP25 (the native sequence) and RHOT1 (where the large cytosolic lumenal domain of RHOT1 is replaced with the smaller, globular VHP domain as described previously (*8*)). Each substrate was incubated with mitochondria isolated from WT or MTCH2 KO human K562 cells expressing either a mock control, 3xFLAG-tagged *H. sapiens* MTCH2, or 3xFLAG-tagged *C. owczarzaki* MTCH. Relative insertion was assessed using a protease protection assay and analyzed using SDS-polyacrylamide gel electrophoresis (SDS-PAGE) and autoradiography, before (–PK) and after (+PK) addition of protease. An immunoblot (3xFLAG) is displayed to indicate relative expression of the MTCH homologs.

To determine if these putative MTCH2 homologs could also function as membrane protein insertases, we tested if they could rescue α-helical OM protein insertion in a MTCH2 depleted human cell. For this, we used a split-GFP reporter assay that specifically monitors OM insertion (*8*, *21*, *22*) (Fig. 1B). Consistent with earlier results (*8*, *12*), we observed that MTCH1 could partially rescue loss of MTCH2 for several α-helical OM substrates (fig. S2A,B). Mammalian mitochondria also express a third SLC25 member, SLC25A46, in their outer membrane, which does not appear to play a role in insertion, and could not rescue loss of MTCH2 (*15*) (fig. S2C,D). Remarkably, we found that several MTCH2 homologs, including those from *X. tropicalis* and the single-celled organism *C. owczarzaki*, rescued a MTCH2 deletion phenotype using our ratiometric fluorescent reporter assay (Fig. 1C, fig. S3A,B). To confirm that this effect was at the level of insertion, we purified human mitochondria from a MTCH2 knockout cell line exogenously expressing either human MTCH2 or the *C. owczarzaki* MTCH2 homolog. Even the distantly related homolog was able to mediate the integration of human α-helical proteins into the OM in vitro (Fig. 1D, fig. S3C), suggesting an exceptional level of conservation across what we will now refer to as the MTCH family of membrane protein insertases.

### The structure and evolution of human MTCH2

To better understand how this family of proteins may have evolved their insertase activity from an ancestral transporter fold, we sought to analyze the structure of a representative member of the MTCH family. While the arrangement of some of the transmembrane helices could be potentially inferred due to their similarity to the SLC25s, the conserved C-terminus, unique to the MTCH family, and thereby likely of functional significance, cannot be confidently predicted by AlphaFold (fig. S1D) (*23*, *24*). Additionally, while canonical SLC25 carriers alternate between conformations open to the cytosol or matrix (*25*, *26*), MTCH2, and all other MTCH homologs, lack conserved residues stabilizing either of these conformations. It is not clear whether MTCH2 adopts multiple conformations, and indeed there is disagreement in the literature about its number of TMs. Therefore, an experimental structure is required to unambiguously assess the molecular basis for insertase activity. For structural analysis we chose to focus on human MTCH2, since it is the best characterized member of the family and because an experimental structure could be therapeutically valuable due to the close association between MTCH2 and human disease (*27*).

Due to its small size (33 kDa) structural determination of MTCH2 by cryoelectron microscopy (cryo-EM) required a strategy to rigidly add asymmetric mass as a fiducial marker for particle alignment. We tested several fusion strategies at various positions within MTCH2. These included fusions with BRIL, which has been successfully used to determine other membrane protein structures and can be complexed with the established anti-BRIL antibody (*28*). While we were able to determine an initial low-resolution structure using a BRIL fusion strategy (fig. S4), preliminary density maps suggested that further optimization would be required to allow *de novo* building of the C-terminal domain of MTCH2. We instead leveraged tools from the recently determined structure of the uncoupling protein (UCP1) (*29*), an SLC25 transporter localized to the inner mitochondrial membrane. Jones et al. generated a nanobody that binds to a short segment of the loop between TMs 4 and 5 (loop 4/5), which is positioned in the innr membrane space (IMS). We determined that the homologous cytosolic loop in MTCH2 was of similar length, its sequence was poorly conserved amongst MTCH homologs, and mutations to this loop did not affect MTCH2 insertase activity in human cells (fig. S5A,B). We therefore reasoned that we could replace these residues in MTCH2 with the nanobody recognition sequence from the UCP1 loop 4/5 without altering MTCH2’s structure or function (fig. S5C,D). Indeed the MTCH2-loop fusion was stably expressed, remained competent for substrate insertion (fig. S5E,F), and could be purified as a homogeneous and monodispersed sample in the detergent UDM. We then generated a version of the UCP1 nanobody that could be recognized by a universal anti-nanobody Fab (NabFab) (*30*) to allow purification of a 112 kDa tetrameric complex of MTCH2, the modified UCP1 nanobody, the NabFab, and finally an anti-Fab nanobody (*31*) (Fig. 2A).

**Fig. 2.**
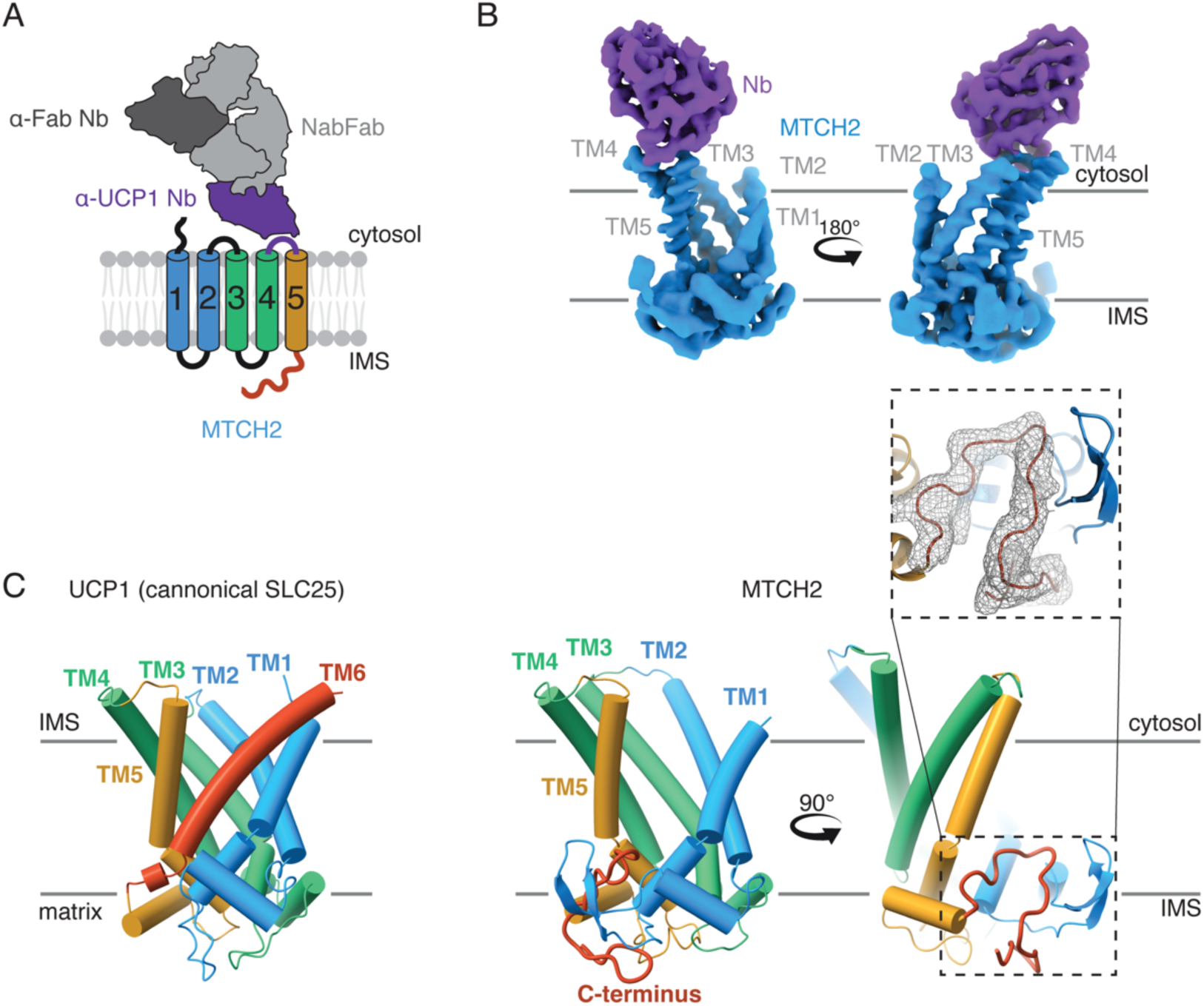
Structure of the 33 kDa human MTCH2 determined by single particle cryo-EM. **A)** Schematic of the strategy used to assemble the tetrameric complexes of human MTCH2 used for structural determination. As described in detail in Fig. S5C,D, the *H. sapiens* MTCH2 sequence was modified to include an epitope that is recognized by the α-UCP1 nanobody (Nb) pMb65 (*29*). pMb65 was modified to permit binding of the universal NabFab (*30*), which was itself bound to an additional α-Fab Nb (*31*). **B)** EM density map of human MTCH2 (blue) and the α-UCP1 Nb (purple) determined to an overall resolution of 3.6 Å. All five TMs were unambiguously resolved and the resolution was sufficient to allow *de novo* building of the C-terminal domain of MTCH2, which is unique to the MTCH insertase family (fig. S1C). **C)** Comparison of MTCH2 with the SLC25 transporter UCP1 (PDB ID: 8HBV, left (*81*)) highlights both the similarities and differences between the MTCH family insertases and their ancestral transporters, including the loss of a 6^th^ TM. Inset is the refined model and EM density for the C-terminus of MTCH2, whose structure was incorrectly predicted by AlphaFold. Its positioning directly below the cavity left by loss of the 6^th^ TM, and role in stabilization of MTCH2 (fig. S8D,E) suggests it may have co-evolved with the insertase activity of the MTCH family.

Using a standard data collection and computational processing pipeline, we determined the structure of human MTCH2 to an overall resolution of 3.6 Å (Fig. 2B, fig. S6-7, Table S1), sufficient to allow unambiguous fitting of the TM helices and *de novo* building of the C-terminal domain (Fig. 2C) (*23*, *24*). Comparison of the EM density from the UCP1-fused and BRIL-fused datasets suggests that the overall shape and conformation is not impacted by either modification (fig. S7C,D). Given that the size of loop 4/5 is similar in most SLC25s, but the sequence is typically not critical for function (*32*), it is likely that this UCP1-fusion strategy is universally applicable to enable structure determination of diverse members of this essential transporter family and all MTCH insertases.

Comparison with the UCP1 structure definitively indicated that unlike a canonical SLC25, MTCH2 contained only five TMs (Fig. 2C). This is consistent with our analysis of its topology in human cells (fig. S8A,B). Instead, the C-terminus of MTCH2, conserved across the MTCH family of insertases (fig. S1A), adopts a defined fold that stabilizes the cavity formed by loss of this sixth TM. Indeed, the deletion of the MTCH2 C-terminal residues 279-303 resulted in a marked destabilization of the protein in cells (fig. S8D,E). Therefore, unlike a transporter, which forms an enclosed hydrophilic pore through the membrane, MTCH2 contains an exposed hydrophilic vestibule within the bilayer (Fig. 3A). Further analysis revealed that the structure of MTCH2 adopts a cytoplasm-open conformation (fig. S9A), despite lacking the conserved salt bridges that stabilize this conformation in canonical SLC25 carriers (fig. S1B). Mutations restoring these salt bridges do not reduce MTCH2s function (fig. S9B). While we cannot rule out transient conformational changes, the presence of a cytosol-accessible cavity with a lateral opening to the lipid bilayer is consistent with a model in which nascent TMs are trafficked from the cytosol to MTCH2 for OM insertion. Notably, the cavity is also lined by a series of bulky hydrophobic residues that are conserved across MTCH family homologs, many of which are within the C-terminal domain that could only be visualized in the experimental structure (fig. S10,B). The MTCH family of insertases is yet another example of the convergent evolution that has led many evolutionarily unrelated insertase families to utilize similar hydrophilic grooves to mediate insertion (fig. S10B). This includes the universally conserved Oxa1/YidC superfamily of insertases (*33*, *34*) such as EMC (*35*), GET1/2 (*36*), and YidC (*37*) (fig. S10C). In all cases, positioning of a hydrophilic surface within the membrane is thought to decrease the energetic barrier of translocation of soluble domains across the hydrophobic core of the bilayer, thereby catalyzing membrane protein insertion (*1*).

**Fig. 3.**
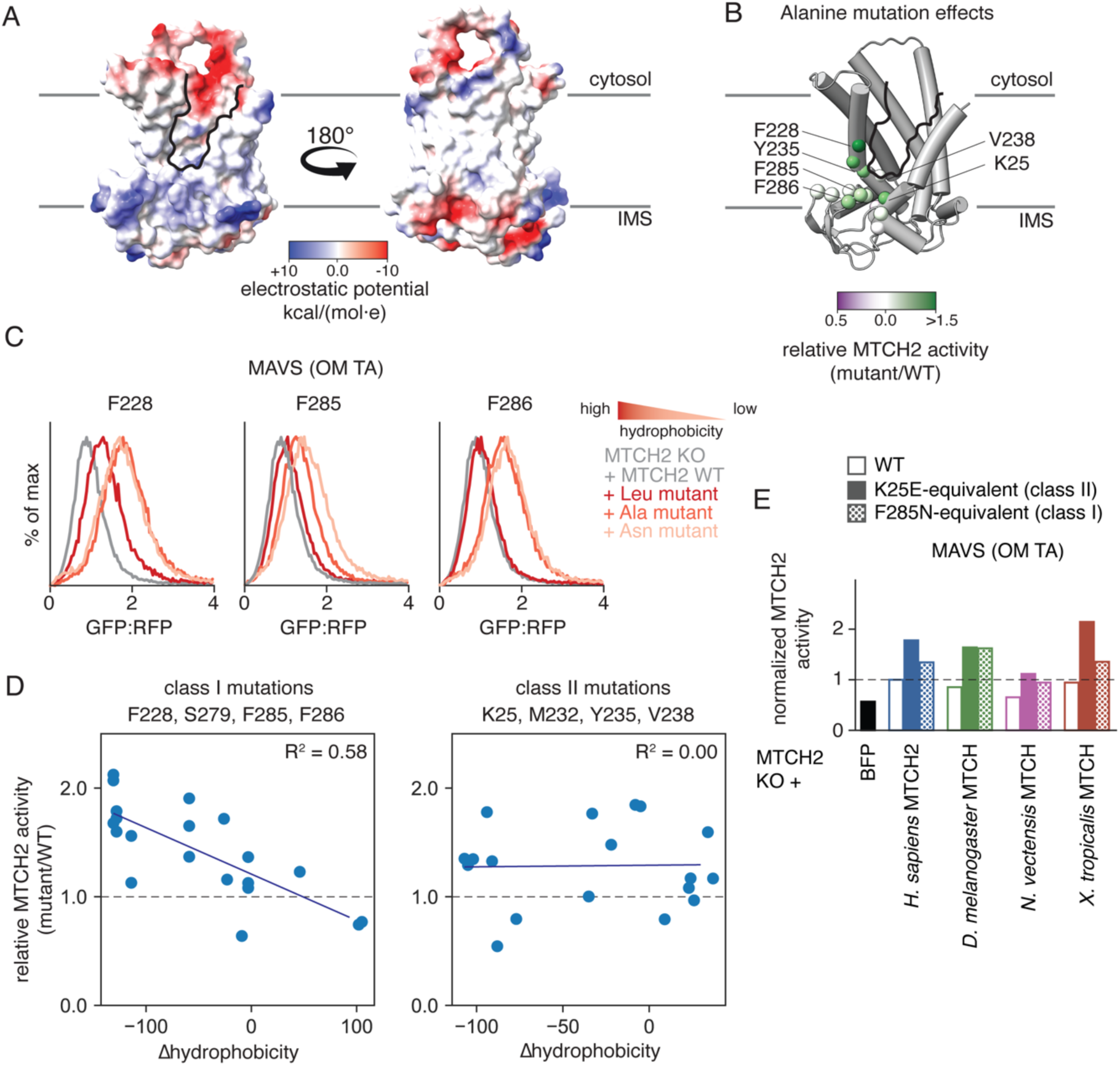
The activity of MTCH2 is attenuated by residues surrounding its hydrophilic groove. **A)** Coulombic electrostatic potential was calculated using ChimeraX and mapped onto a surface visualization of the experimentally determined model of MTCH2. **B)** To test the role of MTCH2’s hydrophilic groove in insertion (outlined in black), we performed alanine scanning mutagenesis on the indicated residues surrounding the groove. The ability of each of these alanine mutants to rescue a MTCH2 KO phenotype on MAVS insertion, when compared to similar expression of WT MTCH2 (additional substrates and relevant controls are displayed in fig. S11B), was assessed using the ratiometric reporter described in Fig 1B. The relative activity of each alanine mutant compared to WT MTCH2 (calculated as GFP:RFP_mut_/GFP:RFP_wt_) was mapped onto the corresponding site using the indicated scale. Darker green indicates positions where mutations leads to increased insertion compared to wild type MTCH2 that cannot be explained by changes in expression (fig. S11C). **C)** Eight positions lining the MTCH2 hydrophilic groove were selected for more extensive mutagenesis (Phe, Leu, Gln, Glu and Arg in addition to Ala). Relative insertion of MAVS and OMP25 and a MTCH2 independent control was assessed as in (B) (additional substrates and relevant controls are displayed in fig. S13). Highlighted here are the effects of a subset of mutations for three representative positions lining the hydrophilic groove. Curves are colored by the hydrophobicity of the amino acid introduced in each mutant. **D)** (left) Data summarizing the effect of mutations (Phe, Leu, Ala, Gln, Glu, Arg) at four MTCH2 positions (F228, S279, F285, and F286) on MAVS insertion as described in (C). Displayed is the relative MTCH2 activity (GFP:RFP_mut_/GFP:RFP_wt_) plotted against the change in hydrophobicity of the mutated amino acid (ΛHydrophobicity=hydrophobicity_aa mut_-hydrophobicity_aa WT_) (*77*). These positions, referred to as Class I, share a correlation between the hydrophobicity of the mutation and activity of MTCH2 in insertion. (right) The same analysis was carried out with four separate positions (K25E, M232, Y235, V238), referred to as Class II, where there is no strong correlation between hyperactivity and hydrophobicity (additional substrates displayed in fig. S14A). **E)** To test whether MTCH2 activating mutations are conserved across the MTCH family, we measured the activity of a panel of MTCH homologs and their respective Class I and II mutants. Using the same rescue assay described in Fig. 1C, we tested the relative activity of a BFP control, WT, K25E, and F285N human MTCH2, and WT and mutant MTCH homologs from *C. owczarzaki*, *D. melanogaster*, or *N. vectensis* (mutations equivalent to K25E and F285N in human MTCH2) on MAVS insertion. Relative activity was calculated as GFP:RFP_mut_/GFP:RFP_wt_ (additional substrates and relevant controls are displayed in fig. S15).

Certainly, loss of this sixth TM is one of the most striking evolutionary adaptations between the SLC25 transporters and the MTCH insertase family. Indeed, SLC25A46, though also localized to the OM like MTCH1 and 2, cannot mediate membrane protein insertion (fig. S2A,C) and is predicted to contain six TMs (*23*, *24*). To test whether this evolved hydrophilic vestibule was important for MTCH family insertase function, we performed alanine scanning mutagenesis to residues around this region (Fig. 3B, fig. S11A,B). While we expected that mutations would primarily decrease MTCH2 activity, many of the mutations increased MTCH2 mediated insertion in cells. The observed activation was independent of effects on the stability or expression of MTCH2 (fig. S11C). Many of these mutations are localized to the C-terminal domain of MTCH2 which appears to have co-evolved with loss of the sixth TM, likely both critical steps during its evolution from a transporter to an insertase.

One possibility is that these mutations cause increased insertion by disrupting interactions between MTCH2 and a constitutively bound inhibitory small molecule or protein. If this were the case, depletion of this putative regulatory molecule would phenocopy the activating effect. However, knockdown of known interaction partners of MTCH2, ARMC1 and DNAJC11 (*38*), did not activate MTCH2 function in K562 cells (fig. S12A-D). We therefore investigated whether these mutations could instead alter the intrinsic insertion propensity of MTCH2 itself, perhaps by relieving an inhibitory effect of certain amino acids towards insertion.

### Hydrophobic residues impede MTCH family insertase activity

To examine the mechanistic basis for this activation, we performed a more extensive mutational analysis aimed at identifying how hydrophobicity, size, and charge affect MTCH2 function. We selected a panel of eight positions, focusing on functionally important regions along the hydrophilic groove identified through the preliminary alanine scanning experiment. Using multiple substrates, we tested the activity of MTCH2 in human cells when one of six amino acids was present at each position (Phe, Ala, Leu, Asn, Glu, Arg) (fig. S13 A-D). For many of the sites tested, we noted a negative correlation between hydrophobicity and insertion propensity across multiple MTCH2 substrates (Fig. 3D, fig. S14A). For example, mutation of residues F228, F285, and F286 to increasingly more polar amino acids (i.e. Ala, Asn) generally resulted in a commensurate activation of MTCH2 in human cells (Fig. 3C). Remarkably this effect held across four of the eight sites tested, suggesting a shared mechanism for activation at these positions, which all line the hydrophilic vestibule. Our analysis suggests that in addition to hydrophobicity, charge and amino acid size can also contribute to MTCH2 activity.

Notably though, this trend was not universal, such as position Y235, for which MTCH2 becomes more active upon mutation to leucine. Indeed, we found a second class of mutations where there is no correlation between change in hydrophobicity and MTCH2 insertion activity (Fig. 3D). Corresponding mutations across the MTCH homologs of this second class also resulted in a similarly activating effect (Fig. 3E, fig. S15). We therefore postulated there were two distinct mechanisms for activation of MTCH2. Further, the features of the hydrophilic groove and the residues that line it seem to be conserved across the MTCH family. We propose that these results are most consistent with an apparent attenuation, or tuning, of MTCH family insertase activity across holozoans, which is altered by these mutations.

### Structural analysis defines two strategies to achieve hyperactivation of MTCH2

To understand the molecular basis for this activation, we chose representative constructs for each of the two classes of activating mutations: (Class I) those where a decrease in hydrophobicity along the groove leads to increased MTCH2 activity; (Class II) those that show no correlation between MTCH2 activity and hydrophobicity. For the former category, we selected a double mutant in which two conserved aromatic hydrophobic residues within its C-terminal domain that line the groove were mutated to smaller polar amino acids. We determined the structure of this mutant, MTCH2^F285N, F286N^, to 3.2Å resolution using the same UCP1-nanobody strategy utilized for the wildtype structure (Fig. 4A, fig. S16, fig. S17, Table S1). Superposition with wildtype MTCH2 identified no major conformational changes (Fig. 4B), suggesting that activation may instead be explained by how mutations to these residues directly change the size, shape, and electrostatic potential of the hydrophilic groove (Fig. 4E). We anticipate that all of the Class I mutations would behave in a similar manner to MTCH2^F285N, F286N^, where mutation of a hydrophobic residue is also predicted to increase the width of the hydrophilic groove (Fig. 4E), resulting in a corresponding increase in insertion activity.

**Fig. 4.**
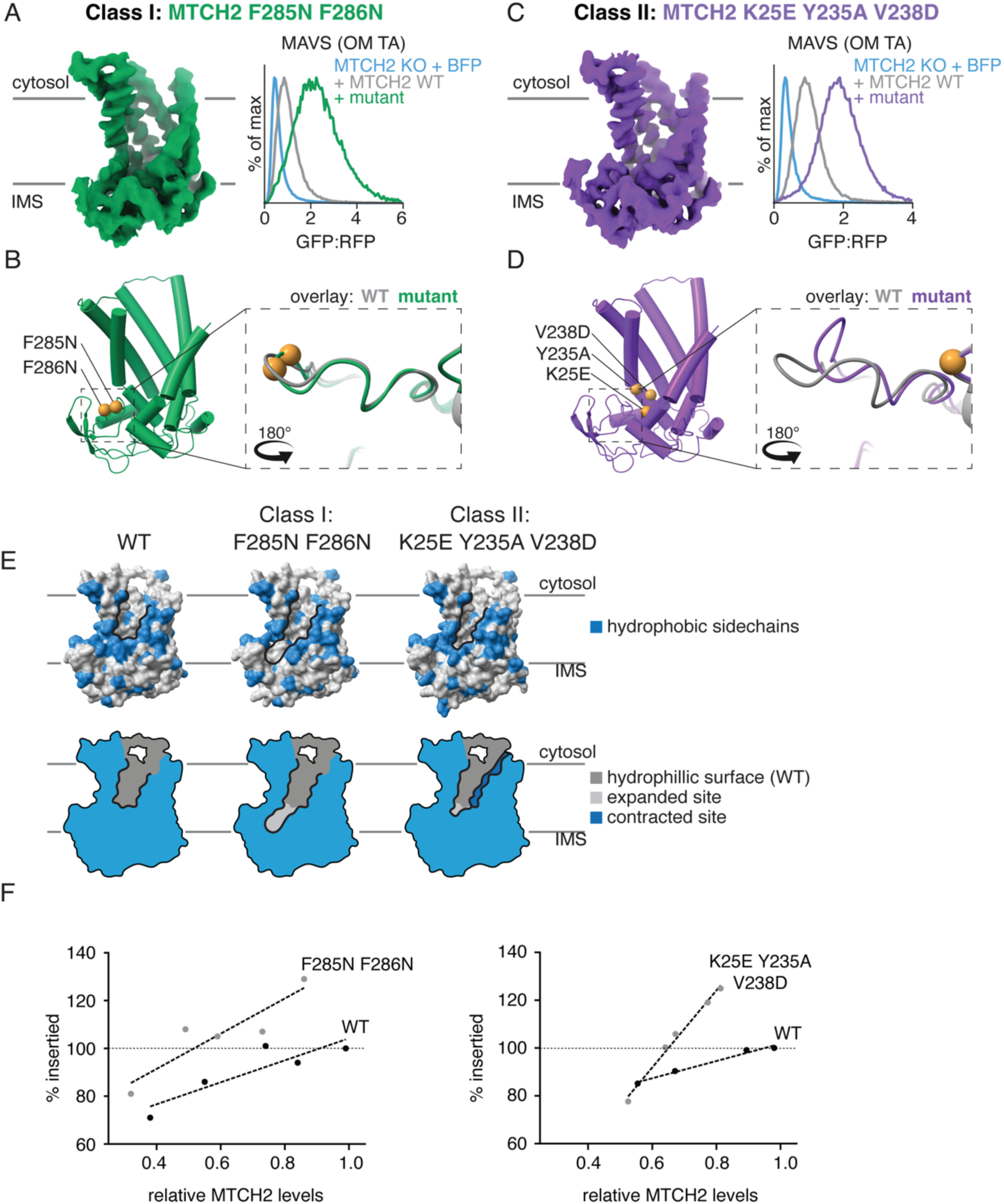
The structural basis for two distinct mechanisms of MTCH2 hyperactivation. **A)** (left) Cryo-EM density of a representative Class I MTCH2 hyperactive mutant (MTCH2^F285N, F286N^), which displays an anti-correlative relationship between hydrophobicity and insertion activity (Fig. 3D, fig. S14B). Structural determination was performed using a similar strategy for the wildtype MTCH2 as shown in Fig. 2A. (right) Activity of WT vs MTCH2^F285N, F286N^ in a MTCH2 KO K562 cell line, compared to a BFP control for MAVS insertion (additional substrates and relevant controls are displayed in fig. S14C, D). **B)** (left) The experimentally determined model of hyperactive MTCH2^F285N, F286N^ is displayed as a cartoon. The location of the two mutations, positioned along the hydrophilic groove, is highlighted (gold spheres). (right) Superposition of the WT (gray) and MTCH2^F285N, F286N^ (green) models, highlighting the C-terminal domain. **C, D)** As in (A, B) for a mutation representing the Class II MTCH2 activating mutants (MTCH2^K25E, Y235A, V238D^). At these three positions, we observed no correlation between hydrophobicity and MTCH2 activity, in contrast to the MTCH2^F285N, F286N^ mutant (Fig. 3D). MTCH2^K25E, Y235A, V238D^ induced a conformational change of residues 283-286 within the C-terminal domain of up to 7 Å, which was not observed for the MTCH2^F285N, F286N^ mutant. This movement directly repositions residues F285 and F286 (already shown to be important for MTCH2 activation in (B)) away from the hydrophilic groove (fig. S19C). **E)** (top) Displayed is the space filling representations of the experimentally determined structures of WT, MTCH2^F285N, F286N^, and MTCH2^K25E, Y235A, V238D^ with bulky hydrophobics (Phe, Leu, Met, Ile, Trp, or Tyr) shown in blue, and the hydrophilic groove outlined in black. (bottom) A simplified 2D representation highlights how these two classes of mutants result in changes to the hydrophilic groove of MTCH2 that correlate with changes in MTCH2 activity. Extensions to the hydrophilic groove when compared to WT are highlighted in light gray, while narrowing is highlighted in dark blue. The two classes of activating mutations use distinct mechanisms for achieving MTCH2 activation: (Class I) by either directly mutating hydrophobic residues that line the groove and apparently impede insertion or (Class II) by inducing a conformational change that then repositions these same hydrophobic residues. **F)** Wildtype or the indicated MTCH2 mutants were reconstituted into proteoliposomes at varying concentrations (fig S20). The MTCH2 dependent tail-anchored protein OMP25 was translated in rabbit reticulocyte lysate in the presence of ^35^S-Methionine and tested for insertion in the indicated proteoliposomes using a protease protection assay as previously described (*8*). The resulting protected fragment was immunoprecipitated using a C-terminal 6x-HIS tag and quantified using autoradiography. % inserted was calculated relative to the maximum insertion observed for wildtype MTCH2 at its highest concentration. Corresponding immunoblots and autoradiographs are shown in fig. S20.

To study the second mechanism used by the Class II mutants for MTCH2 activation, we combined three mutations identified through our systematic analysis into a triple mutant (MTCH2^K25E, V238D, Y235A^) that resulted in a comparable stimulation of insertion as MTCH2^F285N, F286N^ (Fig 4C, fig. S14C-E). Using the same UCP1-nanobody strategy, we determined the structure of MTCH2^K25E, V238D, Y235A^ to 3.1Å resolution (Fig. 4C, fig. S17, fig. S18, Table S1). In contrast to our Class I mutant, we observed a significant conformational change in the C-terminus of MTCH2^K25E, V238D, Y235A^ when compared to the wildtype structure (Fig. 4D, fig. S19B). This conformational change is characterized by a sharp kink in the peptide backbone around S283 and a ∼7 Å shift in the Cα positions of F285 and F286, themselves Class I mutants, repositioning them away from the opening of the hydrophilic crevice towards F228 and M232 on TM5 (fig. S19C, movie S1). This conformational change could result from the combined effects of the charge modification at positions 25 and 238 and the removal of the bulky side chain at position 235. This structure provides strong evidence for the inhibitory effect of hydrophobic residues, including F285 and F286, which can be relieved either by their direct mutation or repositioning. Highlighting the importance of these experimental structures, we found that the observed conformational change is not computationally predicted, consistent with earlier work showing that AlphaFold cannot predict mutation-induced conformational changes (*23*, *24*) (fig. S19D).

To test whether these changes in insertion could be ascribed to intrinsic changes in MTCH2 activity, we purified MTCH2^F285N, F286N^ and MTCH2^K25E, V238D, Y235A^ and optimized conditions for their reconstitution into liposomes alongside wildtype MTCH2. In this purified reconstituted system, we queried the insertion of the TA protein OMP25 into proteoliposomes by either wildtype MTCH2 or these two hyperactive mutants using a protease-protection assay as previously described (*8*). At matched MTCH2 levels, insertion was achieved at greater efficiency by both mutants in this minimal component system (Fig. 4F; fig. S20). We therefore concluded that the biophysical properties of the hydrophilic groove directly impact the intrinsic insertion propensity of MTCH2.

Taken together, both mutant structures highlight the importance of the positioning of hydrophobic amino acids lining the base of the hydrophilic crevice, which impede MTCH2’s insertase activity. While MTCH2^F285N, F286N^ directly targets two such residues, MTCH2^K25E, V238D, Y235A^ repositions these same residues away from the base of the hydrophilic groove to achieve its effect. The end result in either case is an expansion at the base of the hydrophilic groove (Fig. 4E), demonstrating the critical structural features of MTCH2’s hydrophilic groove for its activity, and revealing a concordant strategy for activation that is conserved across MTCH family insertases (Fig. 3E, fig. S15).

### Convergent evolution across OM insertases

While nearly all eukaryotic cells contain mitochondria, we could not identify MTCH family homologs outside of holozoa (Fig. 5A). It has previously been shown that in yeast, α-helical protein insertion into the OM relies on two single-pass proteins, Mim1 and Mim2, which are conserved across fungi (*16*, *17*). Conversely, trypanosomes such as T. brucei rely on the integral OM protein pATOM36 to perform a similar function (*18*, *19*). Plants however, have no known OM insertase.

**Fig. 5.**
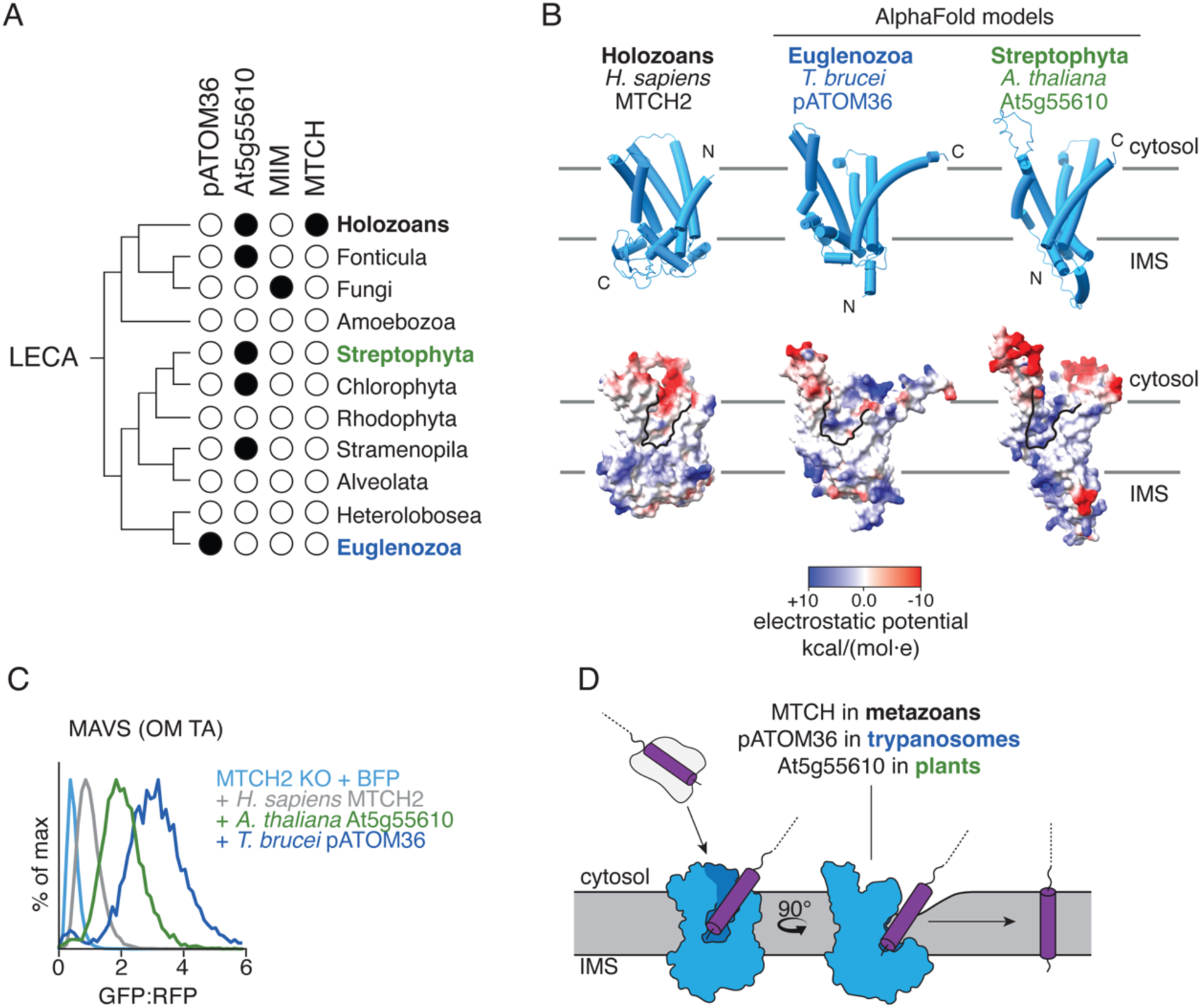
Convergent evolution of mitochondrial OM insertases across all kingdoms of life. **A)** Families of OM insertases were identified by querying Uniprot for sequences annotated with panther families PTHR10780 (MTCH), PTHR28241 (Mim1), PTHR36074 (At5g55610), or the interpro family IPR043645 (pATOM36) across eukaryotic lineages and mapped onto a species tree relative to the last eukaryotic common ancestor (LECA) (*76*). The presence of the putative insertase is indicated with a black circle (table S5). **B)** Comparison between the experimentally determined structure of MTCH2 (left) and AlphaFold-predicted models of *T. brucei* pATOM36 (middle) and *A. thaliana* At5g55610 (*23*, *24*) (right). AlphaFold model confidence metrics are shown in Table S6. Cartoon representations of the structures (top) are shown along with coulombic electrostatic potential mapped onto a space-filling model (bottom) with the respective hydrophilic clefts outlined in black. Topology of each insertase is based on split-GFP experiments shown in fig. S8B and S22A. Residues 1-85 and 1-60 of pATOM36 and At5g55610 respectively, which are predicted with low confidence were omitted for simplicity. **C)** To test whether At5g55610 functioned as an insertase, we tested its ability to rescue a MTCH2 KO phenotype in human cells. We expressed either a BFP control, *H. sapiens* MTCH2, *A. thaliana* At5g55610, or *T. brucei* pATOM36. Their effect on the human OM TA MAVS was measured by flow cytometry and is displayed as a histogram (additional substrates and relevant controls are displayed in fig. S22). **D)** A universal model for insertion of α-helical proteins into the mitochondrial OM across all species. Conserved, hydrophilic grooves, along with a likely contribution of local membrane thinning, are utilized to decrease the energetic barrier for translocation of IMS-localized soluble domains across the bilayer.

We exploited our experimental structure of MTCH2 to perform structural homology analysis in plants using a proteomics dataset from the OM of *A. thaliana* (*39*). We identified a protein, At5g55610, which was predicted by AlphaFold to contain five TMs (*23*, *24*), a hydrophilic groove within the lipid bilayer, and was conserved across plant species (Fig. 5B, fig. S21). While At5g55610 is not an SLC25 carrier, its AlphaFold-predicted model is strikingly similar to that of pATOM36. Further, it has been previously shown to localize to the OM by microscopy in *A. thaliana* (*39*). We found that At5g55610 and pATOM36 both localized to the OM in human cells, and experimental analysis suggested that both have an inverse topology when compared to MTCH2, with their C-terminus localized to the cytosol and N-terminus in the IMS (fig. S22A). Unlike MTCH and Mim1/2, At5g55610 and pATOM36 homologs are sporadically found throughout several distant eukaryotic lineages (Fig. 5A). One possible explanation is that this family emerged during the early evolution of plants and was acquired by other lineages through secondary endosymbiosis or horizontal gene transfer (*40*).

To determine if At5g55610 displayed insertase activity, we leveraged our ratiometric fluorescent reporter system in human cells to test if the protist, and putative plant insertases could rescue loss of MTCH2. Indeed, both insertases can not only express and fold in human mitochondria but can also functionally replace MTCH2 for insertion of human OM proteins (Fig. 5C, fig. S22B,C), illustrating a remarkable example of convergent evolution. In fact, at similar, or lower expression levels, protist pATOM36 and At5g55610 appear to be significantly more active than human MTCH2, emphasizing the apparent attenuation of human MTCH2 activity.

## Discussion

The endosymbiotic origin of mitochondria provides an explanation for the presence of several universally conserved β-barrel proteins in the OM (*41*, *42*). However, their increasingly complex roles in eukaryotic cells necessitated evolution of a nuclear encoded class of mitochondrial OM α-helical proteins. Unlike their β-barrel counterparts, machinery for α-helical insertion was not inherited from bacteria, suggesting that the first α-helical proteins relied on spontaneous insertion into the OM (*43*). The function of an OM insertase may have become necessary to better regulate an expanding OM proteome, or to enable greater plasticity in the OM composition of complex multicellular organisms. Such machinery appears to have evolved independently three times, resulting in MTCH in holozoa (*8*), pATOM36 in plants (*18*, *44*), and Mim1/2 in fungus (*16*, *17*). While these three families of insertases have no sequence similarity, all converged on similar structural features to mediate integration into the bilayer (*45*). In vertebrates, where an additional level of regulation is required to ensure adaptation and tuning in diverse cell types, there are two paralogs of the MTCH family, MTCH1 and MTCH2.

In holozoa, the MTCH family of insertases has evolved from an SLC25 carrier protein to utilize a qualitatively similar mechanism for membrane integration as the widely conserved Oxa1 Superfamily of insertases (*33*, *34*). In particular, we have experimentally shown that MTCH activity relies on a conserved hydrophilic groove within the bilayer, produced by loss of a TM and stabilized by evolution of a conserved C-terminal domain. We propose that the MTCH hydrophilic groove primarily serves two purposes. First, it provides a hydrogen bonding surface within the membrane that can interact with the soluble domain of a substrate as it transverses the hydrophobic core of the bilayer (*46*). Second, numerous experimental and molecular dynamics studies have demonstrated that these grooves induce local membrane thinning (*35*, *47–51*). Together, these effects decrease the energetic barrier for translocating a soluble domain across the membrane, catalyzing membrane insertion (Fig. 5D) (*3*). These hydrophilic grooves are further able to scramble lipids, which may be an additional important facet of these insertases’ function (*52*, *53*). Molecular dynamics simulations of MTCH2’s scramblase activity suggested that many of the same hydrophobic C-terminal residues we show regulate its protein insertion activity are also critical for its scramblase activity (F228, M232, F285, F286). It is therefore likely that MTCH2’s scramblase and insertase functions rely on a qualitatively similar mechanism, as has been previously proposed for other systems (*54*).

However, mutations to the hydrophilic groove of several MTCH family homologs resulted in increased insertase activity. While not common amongst insertases, attenuation of basal activity has been routinely observed for other types of membrane transporters such as ion channels (*55*). Structural analysis of two distinct hyperactive mutants of MTCH2 suggested that the biophysical and structural properties of its hydrophilic groove were closely correlated with its activity. Similar principles would apply to any membrane integrase that must provide a path for substrates across the bilayer. One potential explanation for the observed attenuation of MTCH activity is to tune or inhibit the intrinsic pathway for apoptosis, which plays a unique function in metazoans and is regulated by MTCH2 (*8*, *56*). Attenuation of MTCH2’s activity may also suggest its function is regulated, either through binding of small molecules or proteins (such as ARMC1 (*38*)), which could act by directly blocking the hydrophilic crevice or modulating the C-terminal conformation. Indeed, we found that overexpression of ARMC1 resulted in partial inhibition of MTCH2’s insertase activity via a mechanism dependent on its C-terminal domain (fig. S24). Such regulation could enable rapid remodeling of the OM proteome in a tissue specific manner or in response to metabolic or signaling cues. Given MTCH2’s central role in regulating the composition of the OM, understanding the molecular details of its activity opens the door to development of therapeutics that critically modulate MTCH2 activity as a strategy to treat human disease.

## Materials and Methods

### Plasmids and antibodies

All individual plasmids are listed in Table S2-3 and are available upon request. The majority of plasmids used for expression in K562 cells are in a lentiviral vector with a UCOE EF-1α-promoter and WPRE element as previously described (*57*) (Addgene #135448). Reporter constructs used in the split GFP complementation system were fused to the eleventh strand of GFP (GFP11: RDHMVLHEYVNAAGIT) at the IMS-localized termini of the reporter (*58*, *59*). In some cases, a GS linker was added to ensure 10 amino acids of separation between the transmembrane domain (TM) and GFP11, which we had previously found to be important for sterically allowing complementation with GFP(1–10) off of the lipid bilayer. To monitor mitochondrial outer membrane (OM) insertion of GFP11 tagged substrates (*22*), we used a construct encoding an inner membrane space (IMS) targeting sequence containing MICU1 residues 1-60 fused with GFP(1–10) (*21*). In order to specifically query post-translational processes, such as protein insertion, the red fluorescent protein mCherry (referred to as RFP throughout the publication for brevity) was expressed along with the GFP11-fused reporters separated by a P2A ribosomal skipping sequence, yielding two polypeptides from a single open reading frame (*60–63*).

All MTCH2 rescue experiments were performed using a UCOE EF-1α-BFP-P2A-MTCH2 WPRE construct, to allow analysis of only the transduced cell population (i.e. BFP+ cells) by flow cytometry. The same strategy was used for pATOM36 family rescue experiments. Point mutations to human MTCH2 or MTCH homologs were introduced via Gibson assembly with the mutations designed into the primer overhangs. UCOE EF-1α-BFP-P2A-MTCH1 WPRE rescue constructs included an N-terminal 9x GlySer linker.

For CRISPRi knockdown experiments with a single guide, sgRNAs were ordered as a pair of complementary oligonucleotides (Integrated DNA Technologies) and ligated between the BstXi and BlpI sites in the pU6-sgRNA EF-1α-Puro-T2A-BFP vector (Addgene, #84832) (*64*). To increase knockdown efficiency, particularly for MTCH1, two sgRNAs were cloned into a programmed dual sgRNA guide vector (Addgene #140096) (*65*). To enable simultaneous CRISPRi knockdown and expression of BFP-tagged rescue constructs, the BFP was removed from the CRISPRi knockdown plasmid in some experiments, while retaining the puromycin resistance marker to allow selection of cells transduced with the sgRNA-expressing plasmid. All sgRNA sequences and backbones are listed in Table S3.

The SP64 vector was used to generate PCR templates for *in vitro* transcription reactions for subsequent translation in rabbit reticulocyte lysate (RRL) (*66*). All OM substrates included an N-terminal 3xFLAG and C-terminal 6xHis tags, the latter of which was used to immune-precipitate a proteinase K (PK) protected fragment expected upon insertion into the membrane. For OMP25, the endogenous sequence was utilized. However, because of its large size, the RHOT1 TM was instead expressed as a fusion with an N-terminal villin headpiece (VHP) domain (as previously reported). As a MTCH2 independent control, a previously established matrix-targeted control protein was generated by fusing dihydrofolate reductase (DHFR) with residues 1-69 from *N. crassa* ATP synthase subunit 9 (Su9) (*8*).

For expression in insect cells, all MTCH2 structure constructs were cloned into the pFastBac1 backbone with an N-terminal GFP-SUMO^Eu1^ tag for purification (*35*, *67*). Expression constructs cloned into the pHAGE2 lentiviral backbone were used to generate mammalian stable cells lines for expression and purification of MTCH2 samples used in reconstitution experiments (*8*). Constructs for *E. coli* expression of α-Fab and α-UCP1 nanobodies, used for structure determination were cloned into the pET Duet vector (Millipore-Sigma) with an N-terminal pelB leader sequence for periplasmic expression and a 6x His-tag followed by a TEV protease cleavage site. *E. coli* expression of the α-GFP nanobody (Addgene #199370) (*67*) and the SENP^EuB^ protease (*68*) (Addgene #149333) were performed using a previously established plasmid system.

Reporter and rescue constructs contained several endogenous sequences obtained from UniProtKB/Swiss-Prot, including: synaptojanin-2 binding protein (OMP25/SYNJBP; **P57105-1**), import intermembrane translocase subunit Tim9 (TIM9; **Q9VYD7**), mitochondrial antiviral-signaling protein (MAVS; **Q7Z434-1**), FUN14 domain-containing protein 1 (FUNDC1; **Q8IVP5-1**), mitochondrial Rho GTPase 1 (RHOT1; **Q8IXI2-1**), mitochondrial Rho GTPase 2 (RHOT2; **Q8IXI1-1**), mitochondrial import receptor subunit TOM70 (TOMM70; **O94826**), isopentenyl-diphosphate delta-isomerase (*A. thaliana* At5g55610; **Q9FM77**), mitochondrial carrier homolog 1 (MTCH1; **Q9NZJ7-1** [MTCH1-L] and **Q9NZJ7-2** [MTCH1-S], for both long and short isoforms, respectively), mitochondrial outer membrane protein SLC25A46 (SLC25A46; **Q96AG3-1**), mitochondrial carrier protein (*C. owczarzaki* MTCH; **A0A0D2WXZ5**), mitochondrial carrier homolog 2 (*M. musculus* MTCH2; **Q791V5**), LD43650p (*D. melanogaster* MTCH; **Q9V3Y4**), mitochondrial carrier homolog 2 (*N. vectensis* MTCH; **A7SSN1** (associate sequence has been updated, exact sequence used matches XP_032228681 in NIH database), C1q and TNF-related 4 (*X. tropicalis* MTCH2; **Q6P818**), mitochondrial carrier homolog 2 (*H. sapiens* MTCH2; **Q9Y6C9**), Uncharacterized protein (*T. brucei* pATOM36; **Q582I5** [Met 86 chosen as start codon]), Armadillo repeat-containing protein 1 (ARMC1; **Q9NVT9-1**), and Cytochrome b5 type B (CYB5B; **O43169**).

The primary antibodies used for Western Blotting are: MTCH2 (ab113707, Abcam, UK, diluted 1:1000), MTCH1 (PA5-100201, Invitrogen, USA, diluted 1:1000), SLC25A46 (12277-1-AP, Proteintech, USA, diluted 1:2000), ARMC1 (HPA026085, Atlas Antibodies, Sweden, diluted 1:1000), Tubulin (T9026, Sigma-Aldrich, USA, diluted 1:20000), FLAG-HRP (A8592, Millipore-Sigma, USA, diluted 1:10000-1:20000 depending on expression levels of FLAG-tagged construct), and DNAJC11 (17331-1-AP, Proteintech, USA, diluted 1:1000). The secondary antibodies used are: goat α-mouse-HRP (#172-1011, Bio-Rad, USA, diluted 1:5000), and goat α-rabbit-HRP (#170-6515, Bio-Rad, USA, diluted 1:5000).

### Cell culture and cell lines

Human K562 containing the dCas9-KRAB machinery cells were cultured in RPMI-1640 media supplemented with 25 mM HEPES, 2.0 g/L NaHCO_3_, 10% FBS, 2 mM glutamine, 100 U/mL penicillin, and 100 µg/mL streptomycin. Cells were maintained at a density between 0.25 × 10^6^ – 1 × 10^6^ cells/mL. HEK293T cells were cultured in DMEM supplemented with 10% FBS, 2 mM glutamine, 100 U/mL penicillin and 100µg/mL streptomycin. Human cell lines were grown at 37 °C with 5% CO_2_.

All K562 experiments were performed using a CRISPRi cell line with stable expression of dCas9-BFP-KRAB^KOX1^ (*69*) and the IMS-localized MICU1(1-60)-GFP(1-10) (*8*, *21*). To generate this cell line, CRISPRi K562 cells were transduced with lentivirus expressing IMS GFP(1-10) followed by single cell sorting into a 96-well plate using a Sony Cell Sorter (SH800S). Clones expressing IMS GFP(1-10) were identified by transducing with MICU(1-60)-GFP11 and checking for GFP fluorescence. CRISPR-Cas9-derived MTCH2 KO cells were generated on top of this parental cell line, as previously described (*8*), by nucleofecting cells with two MTCH2 targeting sgRNAs (AGCCGACATGTCTCTAGTGG and GGCTTTGCGAGTCTGAACGT) which were cloned in the pX458 plasmid (Addgene #48138) with the Lonza SF Cell Line 96-well Nucleofector Kit (V4SC-2096).

ExpiSf9 cells were used to express MTCH2 for structure determination. Cells were cultured in ExpiSf CD media (ThermoFisher) and maintained between 1 × 10^6^ – 10 × 10^6^ cells/mL at 27 °C with 0% CO_2_ in an orbital shaker at 125 rpm. Lentivirus-generated Expi293F stable cell lines were used to express WT and mutant MTCH2 for reconstitution experiments. Cells were cultured in Expi293 media (ThermoFisher) and maintained between 0.5 × 10^6^ – 2 × 10^6^ at 37 °C with 8% CO_2_ in an orbital shaker at 125 rpm.

### Lentivirus production and spinfection

To produce lentivirus, HEK293T cells were simultaneously transfected with the two packaging plasmids (psPAX2 and pMD2.G were both gifts from Didier Trono, Addgene #12260 and #12259) and the transfer plasmid using the TransIT-293 transfection reagent (Mirus). Supernatant was collected 48 hours after transfection and aliquoted, frozen, and stored at −80° C for later use.

For generating lentivirus with any MTCH1-encoding plasmids, the pan-caspase inhibitor Q-VD-OPh (MedChemExpress) was added to a final concentration of 25 µM at the time of transfection and 24 after transfection to enable lentivirus generation by preventing apoptosis.

### Bioinformatic sequence analysis

MTCH homolog sequences were identified by querying the UniProt knowledgebase (UniProtKB) for sequences annotated with the PANTHER family PTHR10780 (*70*). In order to study the distribution of MTCH homolog subfamilies across the holozoan clade, 10 holozoan species were used in species-restricted UniProtKB queries for sequences annotated with one of three PTHR10780 subfamilies: SF20 (MTCH2), SF3 (MTCH1), and SF18 (invertebrate MTCH). Query results are summarized in Table S4. To visualize the resulting information, query results were mapped onto a published holozoan species tree (Fig. 1A) (*71*).

In order to identify conserved and diverged features between MTCH and canonical SLC25 family members, we aligned individual SLC25 repeats from multiple human SLC25 family members and MTCH homologs with the hmmalign tool from the HMMER3.3 package (*72*) using a publicly available SLC25 hmm profile (pf00153) (fig. S1B) (*73*).

A putative plant OM insertase was identified from a dataset of 42 proteins confirmed to localize to the *A. thaliana* OM using isolation of mitochondria and proteomics (*39*). AlphaFold-predicted models of each protein (*23*, *24*) were visually inspected for features similar to our MTCH2 structure, leading to the identification of At5g55610 (Fig. 5B).

In order to study the distribution of multiple OM insertase families across eukaryotes, 13 eukaryotic lineages (defined by taxid) were used in lineage-restricted UniprotKB queries for sequences annotated with the panther families PTHR10780 (MTCH), PTHR36074 (At5g55610), or PTHR28241 (Mim1) (*70*), or the pfam family PF19224 (pATOM36) (*73*). Alignments of ATOM36 homologs were produced using MUSCLE (*74*) and visualized using the ESPript 3.2 3.2 server (*75*). Query results are summarized in Table S5. To visualize the resulting information, query results were mapped onto a published eukaryotic species tree (Fig. 5A) (*76*).

To assess the quality of AlphaFold models (*23*, *24*), the following parameters were analyzed: the predicted template modeling score (pTM), the predicted local distance difference test (pLDDT), and the predicted aligned error (PAE), and are listed in table S6. Each of these values are included as output from the AlphaFold3 server (*23*). pLDDT was analyzed by mapping values for each residue onto structural models. Additionally, the average pLDDT and PAE of all atoms was also determined.

### Fluorescent reporter experiments in human cells

#### Flow cytometry data collection and analysis

For all fluorescent reporter experiments, data was collected with an Attune NXT Flow Cytometer (ThermoFisher) and analyzed in Python using the FlowCytometryTools package (v0.5.1). FSC-A vs. SSC-A gates were applied to select live cells and FSC-A vs. FSC-H gates were applied to exclude doublets. Appropriate gates selecting BFP-positive cells (for rescue or CRISPRi experiments) and RFP-positive cells (for all reporter experiments) were determined based on measurements of non-fluorescent control cells with either the VL-1 (BFP) or YL-2 (RFP) channels. The GFP:RFP ratio was calculated from the BL-1 (GFP) and YL-2 (RFP) channels to quantify reporter activity. Histograms displaying rescue phenotypes in MTCH2 KO cells throughout the manuscript have been normalized to the median GFP:RFP ratio obtained from cells expressing WT MTCH2 (GFP:RFP_rescue_/GFP:RFP_wt_). Histograms displaying knockdown effects throughout the manuscript have been normalized to values for cells expressing a non-targeting (NT) sgRNA control (GFP:RFP_knockdown_ /GFP:RFP_NT_).

#### CRISPRi knockdown experiments

To assess the effects of depletion of specific factors on OM protein integration, K562 CRISPRi IMS GFP(1–10) cells were spinfected with lentivirus expressing the desired sgRNA (targeting either MTCH2, ARMC1, DNAJC11 or a dual guide targeting MTCH1 as described in Table S2) and a puromycin resistance marker. Cells were then treated with puromycin (1 ug/mL) at 48, 72, and 96-hour timepoints after transduction to select for sgRNA-expressing cells. Five days following the initial transduction, cells were spinfected again with lentiviral constructs expressing the RFP-P2A-GFP11-tagged reporters. For rescue experiments using MTCH1 and MTCH2 (Fig. S2A), BFP-P2A-tagged rescue constructs, and a BFP-only control, were transduced along with the reporter at an MOI of ∼1. Analysis by flow cytometry took place 72 hours after the second spinfection. For all experiments, live cells expressing both BFP and RFP were analyzed for their GFP:RFP expression. We have previously established that decreases in GFP (relative to our expression control) are a proxy for successful mitochondrial integration (*8*).

#### Homolog rescue experiments

K562 CRISPRi IMS GFP(1-10) MTCH2 KO cells were spinfected with two individual lentiviral constructs expressing an RFP-P2A-GFP11-tagged reporter and a BFP-P2A-tagged MTCH homolog (*M. musculus*, *X. tropicalis*, *D. melanogaster, N. vectensis*, and *C. owczarzaki*), pATOM36, or At5g55610 rescue construct (*8*). Additional samples with WT and MTCH2 KO cells were transduced with a BFP-only control. Analysis by flow cytometry took place after 72 hours. The GFP:RFP ratio was analyzed for BFP- and RFP-positive cells after normalizing to the median GFP:RFP ratio obtained from cells expressing WT MTCH2 (GFP:RFP_homolog_/GFP:RFP_wt_).

#### MTCH2 mutational analysis experiments

K562 CRISPRi IMS GFP(1-10) MTCH2 KO cells were spinfected with two individual lentiviral constructs expressing RFP-P2A-GFP11-tagged reporters and BFP-P2A-tagged human MTCH2 WT and mutant rescue constructs. Additional samples with WT and MTCH2 KO cells were transduced with a BFP-only control. Analysis by flow cytometry took place after 72 hours. GFP:RFP was then calculated in BFP-positive and RFP-positive cells to determine whether each MTCH2 mutant could restore OM integration of each reporter construct. This was assessed by normalizing to the median GFP:RFP ratio obtained from cells expressing WT MTCH2 (GFP:RFP_mutant_/GFP:RFP_wt_). To analyze the relationship between hydrophobicity and mutational effects, a hydrophobicity scale (*77*) was used to calculate the hydrophobicity shift in the amino acid at each position (Δhydrophobicity) caused by each mutation. Specific values used were 100 for Phe, 97 for Leu, 76 for Val, 74 for Met, 63 for Tyr, 41 for Ala, −5 for Ser, −10 for Ala, −14 for Arg, −23 for Lys, −28 for Asn, and −31 for E. Δhydrophobicity was calculated by the following formula: hydrophobicity_aa mut_-hydrophobicity_aa wt_, where hydrophobicity_aa mut_ is the hydrophobicity of the mutated amino acid and hydrophobicity_aa wt_ is the hydrophobicity of the WT amino acid at this position. To assess correlations, the split-GFP reporter mutant phenotypes were plotted against Δhydrophobicity, a linear equation was fit to the resulting data points in python using numpy (*78*), and the coefficient of determination (R^2^) was calculated with the following formula: 1 – SS_res_/SS_tot_ where SS_res_ is the residual sum-of-squares (the squared difference of each GFP:RFP value from the linear fit) and SS_tot_ is the total sum-of-square (the squared difference of each GFP:RFP value from the mean of all GFP:RFP values) (Fig. 3D and S14A,B).

#### OM topology experiments

K562 CRISPRi IMS GFP(1-10) cells were spinfected with lentiviral constructs expressing RFP-P2A-GFP11 tagged proteins. Analysis by flow cytometry took place after 72 hours. GFP:RFP was then calculated in RFP-positive cells to determine if the GFP11 tagged termini of each protein was localized in the IMS (fig. S8B-C, S22A).

### *In vitro* mitochondrial insertion

#### Mitochondrial Isolation

To directly test if *C. owczarzaki* MTCH was active at the level of insertion, we generated human K562 cells depleted of MTCH2 but expressing either the human or *C. owczarzaki* MTCH. More specifically, we generated CRISPRi IMS GFP(1-10) MTCH2 KO cells expressing BFP-P2A tagged human 3xFLAG-MTCH2 or *C. owczarzaki* 3xFLAG-MTCH rescue constructs by lentiviral transduction at an MOI of ∼1. WT cells and MTCH2 KO cells expressing a BFP-only control were also generated. ∼20 million BFP expressing cells for each of the four samples were isolated using a BD Biosciences FACS Aria Fusion Cell Sorter, and expanded for 5-7 days prior to harvesting for use in mitochondrial isolation experiments.

Mitochondria were isolated similarly to a previously described protocol (*8*, *79*, *80*). K562 cells were centrifuged for 5 minutes at 220xg. The resulting pellets were then washed in homogenization buffer (210 mM mannitol, 70 mM sucrose, 5 mM HEPES pH 7.4, 10 mM EDTA, 1 mM PMSF, 2 mg/mL BSA), resuspended in 2 mL of the same buffer, and incubated on ice for 10 minutes. A 2 mL glass Dounce homogenizer with a tight-fitting pestle was used to lyse cells using ∼30 passes. After lysis, the cell lysate was centrifuged at 1300x g for 5 minutes. The supernatant was transferred to a clean tube and centrifuged twice more to remove all nuclei and unlysed cells. The supernatant was then transferred to a new tube and centrifuged at 11,000x g for 10 minutes to pellet mitochondria. Mitochondria were washed 2x in isolation buffer (210 mM mannitol, 70 mM sucrose, 5 mM HEPES pH 7.4, 10 mM EDTA) and the final pellet was resuspended in 5-10µl isolation buffer. Prior to downstream use, protein concentration was measured using a Bradford assay and each mitochondrial sample was normalized by diluting to a protein concentration of 5 mg/mL.

#### In vitro translation and insertion

*In vitro* translation was performed using rabbit reticulocyte lysate (RRL) (*66*) and mitochondrial insertion was assessed as previously described (*8*, *79*). Transcription templates were amplified via PCR using primers which bound upstream of the SP6 promoter and ≥ 12 bp downstream of the stop codon, and purified using the QIAquick PCR purification kit (Qiagen). The DNA template was then transcribed using SP6 polymerase at 37 °C for 90 minutes. mRNA was then translated for 15-30 minutes at 32 °C in the presence of radioactive ^35^S-methionine, followed by addition of 1 mM puromycin to end translation and ensure nascent chain release from the ribosome.

Insertion reactions with isolated mitochondria were performed by diluting 4µl translation into 50µl import buffer (250 mM sucrose, 5 mM Mg(Acetate)_2_, 80 mM KAcetate, 20 mM HEPES pH 7.4, 2.5 mM ATP, and 15 mM succinate) and adding 15 µg purified mitochondria, followed by incubation at 32 °C for 30 minutes.

To determine insertion efficiency, proteinase K (PK) was added at a concentration of 0.25 mg/mL followed by incubation on ice for 1 hour. Reactions were quenched by addition of 5 mM PMSF in DMSO, followed by transfer to boiling 1% SDS in 0.1 mM Tris/HCl pH 8.0. 6xHis-tagged protected fragments were affinity purified by diluting PK-digested samples in IP buffer (50 mM HEPES pH 7.5, 500 mM NaCl, 10 mM imidazole, 1% Triton X-100) and incubating with 10 µL NiNTA resin while mixing for 1.5 hours at 4 °C. The resin was then washed 3x with 1 mL IP buffer, and the enriched fragments were then eluted from resin in Laemmli Sample buffer supplemented with 50 mM EDTA pH 8.0. PK protected fragments were quantified after running samples on SDS-PAGE gels and imaging with autoradioagraphy.

#### Proteoliposome reconstitutions and insertion reactions

MTCH2-containing proteoliposomes were prepared as previously described (*8*, *61*, *79*). Egg-derived phosphatidyl-choline (PC), egg-derived phosphatidylethanolamine (PE), and synthetic 1,2-dioleoyl-sn-glycero-3-phosphoethanolamine-N-lissamine rhodamine B (Rh-PE) were obtained from Avanti Polar Lipids. Rh-PE was included to standardize lipid quantities across conditions after reconstitution. Chloroform stocks of each lipid were combined at a mass ratio of 8:1.9:0.1 PC:PE:Rh-PE, DTT was added to 10 mM, and the mixture was then evaporated under vacuum overnight. The dried lipids were then rehydrated in lipid buffer (15% glycerol, 50 mM HEPES pH7.4) by mixing for 8 hours at 25 °C and then diluted to a final concentration of 20 mg/mL. BioBeads-SM2 (BioRad) were activated with methanol and then washed thoroughly in water before being resuspended in a 50% slurry. Reconstitution mixtures were prepared by mixing 30-55µl purified MTCH2 and 8µl 20 mg/mL lipids and brought up to a final volume of 80µl with additional components added to achieve a final buffer condition of 100 mM NaCl, 25 mM HEPES pH 7.4, 2 mM MgCl_2_, 0.8% DBC. Empty liposomes were produced in the same manner except the purification buffer was added in place of MTCH2. This protein/lipid/detergent mixture was incubated on ice for 10 minutes and then added to 80µl packed biobeads with all water removed in a 1.5 mL eppendorf tube, and then mixed for 18 hours at 4 °C in a thermomixer. Next, the liquid was removed, diluted into 500µl ice-cold water, and pelleted by ultracentrifugation in a TLA100.3 rotor at 70,000 rpm for 30 minutes. After removal of the supernatant, the resulting pellet was resuspended in 14µl liposome resuspension buffer (100 mM KAcetate, 50 mM HEPES pH 7.4,, 2 mM Mg(Acetate)_2_, 250 mM sucrose, 1 mM DTT). Proteoliposomes were normalized by measuring the fluorescence intensity of Rh-PE in a BioTek Synergy HTX plate reader (Agilent) and diluting with additional liposome resuspension buffer to achieve matched concentrations. OMP25-6xHis was used as a substrate. After a 20 minute translation in RRL in the presence of ^35^S-methionine, the reaction was quenched with 1 mM puromycin to inhibit synthesis and trigger ribosomal release. This translation reaction was then separated into multiple samples for post-translational insertion into proteoliposomes. Insertion reactions were prepared by mixing 1µl proteoliposome resuspension with 9µl translation and incubating at 32°C for 10 minutes. Next, reactions were treated with PK at 0.5 mg/mL on ice for 1 hour and quenched by adding PMSF and boiling in SDS. His-tagged protected fragments were purified with NiNTA as described above. Samples were analyzed by on SDS-PAGE and imaged via autoradiography to quantify substrate insertion. Insertion experiments were performed with a dilution series to determine conditions with matching MTCH2 levels. To quantify the relative amount of wildtype and mutant MTCH2 in each reaction, a dilution series of wildtype MTCH2 was measured by Western Blot. Using this dilution series, a linear regression was performed and applied to quantify the relative amount of each MTCH2 mutant utilized in each insertion reaction. The % inserted for each wildtype and mutant MTCH2 was calculated relative to the maximum insertion observed for wildtype MTCH2 at its highest concentration (i.e. 100*[intensity of protease protected fragment MTCH2 mutant/intensity of protease protected fragment of wildtype MTCH2 at highest concentration]).

### Structural determination of human MTCH2

#### Design of human MTCH2 expression constructs for structure

Due to its small size (33 kDa), structure determination of MTCH2 required fiducial markers to enable the accurate determination of particle orientation and to allow 3D reconstruction. We utilized two strategies to do this. First, we tested fusion to the small globular domain, BRIL, which itself can be recognized by a semi-synthetic Fab (BAK5) (*28*) which was further stabilized by an additional α-Fab nanobody (Nb) (*31*), resulting in a trimeric complex of ∼100 kDa. While partially successful, we observed significant flexibility between the BRIL fusion and MTCH2 in these reconstructions that limited the resulting resolution.

In addition, we reasoned we could leverage the tools developed for determination of the recent structure of the SLC25, UCP1 (*29*, *81*). In this case, an α-UCP1 Nb (pMb65) was developed that specifically recognized the loop between TMs 4 and 5 of UCP1, which is a similar length in all SLC25s and MTCH family homologs. We therefore created a MTCH2 fusion with the short sequence of UCP1 that is recognized by this α-UCP1 Nb. The α-UCP1 Nb was engineered to enable the binding of a universal α-nanobody Fab which was further stabilized through the binding of an additional α-Fab Nb, resulting in a tetrameric complex of ∼110 kDa.

#### BRIL fusion construct

To design the MTCH2-BRIL fusion MTCH2(BRIL), human MTCH2 residues 207-214 were replaced with an engineered variant of apocytochrome b562 (BRIL) flanked by two α-helical linkers, in order to enable complexing with synthetic antibodies to facilitate cryo-EM structure determination (*28*). Rosetta (*82*, *83*) was used to identify specific linker lengths predicted to form rigid α-helices, and optimal sequences were designed from the resulting backbone using ProteinMPNN (*84*). The designs were then further optimized by iterative modification based on AlphaFold predictions (*23*, *24*).

#### UCP1 fusion construct

To design the MTCH2-UCP1 fusion, MTCH2(UCP1), human MTCH2 residues 203-222 were replaced with UCP1 residues 198-218, and an additional 3 stabilizing mutations were included: L199V, A200Y, T226A to recreate an α-UCP1 Nb (pMb65) binding epitope (*29*).

To enable the formation of a larger complex containing MTCH2 (UCP1), the TC-Nb4 interface was engineered onto pMb65 to allow the binding of the nanobody-binding-Fab (NabFab) (*30*) by removing the first three residues (GPS) and introducing 6 point mutations: V5R, A17P, P27A, K89E, Q123K, and Q126P.

#### MTCH2 UCP1 mutant constructs

To determine structures of representative members of the Class I and Class II hyperactive MTCH2 mutants, we introduced mutations into the MTCH2 UCP1 fusion construct. For the Class I F285N F286N mutant, we used blunt end ligation with phosphorylated primers in which the mutations were designed into the 5’ end of one primer. For the Class II K25E Y235A V23D triple mutant, we used Gibson assembly in which the mutations were designed into the primer overhangs.

#### Expression construct design

The resulting MTCH2 sequences were cloned into a baculovirus expression plasmid pFastBac1 to enable expression in Sf9 cells. The MTCH2 sequence was fused to a GFP-SUMO^Eu1^ tag to enable native purification using the GFP Nb strategy previously described by our lab in which GFP Nb is immobilized to affinity resin (in this case NiNTA), and MTCH2 is eluted by cleavage of the SUMO^Eu1^ by incubation the engineered protease SENP^EuB^.

#### Expression and purification of SENP^EuB^ protease

To allow the native elution of untagged MTCH2 following isolation using the GFP Nb, we used SENP^EuB^ protease, because we had previously shown this results in better yield and cleaner preparations than other strategies. Expression and purification of the SENP^EuB^ protease (*67*) was conducted based on a previously described protocol, modified with an additional size exclusion step to reduce contaminants interfering with structure determination efforts. In brief, the plasmid was transformed into NEBExpress *E. coli* expression strain (New England Biolabs). A single colony was inoculated into 50 mL Luria-Bertani (LB) medium supplemented with 50 µg/mL kanamycin and incubated shaking at 37 °C for 4-5 hours. 5 mL of overnight culture was inoculated into 1 L SuperBroth (SB) medium supplemented with 50 µg/mL kanamycin and incubated at 37 °C while shaking at 220 rpm. Expression was induced by addition of 1mM IPTG when OD_600_ = 0.8. Cells were grown for 6 hours at 18 °C post induction and harvested by centrifugation. Cells were resuspending in 50 mL resuspension buffer (50 mM Tris/HCl pH 7.5, 500 mM NaCl, 10 mM imidazole, and 5 mM beta-mercaptoethanol) and flash frozen for later use.

The resulting resuspension was thawed and supplemented with cOmplete^TM^ EDTA-free Protease Inhibitor Cocktail (Roche), lysed by sonication, and cleared by centrifugation at 18,000 rpm and 4 °C for 30 minutes using an SS-34 rotor (Beckman-Coulter). Cleared lysate was incubated with 1 mL Ni-NTA agarose (Qiagen) for 15 minutes while rolling at 4 °C. Ni-NTA resin was then loaded onto a gravity flow column (Bio-Rad) and washed with 100 mL resuspension buffer. Protein was eluted with 5 mL of elution buffer (50 mM Tris pH 7.5, 150 mM NaCl, 500 mM Imidazole, and 5 mM beta-mercaptoethanol). Eluted protein was then buffer exchanged into storage buffer (50 mM Tris pH 7.5, 150 mM NaCl, 10 mM Imidazole, 10% [v/v] glycerol, and 5 mM beta-mercaptoethanol) using PD-10 desalting column (Cytiva). Protein was then cleaved overnight with TEV protease. After cleavage, protein was flowed over NiNTA resin to remove any remaining un-cleaved protein. The cleaved flowthrough was then concentrated using a 10K MWCO concentrator (Millipore-Sigma) and injected onto a S200 increase 10/300 GL size exclusion column (Cytiva) equilibrated in SEC buffer (150 mM NaCl, 50 mM Tris pH 7.5, 10% glycerol [v/v], and 1 mM DTT). The protein-containing peak was pooled, concentrated, aliquoted, and then flash frozen in liquid nitrogen and stored for later use.

#### Expression and purification of α-GFP Nb

We leveraged the extremely specific and high affinity interaction between the GFP Nb and GFP-MTCH2 to allow isolation in a single step from Sf9 cells. Expression and purification of the α-GFP Nb was conducted as previously described. Briefly, the plasmid was transformed into AVB101 *E. coli* expression strain (New England Biolabs). A single colony was inoculated into 100 mL LB medium supplemented with 50 µg/mL kanamycin plus 10 µg/mL carbenicillin and incubated at 37 °C while shaking at 220 rpm overnight. 5 mL overnight culture was inoculated into 1 L SB medium supplemented with 50 µg/mL kanamycin, 10 µg/mL carbenicillin, and 50 µM biotin, and grown to an OD_600_ = 1 up while shaking at 37 °C. Expression was induced by addition of 1 mM IPTG. Cells were grown overnight at 18 °C post induction and harvested by centrifugation. Cells were resuspending in 50 mL wash buffer (50 mM Tris/HCl pH 7.5, 500 mM NaCl, and 10 mM imidazole) and flash frozen for later use.

After thawing, the resuspension was supplemented with cOmplete^TM^ EDTA-free Protease Inhibitor Cocktail (Roche), subsequently lysed by sonication, and cleared by centrifugation at 18,000 rpm and 4 °C for 30 minutes using an SS-34 rotor (Beckman-Coulter). Cleared lysate was incubated with 1 mL Ni-NTA agarose (Qiagen) for 15 minutes while rolling at 4 °C. Ni-NTA resin was then loaded onto a gravity flow column (Bio-Rad) and washed with 100 mL or wash buffer. Protein was eluted with 5 mL of elution buffer (50 mM Tris pH 7.5, 150 mM NaCl, and 500 mM Imidazole). Eluted protein was then buffer exchanged into storage buffer (50 mM Tris pH 7.5, 150 mM NaCl, 10 mM Imidazole, and 10% [v/v] glycerol) using a PD-10 desalting column (Cytiva), aliquoted, and flash frozen for later use.

#### Expression and purification of α-UCP1 and α-Fab Nbs

To increase the size of MTCH2 for structure determination, we used purified α-UCP1 and α-Fab Nbs to form the MTCH2(BRIL)•BAK5•α-Fab Nb and MTCH2 (UCP1)•α-UCP1 Nb•NabFab•α-Fab Nb complexes with the modified MTCH2 sequences described above. The α-UCP1 Nb is based on a previously characterized nanobody (*29*) further engineered to allow binding to NabFab (*30*), while the α-Fab Nb (*31*) aids in structure determination by binding to the hinge region of the NabFab and BAK5 synthetic antibodies in order to increase overall complex size, break symmetry of the Fab, and reduce flexibility. To express the α-UCP1 and α-Fab Nbs, plasmids were transformed into BL21 (DE3) *E. coli* expression strain. A single colony was inoculated into 50 mL Luria-Bertani medium supplemented with 100 µg/mL carbenicillin and incubated at 37 °C while shaking at 220 rpm overnight. 5 mL of overnight culture was inoculated into 1 L SB medium supplemented with 100 µg/mL carbenicillin. Expression was induced by addition of 1mM IPTG when OD_600_ = 0.8. Cells were grown for 4 hours after induction and harvested by centrifugation. Pelleted cells were resuspended in 50 mL wash buffer (50 mM Tris pH 7.5, 200 mM NaCl, 10 mM Imidazole) and flash frozen for later use.

After thawing, the resuspension was supplemented with cOmplete^TM^ EDTA-free Protease Inhibitor Cocktail (Roche), subsequently lysed by sonication, and cleared by centrifugation at 18,000 rpm and 4 °C for 30 minutes using an SS-34 rotor (Beckman-Coulter). Cleared lysate was incubated with 1 mL Ni-NTA agarose (Qiagen) for 15 minutes while rolling at 4 °C. Ni-NTA resin was then loaded onto a gravity flow column (Bio-Rad) and washed with 100 mL of wash buffer. Protein was eluted with 5 mL of elution buffer (50 mM Tris pH 7.5, 150 mM NaCl, and 500 mM Imidazole). Eluted protein was then buffer exchanged into storage buffer (50 mM Tris pH 7.5, 150 mM NaCl, 10 mM Imidazole, and 10% [v/v] glycerol) using PD-10 desalting column (Cytiva). Protein was then cleaved overnight with TEV protease. After cleavage, protein was flowed over NiNTA resin to remove any remaining uncleaved protein. The cleaved flowthrough was then concentrated using a 10K MWCO concentrator (Millipore-Sigma) and injected onto a S200 increase 10/300 GL size exclusion column (Cytiva) equilibrated in SEC buffer (150 mM NaCl, 50 mM Tris pH 7.5, 10% glycerol [v/v]). The protein-containing peak was pooled, concentrated, aliquoted, and then flash frozen in liquid nitrogen and stored at −80 °C for later use.

#### Expression and purification of synthetic antibodies NabFab and BAK5

To further increase the size of MTCH2 for structure determination, we used the synthetic antibodies BAK5 and NabFab to form the MTCH2(BRIL)•BAK5•α-Fab Nb and MTCH2 (UCP1)•α-UCP1 Nb•NabFab•α-Fab Nb complexes with the modified MTCH2 sequences described above. BAK5 binds to a BRIL domain that we engineered into the loop in between MTCH2 TM4 and TM5, while NabFab binds to an interface that was engineered into the α-UCP1 Nb sequence. The pRH2.2 NabFab plasmid was transformed into C43(Pro+) *E. coli* expression strain (both plasmid and expression strain provided as a kind gift by A. Kossiakoff) (*30*). The pRH2.2 α-BRIL Fab BAK5 was transformed into BL21-Gold *E. coli* expression strain (Agilent). All remaining steps were the same for both constructs. A single colony was used to inoculate a culture in Luria Broth (LB) medium supplemented with 100 µg/mL carbenicillin for overnight incubation while shaking at 37 °C 220 rpm. 20mL of overnight culture was subsequently inoculated into 2 L SB medium supplemented with 100 µg/mL Carbenicillin. Expression was induced by addition of 1mM IPTG when OD_600_ = 0.8. Cells were grown for 4 hours at 37 °C after induction and harvested by centrifugation. Pelleted cells were flash frozen for storage at −80 °C.

The resulting pellet was resuspended in lysis buffer (50 mM Tris pH 7.5, 200 mM NaCl, and 1x cOmplete^TM^ EDTA-free Protease Inhibitor Cocktail [Roche]). Cells were lysed by sonication. The lysate was incubated at 62 °C for 30 minutes and cleared by centrifugation at 18,000 rpm and 4 °C for 30 minutes using a SS-34 rotor (Beckman-Coulter). Cleared lysate was subsequently loaded onto a 5 mL Pierce™ Protein L column (ThermoScientific) and washed with 50 mL of wash buffer (50 mM Tris pH 7.5 and 500 mM NaCl). Protein was eluted with 100 mM acetic acid and immediately neutralized by 1M Tris. Protein-containing fractions were loaded onto a 1 mL HiTrap SP cation-exchange column (Cytiva) and washed with 5 mL of cation exchange wash buffer (50 mM Na Acetate pH 5.0). The protein was eluted by running a 0-100% gradient over 25 mL of cation exchange elution buffer (50mM Na Acetate pH 5.0 and 2M NaCl) followed by immediate neutralization using 1 M Tris. Purified protein was buffer exchanged into storage buffer (50 mM Tris pH 7.5, 200mM NaCl, and 5% glycerol) using a PD-10 desalting column (Cytiva). Buffer exchanged protein was concentrated using a 30K MWCO concentrator (Millipore-Sigma), then flash frozen in liquid nitrogen and stored at −80 °C for later use.

#### Baculovirus-based expression of MTCH2 in insect cells for structure determination

To obtain large quantities of the complex-forming MTCH2 variants described above (BRIL and UCP1), sequences were fused to an N-terminal GFP-SUMO^Eu1^ tag (GFP-SUMO^Eu1^-MTCH2) and cloned into the pFastbac1 backbone to enable expression in insect cells with baculovirus. The plasmids were transformed into MAX Efficiency^TM^ DH10Bac competent cells (Gibco) and the PureLink^TM^ HiPure plasmid miniprep kit (Invitrogen) was used to obtain bacmid DNA. Bacmid DNA was then transfected in ExpiSf9 cells using the ExpiFectamine™ Sf transfection reagent (Gibco) and cell viability was monitored over several days. After cell viability dropped below 80%, cells were pelleted by centrifugation at 300x rcf for 5 minutes and the baculovirus containing supernatant (P0) was stored at 4 °C protected from light for later use. ExpiSF cells were mixed with a dilution series of baculovirus and cultured for 24 hours, and the then the % GFP-positive cells was measured using the Attune NxT flow cytometer (ThermoFisher) with a BL1 channel gate generated using uninfected ExpiSF cells. Next, a larger culture of ExpiSF cells was mixed with P0 baculovirus at a multiplicity of infection (MOI) of 0.075 and cell viability was monitored for several days. After cell viability dropped below 80%, cells were pelleted by centrifugation at 300x g for 5 minutes and the baculovirus containing supernatant (P1) was stored at 4 °C protected from light for later use.

For protein expression, ExpiSf9 cells were seeded at a density of 2.5 x 10^6^ cells/mL and ExpiSf Enhancer (Gibco) was added. On the next day, P1 baculovirus was added at a MOI of 2. The cells were harvested after 48 hours by centrifugation at 6,000 rpm and 4 °C for 10 minutes in a H-6000A rotor (Sorvall). The resulting pellet was washed by resuspending in phosphate-buffered saline and centrifuging again at 3500x rcf and 4 °C for 10 minutes. The supernatant was then discarded and the pellet was weighed, then flash frozen in liquid nitrogen and stored at −80 °C for later use.

#### Purification of MTCH2 (UCP1) and (BRIL) for structure determination

GFP-SUMO^Eu1^-MTCH2 (UCP1 or BRIL) was purified using an α-GFP Nb as previously described (*67*) with some modifications. To isolate a mitochondrially-enriched membrane fraction, cells were resuspended in hypotonic lysis buffer (10mM HEPES/KOH pH 7.5, 10 mM K Acetate, 1.5 mM Mg Acetate, 2 mg/mL Iodoacetamide, and 1x cOmplete^TM^ EDTA-free Protease Inhibitor Cocktail [Roche]) and incubated on ice for 10 minutes followed by lysis using a 40 mL glass Dounce homogenizer with a loose-fitting pestle, followed by addition of NaCl to 190 mM. Membranes were pelleted by centrifugation for 10 minutes at 18,000 rpm and 4 °C using an SS-34 rotor (Beckman-Coulter). The pelleted membranes were washed 1x with membrane wash buffer (10 mM HEPES/KOH pH 7.5, 200 mM NaCl, 1.5 mM Mg Acetate) and resuspended at a total protein concentration of 2.5 mg/mL in solubilization buffer (50 mM HEPES/KOH pH7.5, 200 mM NaCl, 2 mM Mg Acetate, 50 mM imidazole, 2 mg/mL Iodoacetamide, 1x cOmplete^TM^ EDTA-free Protease Inhibitor Cocktail [Roche], and 2% [w/v] n-Undecyl-β-D-Maltoside [UDM; Anatrace]) and incubated for 1 hour at 4 °C while rolling. Debris was removed by centrifugation for 30 minutes at 18,000 rpm and 4 °C using an SS-34 rotor.

Molar concentration of GFP-SUMO^Eu1^-MTCH2 in the solubilized membrane fraction was estimated by measuring fluorescence with a purified GFP standard in a BioTek Synergy HTX plate reader (Agilent). 14x His-tagged GFP Nb was then added to the solubilized membrane fraction at a 2-fold molar excess to GFP-SUMO^Eu1^-MTCH2 and incubated with Ni-NTA agarose resin (QIAGEN, 30210) at 4 °C for 1 hour while rolling. Ni-NTA resin was then washed two times in wash buffer A (50 mM HEPES/KOH pH 7.5, 200 mM NaCl, 2 mM Mg Acetate, 50 mM imidazole, 1x cOmplete^TM^ EDTA-free Protease Inhibitor Cocktail [Roche], 0.5 mM ATP, and 0.1% UDM), with a 15 minute incubation per wash step. Lastly, Ni-NTA resin was washed twice with wash buffer B (50 mM HEPES/KOH pH 7.5, 200 mM NaCl, 2 mM Mg Acetate, 50 mM imidazole, 1x cOmplete^TM^ EDTA-free Protease Inhibitor Cocktail [Roche], and 0.07% UDM). Protein was eluted by incubating with wash buffer B + 600 nM SENP^EuB^ protease for 1 hour at 4 °C while rolling. Eluted protein was concentrated to 500uL using 30K MWCO concentrator (Millipore-Sigma) and injected onto a Superdex 200 Increase 10/300 GL column (Cytiva) equilibrated in size exclusion buffer (50 mM HEPES/KOH pH 7.5, 200 mM NaCl, 2 mM Mg Acetate, and 0.07% UDM). MTCH2-containing fractions were then pooled prior to downstream complexing steps.

For MTCH2^F285N,F286N^ and MTCH2^K25E,Y235A,V238D^, the same purification strategy was used with the following differences: after washing twice with buffer A, an additional wash step was performed with wash buffer A2 (50 mM HEPES/KOH pH 7.5, 200 mM NaCl, 10 mM K Acetate, 2 mM Mg Acetate, 50 mM imidazole, 1x cOmpleteTM EDTA-free Protease Inhibitor Cocktail [Roche], and 1% Triton X-100 [v/v]).

#### Purification of WT and mutant MTCH2 for reconstitution experiments

WT and mutant MTCH2 used in reconstitution experiments were purified from detergent solubilized mammalian cells using a biotinylated anti-GFP nanobody as previously described (*67*). MTCH2 expressing stable cell lines were induced with 1 µg/mL doxycycline and cells were grown for 48 more hours before harvesting by centrifugation at 6,000 rpm and 4 °C for 10 minutes in a H-6000A rotor (Sorvall) (*67*). The resulting pellet was then resuspended in phosphate-buffered saline and centrifuged at 3,500x rcf and 4 °C for 10 minutes. After discarding supernatant, pellets were weighed, flash frozen in liquid nitrogen, and stored for later use.

Cell pellets were resuspended at a 1:10 (m/v) ratio of cell pellet to hypotonic lysis buffer (10 mM HEPES/KOH pH 7.5, 10 mM K(Acetate), 0.15 mM Mg(Acetate)_2_, 0.5 mM DTT, 1x cOmpleteTM EDTA-free Protease Inhibitor Cocktail [Roche]). After incubating on ice for 10 minutes, cells were lysed with 10x strokes in a Dounce homogenizer with a loose-fitting pestle. NaCl was then added to 190 mM and cell membranes were pelleted by centrifugation at 18,000x rcf at 4 °C for 10 minutes in a SS-34 rotor (Beckman-Coulter). After discarding supernatant, cell membranes were resuspended in membrane wash buffer (10 mM HEPES/KOH pH 7.4, 200 mM NaCl, 0.15 mM Mg(Acetate)_2_, 0.5 mM DTT) and pelleted once more by centrifugation at 18,000x rcf at 4 °C for 10 minutes in a SS-34 rotor. The resulting pellet was then solubilized by resuspending in solubilization buffer (50 mM HEPES/KOH pH 7.5, 200 mM NaCl, 2 mM Mg(Acetate)_2_, 1% deoxy-BigCHAP [DBC; Millipore-Sigma], 1 mM DTT, 1x cOmpleteTM EDTA-free Protease Inhibitor Cocktail [Roche]) at a ratio of 6.8 mL buffer per 1g original cell pellet. After incubating at 4 °C while rolling for 30 minutes, the insoluble material was pelleted by centrifugation at 18,000x rcf and 4 °C for 30 minutes in a SS-34 rotor (Beckman-Coulter). Next, the supernatant was added to magnetic Streptavidin resin (Thermo Fisher) which were bound to biotinylated α-GFP Nb, blocked with free biotin, and pre-equilibrated. After binding for 1 hour while rolling at 4 °C, the ATP was added to 0.5 mM, the input material was removed, and the resin was then washed four times with 1 mL wash buffer A (solubilization buffer with 0.2% DBC and 0.5 mM ATP). Next the resin was washed with 1 mL wash buffer B (wash buffer A with no ATP) and transferred to a clean tube. To elute protein, resin was resuspended in wash buffer + 600 nM SENP^EuB^ while rolling head-over-tail for 2h at 4 °C.

#### MTCH2 complex assembly for structure determination

*MTCH2 (BRIL)•BAK5•α-Fab Nb complex formation |* Size-exclusion purified MTCH2 (BRIL) was mixed with BAK5 and α-Fab Nb at a 1:1.5:2 molar ratio followed by incubation at 4 °C overnight. The mixture was then concentrated to 100 µL and injected onto a Superdex 200 Increase 10/300 GL column (Cytiva) equilibrated in size exclusion buffer (fig. S4B). Fractions containing the MTCH2 (BRIL)•BAK5•α-Fab Nb complex were pooled and concentrated using a 100K MWCO concentrator (Millipore-Sigma).

*MTCH2 (UCP1)•α-UCP1 Nb•NabFab•α-Fab Nb complex formation* | Size-exclusion purified MTCH2 (UCP1) was mixed with α-UCP1 Nb, NabFab, and α-Fab Nb at a 1:1.5:2:2.5 molar ratio and incubated overnight at 4 °C. The mixture was concentrated to 500 µL and injected onto a Superdex 200 Increase 10/300 GL column (Cytiva) equilibrated in size exclusion buffer (fig. S6B). Fractions containing the MTCH2 (UCP1)•α-UCP1 Nb•NabFab•α-Fab Nb complex were pooled and concentrated using a 100K MWCO concentrator (Millipore-Sigma).

#### Cryo-EM grid preparation and data collection

To prepare cryo-EM grids, 3.1 µL of either MTCH2 (BRIL)•BAK5•α-Fab Nb complex or MTCH2 (UCP1)•α-UCP1 Nb•NabFab•α-Fab Nb complex was applied to Quantifoil R 1.2/1.3 holey carbon film grids (Ted Pella, Inc.) which had been glow-discharged with a PELCO easiGlow^TM^ (Ted Pella, Inc.) at 20 mA for 60 s. A FEI Vitrobot Mark IV (Thermo Fisher) was used to blot grids with filter paper and then plunge freeze them in liquid ethane. Blotting was carried out at 6 °C and 100% humidity with −1 blot force for 5s. The protein concentration used for grid preparation was 3.5 mg/mL for the MTCH2 (BRIL)•BAK5•α-Fab Nb complex, 5.9 mg/mL for the WT MTCH2 (UCP1)•α-UCP1 Nb•NabFab•α-Fab Nb complex, 3.4 mg/mL for the MTCH2^F285N,F286N^ (UCP1)•α-UCP1 Nb•NabFab•α-Fab Nb complex, and 4.9 mg/mL for the MTCH2^K25E,Y235A,V238D^ (UCP1)•α-UCP1 Nb•NabFab•α-Fab Nb complex.

Data were collected on a Titan Krios electron microscope (Thermo Fisher) at 300 keV equipped with a K3 direct electron detector (Gatan) and a 20 eV slit width energy filter. Automatic image acquisition was performed using SerialEM in super resolution mode at a magnification of 105,000, corresponding to a calibrated pixel size of 0.418 Å/pixel. Images were collected at a defocus range of −0.8 to −1.8 µm and a total dose of 60 e^−^/Å^2^. We collected for a total of 26,889 micrographs for the MTCH2 (BRIL)•BAK5•α-Fab Nb complex, 15,334 micrographs for the WT MTCH2 (UCP1)•α-UCP1 Nb•NabFab•α-Fab Nb complex, 10,506 micrographs for the MTCH2^F285N,F286N^ (UCP1)•α-UCP1 Nb•NabFab•α-Fab Nb complex, and 10,816 micrographs for the MTCH2^K25E,Y235A,V238D^ (UCP1)•α-UCP1 Nb•NabFab•α-Fab Nb complex.

#### Cryo-EM image processing

Graphical schematics of the processing workflow for each structure are shown in fig. S4, S6, S16, and S18. All data processing was performed with CryoSPARC v4.7.0 (*85–87*). Movies were downsampled to 0.836 Å/pixel, motion-corrected, dose-weighted, and the contract transfer function (CTF) was estimated in CryoSPARC Live.

For MTCH2 (BRIL)•BAK5•α-Fab Nb, 23,738 out of 26,889 micrographs were selected for downstream processing based on CTF quality, ice thickness, and visual inspection. The blob picker was used on 6,937 micrographs to pick 4,622,038 particles which were extracted at 4.16 Å/pixel. After iterative rounds of 2D classification, classes containing 50,000 particles were selected for ab-initio reconstruction to generate 3 volumes. One of the 3 volumes resembled a Fab attached to a detergent solubilized membrane protein and was used to generate templates for template picker. Resulting particle picks were combined with the initial particle set for a total of 7,786,491 particles. Together, 11 rounds of heterogenous refinement were conducted using one ab-initio volume corresponding to the MTCH2 complex and four decoy volumes, resulting in a final particle set containing 199,397 particles, which were then extracted at 0.998 Å/pixel. A final map was obtained from non-uniform refinement. Local resolution was calculated using a mask from the non-uniform refinement job and visualized in ChimeraX. Settings of the data collection are reported on Table S1.

For MTCH2 (UCP1)•α-UCP1 Nb•NabFab•α-Fab Nb, 14,190 of out 15,334 micrographs were selected for downstream processing based on CTF quality, ice thickness, and visual inspection. The blob picker was used on 4,885 micrographs to pick 1,082,415 particles which were extracted at 3.44 Å/pixel. After iterative rounds of 2D classification, classes containing 160,865 particles were selected for to ab-initio reconstruction to generate 3 volumes. One of the 3 volumes resembled a Fab attached to a detergent solubilized membrane protein and was used to generate templates for template picker. Resulting particle picks were combined with the initial particle set for a total of 4,791,546 particles. Together, 11 rounds of heterogenous refinement were conducted using one ab-initio volume corresponding to the MTCH2 complex and four decoy volumes, resulting in a final particle set containing 485,622 particles, which were then extracted at 0.83 Å/pixel. The extracted particles were processed to correct for anisotropic magnification using the global CTF refinement job and the motion-correction and dose-weighting was refined using reference-based motion correction. After this processing, a final map was obtained from non-uniform refinement followed by local refinement with mask on MTCH2 only. Local resolution was calculated using a mask for MTCH2 and visualized in ChimeraX. B-factor was calculated and visualized in ChimeraX. Settings of the data collection are reported on Table S1.

For MTCH2^F285N,F286N^ (UCP1)•α-UCP1 Nb•NabFab•α-Fab Nb, the same processing steps were performed with the following differences: 6,937 out of 10,506 micrographs were selected for downstream processing. A total of 2,013,310 particles from blob picker and template picker were obtained. After running 5 rounds of heterogenous refinement, 321,419 particles were selected for further processing.

For MTCH2^K25E,Y235A,V238D^ (UCP1)•α-UCP1 Nb•NabFab•α-Fab Nb, the same processing steps were performed with the following differences: 9,633 out of 10,816 micrographs were selected for downstream processing. A total of 3,547,162 particles from blob picker and template picker were obtained. After running 7 rounds of heterogenous refinement, 612,877 particles were selected for further processing.

#### Model building and refinement

For MTCH2 (BRIL)•BAK5•α-Fab Nb, an AlphaFold predicted model of MTCH2 (BRIL) (*23*, *24*) was docked into the EM density using ChimeraX (*88*). There was prominent density for the four-helix bundle of BRIL with one rigid helix linking the C-terminus of BRIL with TM5 of MTCH2. Low resolution density could be observed for the remainder of MTCH2, roughly confirming the positions of TM3 and TM4, but the rest of the density was too poor to be useful for modelling.

To build MTCH2 (UCP1)•α-UCP1 Nb, the initial model for MTCH2 (UCP1) was generated with AlphaFold (*23*, *24*) using the sequence corresponding to the MTCH2 (UCP1), and the initial model for the α-UCP1 Nb was derived from a previously determined structure (PDB ID 8G8W) (*29*). The initial models were loaded into ChimeraX (*88*) and docked into the EM density, then the docked models were loaded into COOT (*89*) and the TC-Nb4 scaffold residues which enable NabFab binding were manually mutated. The models were manually rebuilt using COOT, including adjustment to the C-terminal region (285–301) which was poorly predicted by AlphaFold. The model was then subsequently subjected to iterative refinement with *phenix.real_space_refinement* (*90*) using secondary restraints and Ramachandran restrains. Statistics of the EM map and model refinement are reported in Table S1.

The complete model for the WT MTCH2^UCP1^•α-UCP1 Nb complex was docked into the maps for MTCH2^F285N,F286N^ (UCP1)•α-UCP1 Nb•NabFab•α-Fab Nb and MTCH2^K25E,Y235A,V238D^ (UCP1)•α-UCP1 Nb•NabFab•α-Fab Nb and subject to further manual building in COOT, including a significant conformational change of residues 283-286 in the MTCH2^K25E,Y235A,V238D^ model, and further rounds of iterative refinement with *phenix.real_space_refinement*. Since no density was observed for the loop in between TM2 and TM3 (residues 100-115) this region was excluded from the final model.

#### Structural model analysis

Molecular analysis and graphic were performed utilizing USCF ChimeraX (*88*) and PyMOL (*91*). Coulombic electrostatic potential was calculated and mapped onto the surface representation of each structure in ChimeraX. Previously determined experimental structures of Ylp1, YidC, the endoplasmic reticulum (ER) membrane protein complex (EMC), uncoupling protein 1 (UCP1), and the ATP/ADP translocase were used in comparisons with our MTCH2 structure (*25*, *26*), while models of MTCH homologs, *T. brucei* pATOM36, and *A. thaliana* At5g55610 were generated using AlphaFold (*23*, *24*). In surface representations of MTCH2 and other models, a black line was drawn to outline the hydrophilic crevice, which we define as a concave hydrophilic surfaces flanked by hydrophobic TM residues.

## Acknowledgements

We thank F. Y. Wang, Z. Fan, and M. Luo for assistance with protein purification. We thank the Caltech cryo-EM facility, S. Chen and T. Brittain for support for cryo-EM data collection. We thank C. Gati, P. Dutka, Y. Li, and G. P. Tomaleri for cryo-EM data processing advice. We thank A. Kossiakoff and S. Mukherjee for providing reagents necessary to generate synthetic antibodies. We thank the Caltech protein expression center for insect cell expression assistance.

## Funding

This work was supported by funding from: the NIH’s National Institute of General Medicine 1K99GM151478 (A.G.), the Dr. Nagendranath Reddy Graduate Fellowship (T.A.S.), the Hevolution/AFAR new investigator award (R.M.V.), a Caltech Center for Evolutionary Science Grant (R.M.V./T.A.S.), the Caltech OTCCP Rothenberg Innovation Initiative grant (T.A.S.), the Merkin Institute Translational Research Grant (T.A.S.), and the Sontag Foundation (R.M.V.). R.M.V. is a Howard Hughes Medical Institute Freeman Hrabowski Scholar.

## Competing interests

R.M.V. is a consultant and equity holder of Gate Bioscience. The authors have no additional competing financial interests.

## Data and materials availability

All data needed to evaluate the conclusions in the paper are present in the paper and/or the Supplementary Materials. Cryo-EM maps are deposited in the PDB with IDs 12OY (WT), 12OZ (F285N F286N), and 12PB (K25E Y235A V238D), and in the EMDB with IDs EMDB-76655 (WT), EMDB-76656 (F285N F286N), EMDB-76659 (K25E Y235A V238D), and EMDB-76658 (MTCH2-BRIL).

## Supplementary Materials

Figs. S1 to S24

Tables S1 to S6

References (*94*)

Movies S1

## Notes

### Summary of Updates

Additional data shown in Fig. 3, Fig. 4, Fig. S1, Fig. S7, Fig. S10, Fig. S15, Fig. S20, Fig. S24, and Table S6; Introduction, results, and discussion sections revised; Methods section moved from supplement to main text.

