## Supplementary Materials for "Structural evolution of the MTCH family of mitochondrial insertases"

**The PDF file include**s:

Figs. S1 to S24
Tables S1 to S6
References *(94)*

**Other Supplementary Materials for this manuscript include the following:**

Movies S1

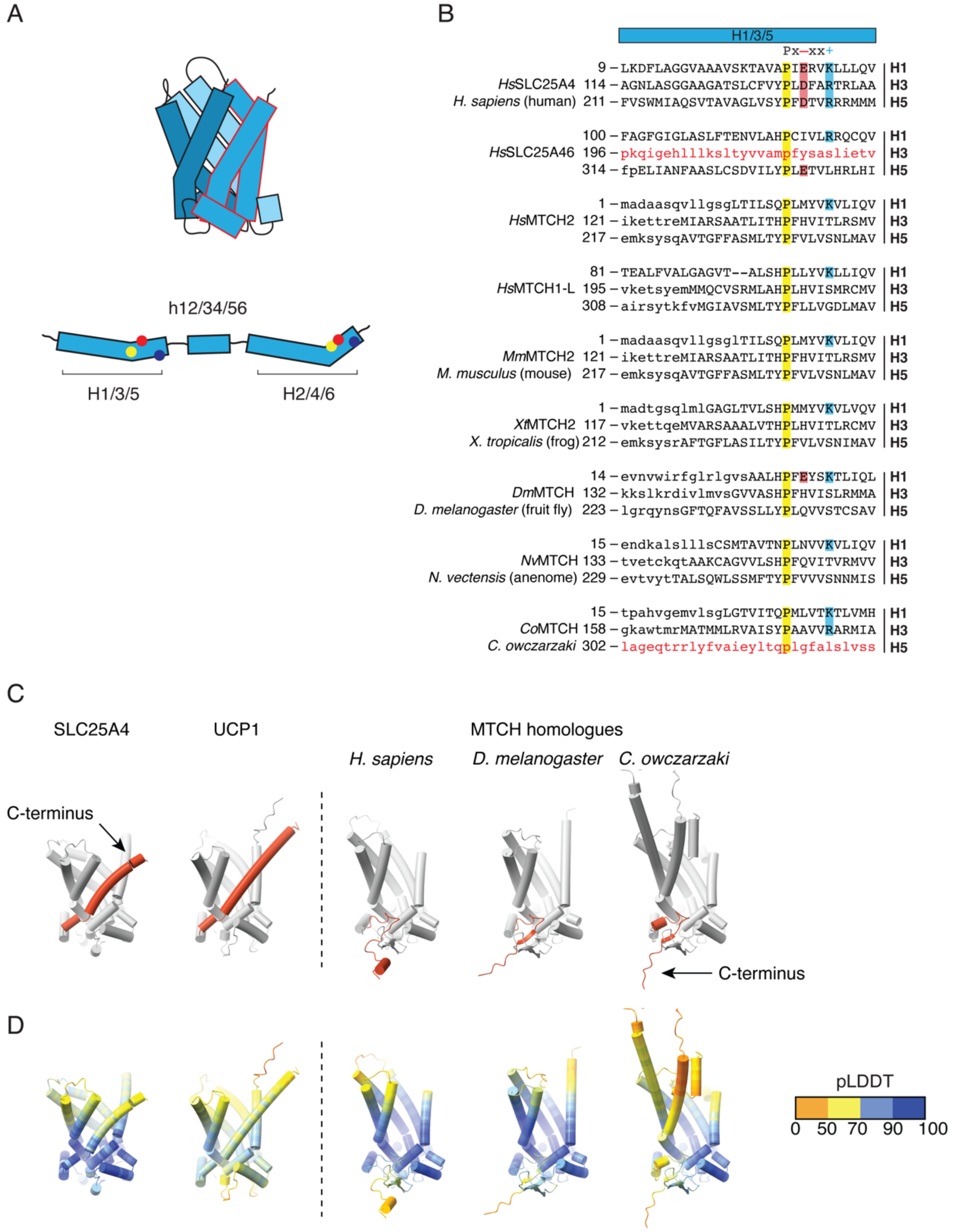


**Fig. S1. Sequence analysis of conserved features across the MTCH insertase family. A)** Schematic of the canonical SLC25 carrier fold highlighting the three pairs of TMs (six total) and their features that are conserved in SLC25 transporters. **B)** Hmmalign based sequence alignment of H1/3/5 of a canonical SLC25 transporter (the inner membrane localized SLC25A4), the OM localized SLC25A46, and 7 MTCH homologs. MTCH1 sequence numbering is based on the long isoform (MTCH1-L). Unaligned sections are shown in lowercase with no gaps. Highlighted columns correspond by color to the features shown in (A). The universally conserved proline, found in both the SLC25 and MTCH family is highlighted in yellow. The two charged residues shown in red and blue form a salt bridge that is important for stabilizing the cytosol-open conformation of canonical SLC25 carriers. The two repeats shown in red text were not recognized by hmmalign and were positioned based on the location of the conserved proline. **C)** Comparison between AlphaFold predicted models of the canonical SLC25 carriers, SLC25A4 and UCP1 (from *H. sapiens*), and the indicated MTCH homologs from *H. sapiens* (MTCH2), *D. melanogaster*, and *C. owczarzaki*. In each model the C-terminal domain (MTCH homologs) or sixth TM (SLC25s) is highlighted in red to highlight its divergence. **D)** The same models as in (C) colored by pLDDT. AlphaFold model confidence values are reported in table S6.

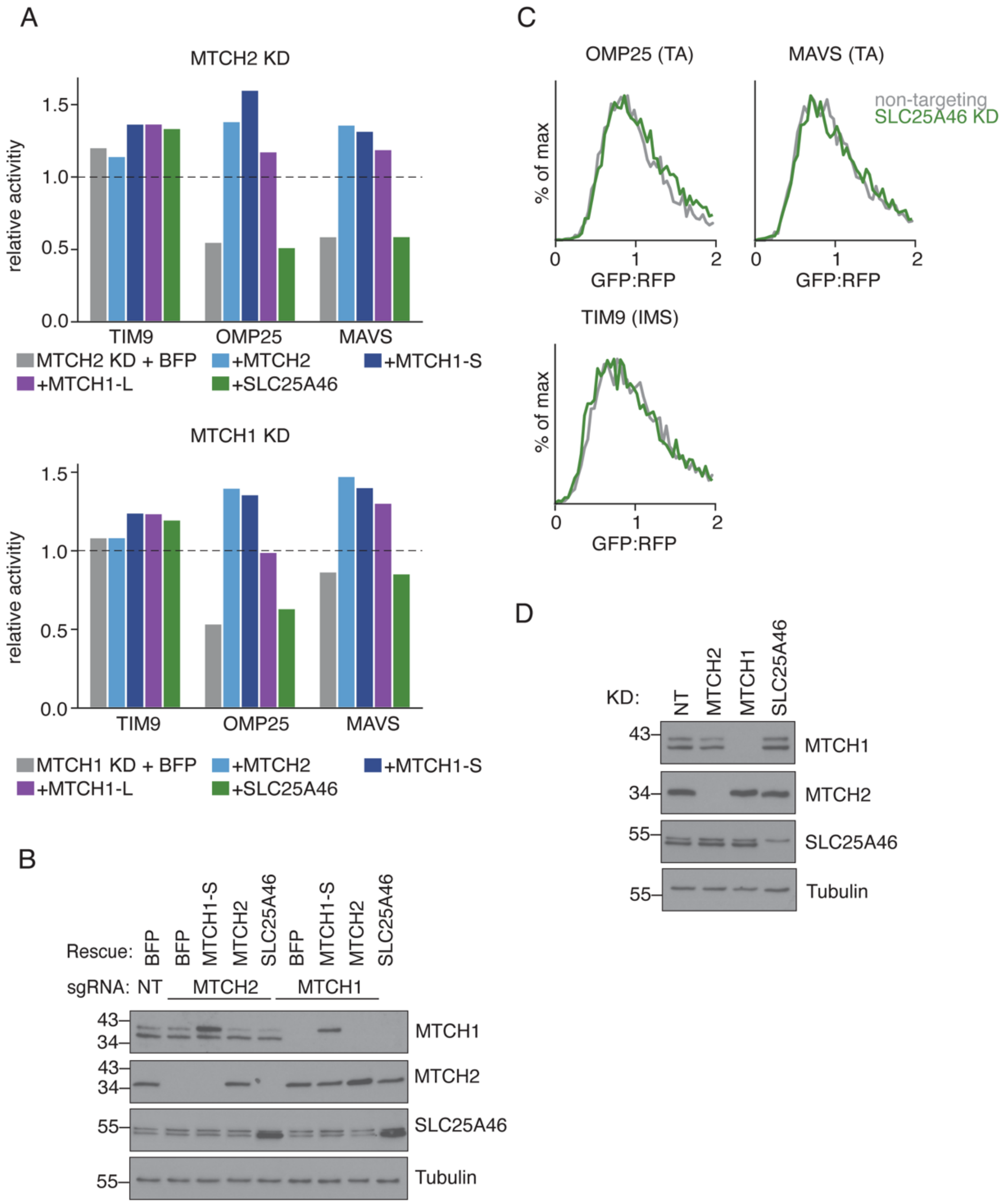


**Fig. S2. OM insertion is a unique feature of MTCH1 and MTCH2 among human SLC25 carriers. A)** (Top) To test whether the three SLC25 family members localized to the OM in human cells, MTCH1, MTCH2, and SLC25A46, are capable of mediating α-helical membrane protein insertion, we leveraged the ratiometric fluorescent reporter system described in Fig. 1B. Using human K562 CRISPRi cells constitutively expressing GFP(1-10) in the mitochondrial IMS, we depleted MTCH2 by transducing either a non-targeting (nt) or a MTCH2-targeting sgRNA. Rescue constructs encoding a BFP control, MTCH2-P2A-BFP, MTCH1-P2A-BFP (long [-L] and short [-S] splice isoforms), and SLC25A46-P2A-BFP were expressed to determine if they could rescue the MTCH2 KD insertion phenotype on three GFP-11 fused reporters: the OM proteins OMP25 and MAVS, and a soluble IMS control, TIM9. Relative activity was calculated using the median GFP:RFP values for each rescue condition normalized to the median GFP:RFP values from nt cells (calculated as $GFP{:}RFP_{knockdown}/GFP{:}RFP_{nt}$). (Bottom) The same rescue experiment was performed with K562 CRISPRi cells expressing a MTCH1-targetting sgRNA. MTCH1 and MTCH2 are both capable of rescuing the phenotype when the other is depleted, while SLC25A46 is not. **B)** Immunoblotting was performed to confirm the MTCH1 and MTCH2 knockdown efficiency and the expression of the rescue constructs (MTCH1-S, MTCH2, and SLC25A46) shown in (A) relative to a housekeeping control (tubulin). While both MTCH1 isoforms are expressed in K562, the MTCH1-S isoform appears to be the dominant one. Note that our MTCH1-S rescue construct runs at a slightly higher size due to the presence of a 9-amino acid GS linker at its N-terminus. **C)** As in (A) except testing the effect of depletion of SLC25A46 on OM insertion of the indicated reporter constructs. Consistent with the results in (A) knockdown of SLC25A46 had no effect on OM insertion. **D)** Immunoblotting to show efficient depletion of SLC25A46 for the experiment in (C).

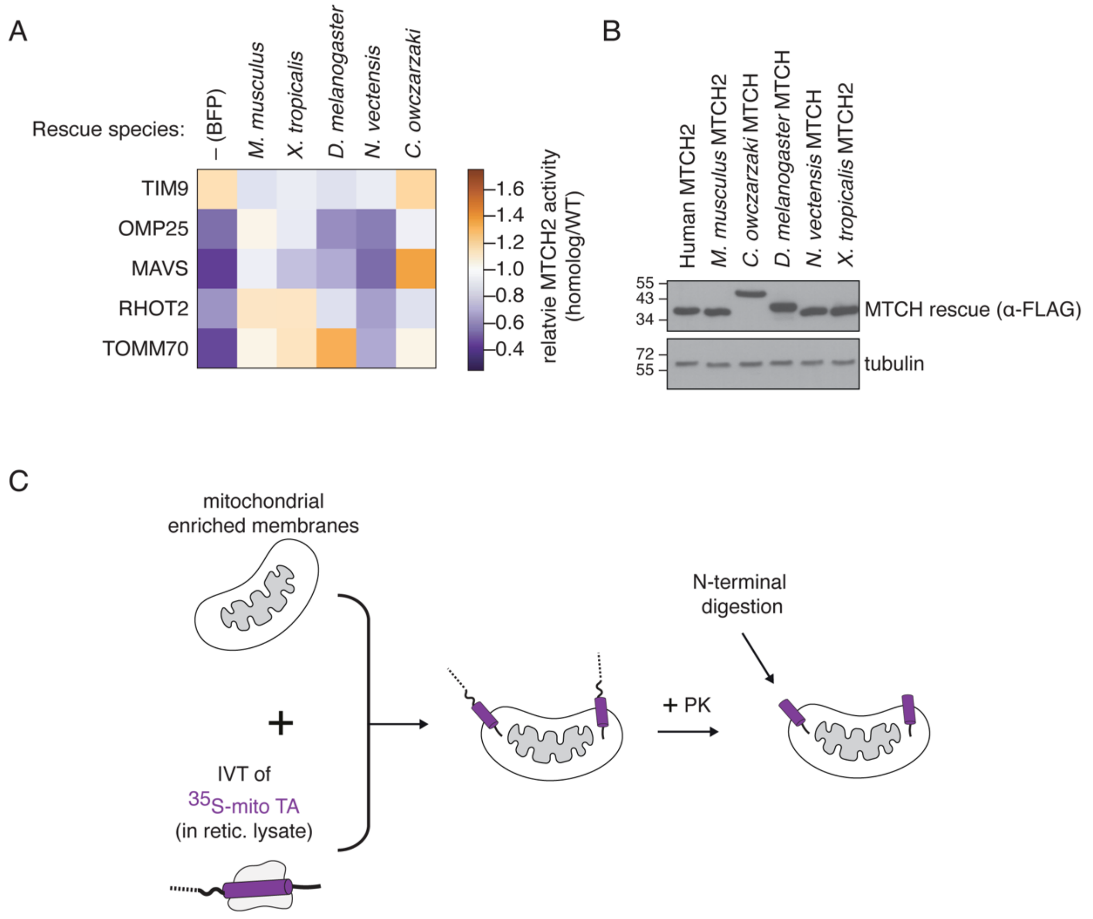


**Fig. S3**. **MTCH homologs are highly conserved across holozoans and all function as OM insertases. A)** The activity of MTCH homologs from *M. musculus, C. owkzarzaki, D. melanogaster*, *N. vectensis*, and *X. tropicalis* were assessed to determine if OM insertion of α-helical proteins is a conserved function across the MTCH family. The ratiometric reporter system was used to measure the insertion of five OM reporter proteins (OMP25, MAVS, RHOT2, FUNDC1, TOMM70) and the soluble IMS control, TIM9, in human K562 MTCH2 KO cells. Rescue constructs encoding either a BFP control, human (*H. sapiens*) MTCH2, or five different MTCH homologs, all 3xFLAG-tagged, were transduced along with the indicated reporters. 48 hours after transduction, reporter insertion was measured using flow cytometry. Displayed is the insertion activity of each homolog relative to the human MTCH2 rescue, calculated as: median $GFP{:}RFP_{homolog}$/median $GFP{:}RFP_{human}$. The resulting values are displayed as a heat map with orange representing increased rescue activity compared to human MTCH2, white representing equal rescue activity, and blue representing reduced or no rescue activity. Representing a remarkable level of conservation, the majority of MTCH homologs, even from distant organisms such as the single-celled *C. owkzarzaki,* are capable of rescuing the MTCH2 knockout phenotype in human cells for several human substrates. Data for select homologs are plotted as histograms in Fig. 1C. **B)** Immunoblotting was performed to confirm expression of the MTCH homologs shown in (A), which was normalized for transduced cells using % BFP positive cells. **C)** Schematic of an in vitro insertion assay into purified mitochondria as previously described (*8*, *79*), used to generate the data displayed in Fig. 1D. An $^{35}$S-methionine -labeled OM tailanchored (TA) substrate is translated in rabbit reticulocyte lysate. Translation is terminated by incubation with puromycin, followed by incubation with mitochondria purified from K562 cells expressing a particular MTCH family member. Insertion of the OM substrate in the correct topology is measured by a protease protection assay, in which mitochondria are incubated with proteinase K (PK), followed by affinity purification of the protease protected fragment using a C-terminal 6x-HIS tag. Insertion is quantified by SDS-PAGE and autoradiography.

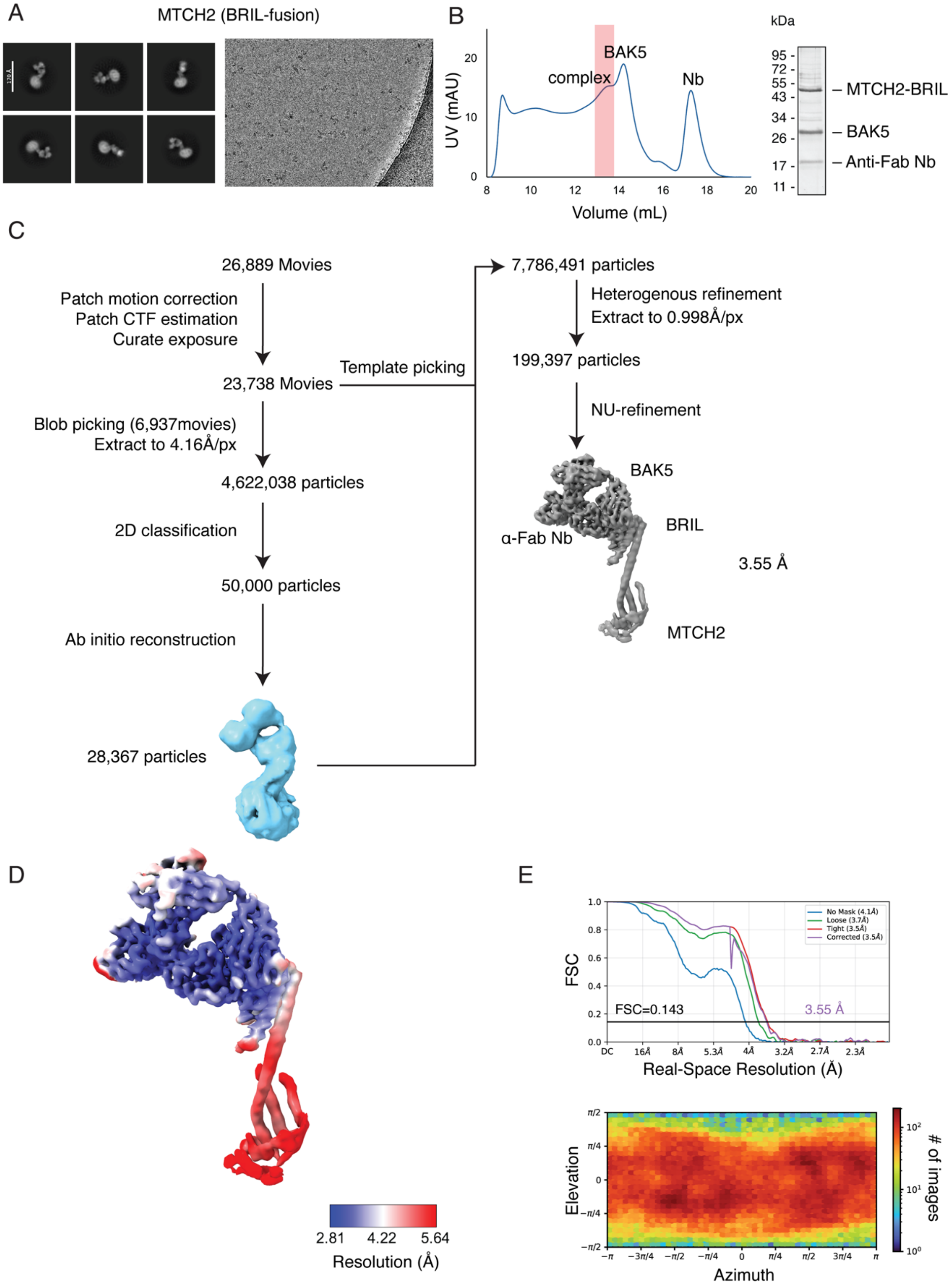
A
MTCH2 (BRIL-fusion)
B
UV (mAU)
complex
BAK5
Nb
Volume (mL)
kDa
– MTCH2-BRIL
– BAK5
– Anti-Fab Nb
C
26,889 Movies
Patch motion correction
Patch CTF estimation
Curate exposure
23,738 Movies
Template picking
Blob picking (6,937movies)
Extract to 4.16Å/px
4,622,038 particles
2D classification
50,000 particles
Ab initio reconstruction
28,367 particles
7,786,491 particles
Heterogenous refinement
Extract to 0.998Å/px
199,397 particles
NU-refinement
BAK5
BRIL
α-Fab Nb
3.55 Å
MTCH2
D
2.81
4.22
5.64
Resolution (Å)
E
FSC
FSC=0.143
3.55 Å
Real-Space Resolution (Å)
Elevation
Azimuth
### of images

**Fig. S4. Cryo-EM data processing workflow for MTCH2(BRIL)•BAK5•α-Fab Nb. A)** (left) Representative 2D class averages of the MTCH2(BRIL)•BAK5•α-Fab Nb complex, (right) and a representative micrograph from the data collection. **B)** (left) Size exclusion chromatography trace with the peak corresponding to the MTCH2 (BRIL)•BAK5•α-Fab Nb complex highlighted in red. (right) Pooled peak fractions were analyzed by SDS-PAGE and SYPRO Ruby stain. **C)** Data processing pipeline using cryoSPARC v4.7.0. Micrographs were curated and a subset was used for blob picking. The resulting particle set was extracted at 4.16 Å/pixel and subject to iterative rounds of 2D classification. Particles from selected 2D classes were used in ab-initio reconstruction to generate 3D volumes. The volume representing a Fab attached to a detergent micelle was selected and used to generate 2D templates for template picker. After re-picking particles from the entire dataset, the resulting particle set was combined with the initial set and processed through iterative rounds of heterogenous refinement, yielding a final particle set of 199,397 particles. The particles were re-extracted at 0.998 Å/pixel and used to generate a final map in a non-uniform refinement job. **D)** Cryo-EM density map of MTCH2 (BRIL)•BAK5•α-Fab Nb colored by local resolution in Å as calculated by cryoSPARC v4.7.0. **E)** Gold Standard Fourier Shell Correlation (GSFSC) curves (top) of the MTCH2 (BRIL)•BAK5•α-Fab Nb complex map with a loose mask (green), tight mask (red), corrected mask (purple) or no mask (blue), and particle Euler angle distribution (bottom). A nominal resolution of 3.55 Å was determined for this map based off of an FSC cutoff of 0.143. Both plots were generated from the local refinement job in cryoSPARC v4.7.0.

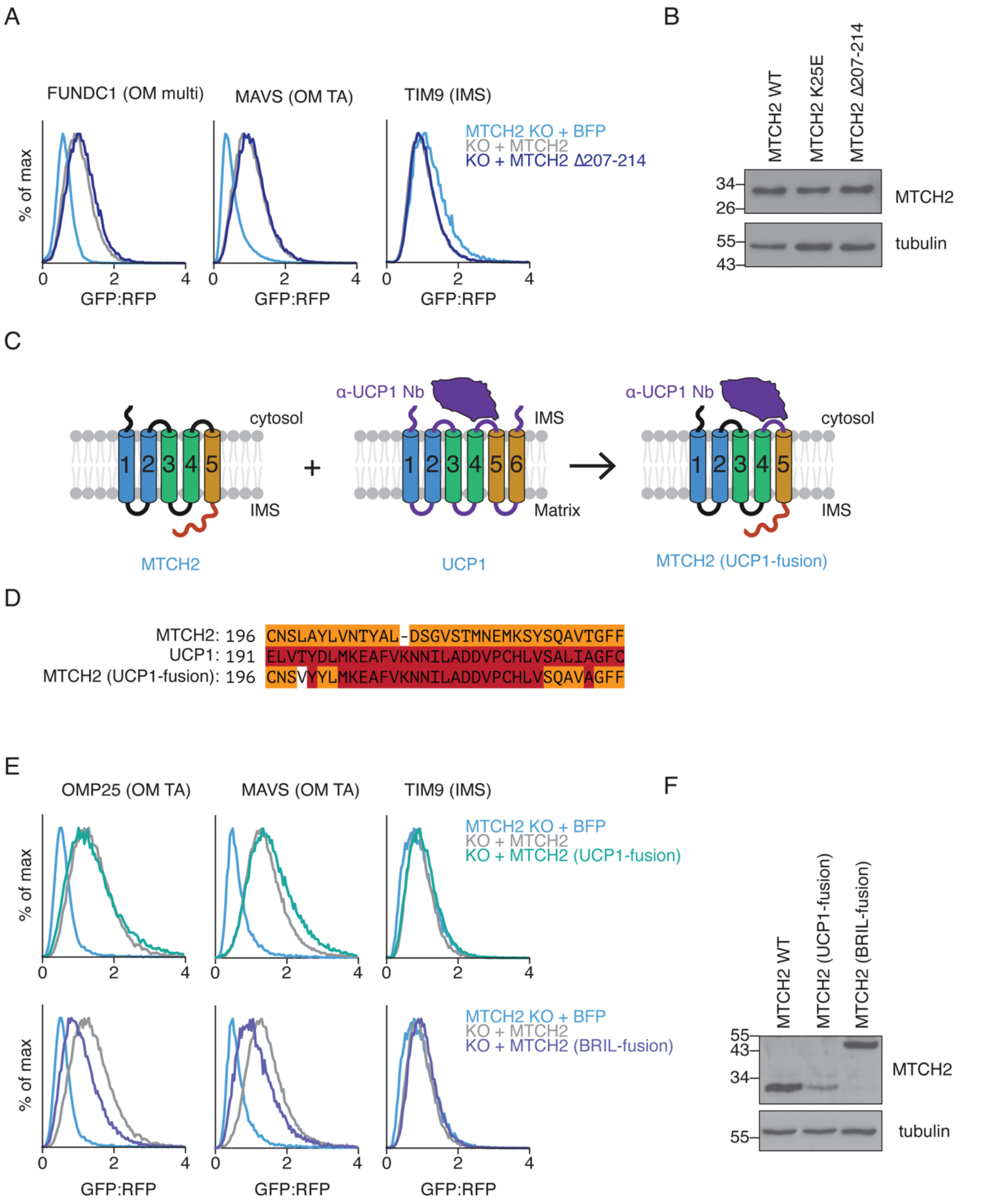


**Fig. S5. A universal strategy for determination of SLC25 structures including the diverged MTCH family homologs. A)** The functional importance of the MTCH2 loop 4/5 was tested using a deletion mutant. The ratiometric reporter system was used to measure the insertion of two OM proteins, FUNDC1 (a multi-pass protein) and MAVS (a TA), and the soluble IMS control TIM9 in MTCH2 KO cells. Rescue constructs encoding either a BFP control, WT MTCH2, or a MTCH2 variant with residues 207-214 in loop 4/5 replaced with a glycine-serine linker of equivalent length were expressed to determine if the loop 4/5 mutant could rescue the

MTCH2 KO phenotype. MTCH2 activity is plotted as a histogram. Loop 4/5 mutants were equally as active as WT MTCH2, suggesting that loop 4/5 is dispensable for its insertase activity. **B)** Immunoblotting analysis of MTCH2 was performed to verify expression of the constructs from (A). To account for differences in transduction efficiency, samples are normalized to % BFP positive cells. Expression of the loop4/5 mutant is displayed relative to that of other MTCH2 mutations that do not cause destabilization. **C)** Schematic of the strategy used to determine the structure of human MTCH2 using an α-UCP1 nanobody. This design strategy exploits the fact that MTCH2 and UCP1 are both SLC25 carriers and thus have a similar fold. A previously characterized α-UCP1 nanobody (pMb65) recognizes a small number of sequential UCP1 residues within the IMS-facing loop between TM4 and TM5 (*29*), a region known to be dispensable for function in other SLC25 carriers (*32*) as well as MTCH2 (A). To generate a MTCH2 variant capable of binding pMb65, several residues in MTCH2 loop 4/5 were replaced with those from UCP1 (shown in purple). **D)** Sequence alignment showing the mutations to WT MTCH2 within cytosolic loop 4/5 made to allow the α-UCP1 antibody to recognize MTCH2. Human MTCH2 residues 203-222 were replaced with UCP1 residues 198-218, and three point mutations, L199V, A200Y, and T226A, were introduced to recreate the α-UCP1 nanobody (pMb65) binding epitope Residues of the MTCH2 (UCP1) originated from human MTCH2 are shown in orange, UCP1 residues in red, and stabilizing mutations are left uncolored. **E)** The activity of the MTCH2 (UCP1) and (BRIL) variants were assessed to determine if the modifications interfered with insertase function. Insertion of two OM proteins, OMP25 and MAVS, and the soluble IMS control TIM9 were tested in human K562 MTCH2 KO cells. Rescue construct encoding either a BFP control, WT MTCH2, MTCH2 (UCP1), or MTCH2 (BRIL) were expressed to determine if the variants designed for structure determination could rescue the MTCH2 KO phenotype. Both MTCH2 fusions mediate insertion of the OM reporters as well as WT suggesting the mutations to enable structure determination do not affect MTCH2 activity in cells**. F)** Immunoblotting analysis of MTCH2 was performed to verify the expression of the constructs from (E). To account for differences in transduction efficiency, samples are normalized to the % BFP positive cells.

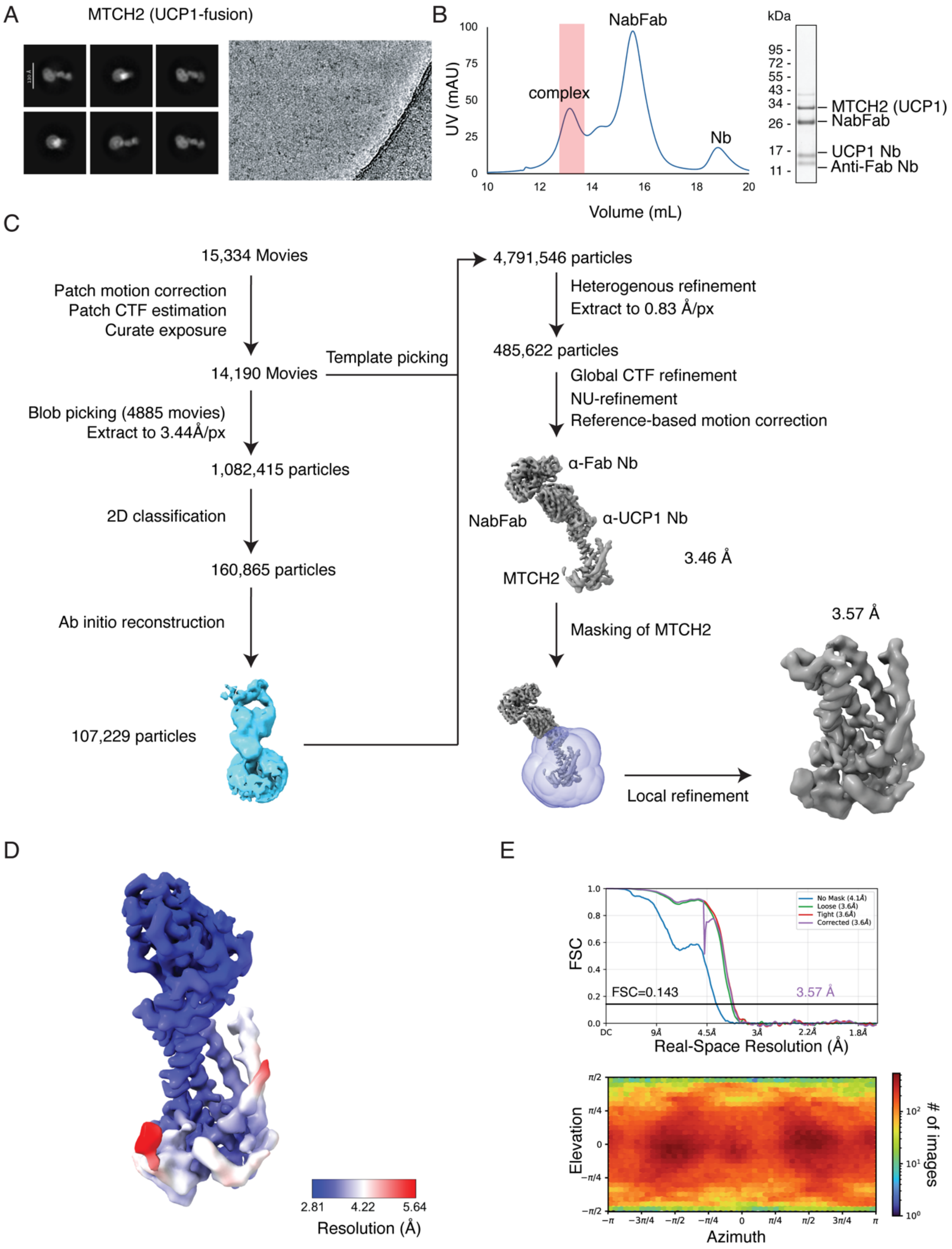
A
MTCH2 (UCP1-fusion)
B
UV (mAU)
Volume (mL)
NabFab
complex
Nb
kDa
95 -
72 -
55 -
43 -
34 -
26 -
17 -
11 -
MTCH2 (UCP1)
NabFab
UCP1 Nb
Anti-Fab Nb
C
15,334 Movies
Patch motion correction
Patch CTF estimation
Curate exposure
14,190 Movies
Template picking
Blob picking (4885 movies)
Extract to 3.44Å/px
1,082,415 particles
2D classification
160,865 particles
Ab initio reconstruction
107,229 particles
4,791,546 particles
Heterogenous refinement
Extract to 0.83 Å/px
485,622 particles
Global CTF refinement
NU-refinement
Reference-based motion correction
α-Fab Nb
NabFab
α-UCP1 Nb
MTCH2
3.46 Å
Masking of MTCH2
3.57 Å
Local refinement
D
2.81
4.22
5.64
Resolution (Å)
E
FSC
No Mask (4.1Å)
Loose (3.6Å)
Tight (3.6Å)
Corrected (3.6Å)
FSC=0.143
3.57 Å
Real-Space Resolution (Å)
Elevation
Azimuth
### of images

**Fig. S6. Cryo-EM data processing workflow for MTCH2•α-UCP1 Nb•NabFab•α-Fab Nb.** **A)** (left) Representative 2D class averages of the MTCH2•α-UCP1 Nb•NabFab•α-Fab Nb tetrameric complex, (right) and a representative micrograph from the data collection. **B)** (left) Size exclusion chromatography trace with the peak corresponding to the MTCH2•α-UCP1 Nb•NabFab•α-Fab Nb complex highlighted in red. (right) Pooled peak fractions were analyzed by SDS-PAGE and SYPRO Ruby stain (right). **C)** Data processing pipeline of MTCH2 using cryoSPARC v4.7.0. Micrographs were curated and a subset was used for blob picking. The resulting particle set was extracted at 3.44 Å/pixel and subject to iterative rounds of 2D classification. Particles from selected 2D classes were used in ab-initio reconstruction to generate 3D volumes. The volume representing a Fab attached to a detergent micelle was selected and used to generate 2D templates for template picker. After re-picking particles from the entire dataset, the resulting particle set was combined with the initial set and processed through iterative rounds of heterogenous refinement, yielding a final particle set of 485,622 particles. The particles were re-extracted at 0.83 Å/pixel and subject to global CTF refinement and reference-based motion correction. The final map was obtained from a non-uniform refinement job followed by the local refinement job using a mask on MTCH2 only. **D)** Cryo-EM density map of MTCH2•α-UCP1 Nb colored by local resolution in Å as calculated by cryoSPARC v4.7.0. **E)** Gold Standard Fourier Shell Correlation (GSFSC) curves (top) of the MTCH2•α-UCP1 Nb map with a loose mask (green), tight mask (red), corrected mask (purple) or no mask (blue), and particle Euler angle distribution (bottom). A nominal resolution of 3.57 Å was determined for this map based off of an FSC cutoff of 0.143. Both plots were generated from the local refinement job in cryoSPARC v4.7.0.

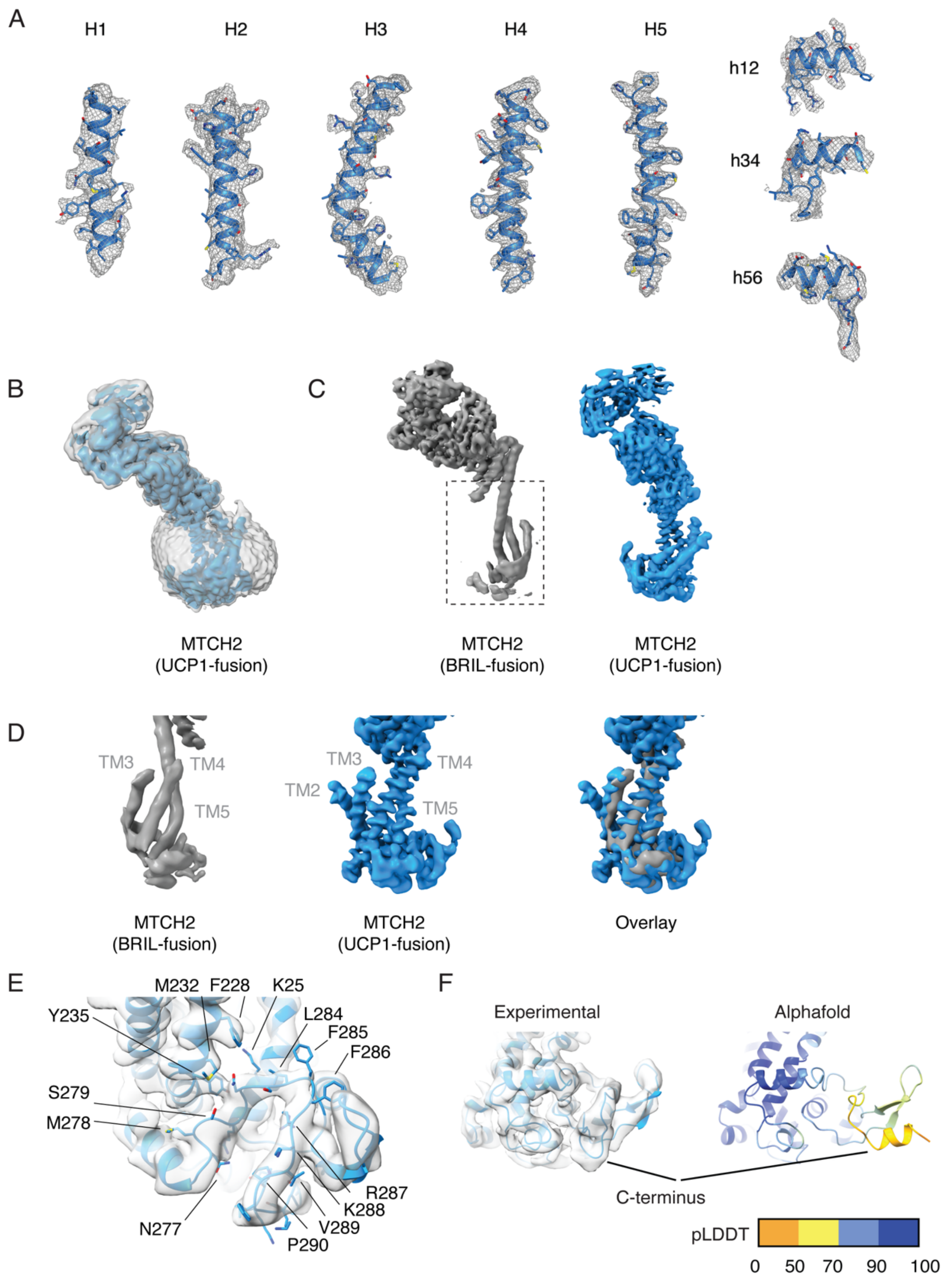


**Fig. S7. Cryo-EM map assessment and MTCH2 fusion constructs comparison. A)** Cryo-EM density map of MTCH2 (UCP1) displayed as mesh around the 5 TM helices, and helical linker h12, h34, and h56, and the C-terminus, highlighting overall map quality, with density for at least one or two prominent side chains visible on each helix. **B)** Low-threshold (gray, transparent) overlayed with high-threshold (blue, solid) cryo-EM density map of MTCH2 •α-UCP1 Nb•NabFab•α-Fab Nb complex highlighting MTCH2 embedded into UDM detergent micelle. **C)** Comparison between cryo-EM density map of MTCH2 (BRIL)•BAK5•α-Fab Nb complex (left, gray) and MTCH2•α-UCP1 Nb•NabFab•α-Fab Nb complex (right, blue). **D)** MTCH2-focused comparison of the EM density maps for MTCH2(BRIL) (gray) with that from MTCH2(UCP1) (blue). The overall conformation of MTCH2 in both fusions is similar, suggesting the fusions used for structure determination are not altering the conformation of MTCH2. **E)** Detailed model with side chains displayed for the MTCH2 C-terminus overlaid with cryo-EM density. **F)** C-terminus-focused comparison of the experimentally-determined and AlphaFold-predicted model (colored by pLDDT) of MTCH2 highlights major conformational differences. AlphaFold model confidence values are reported in table S6.

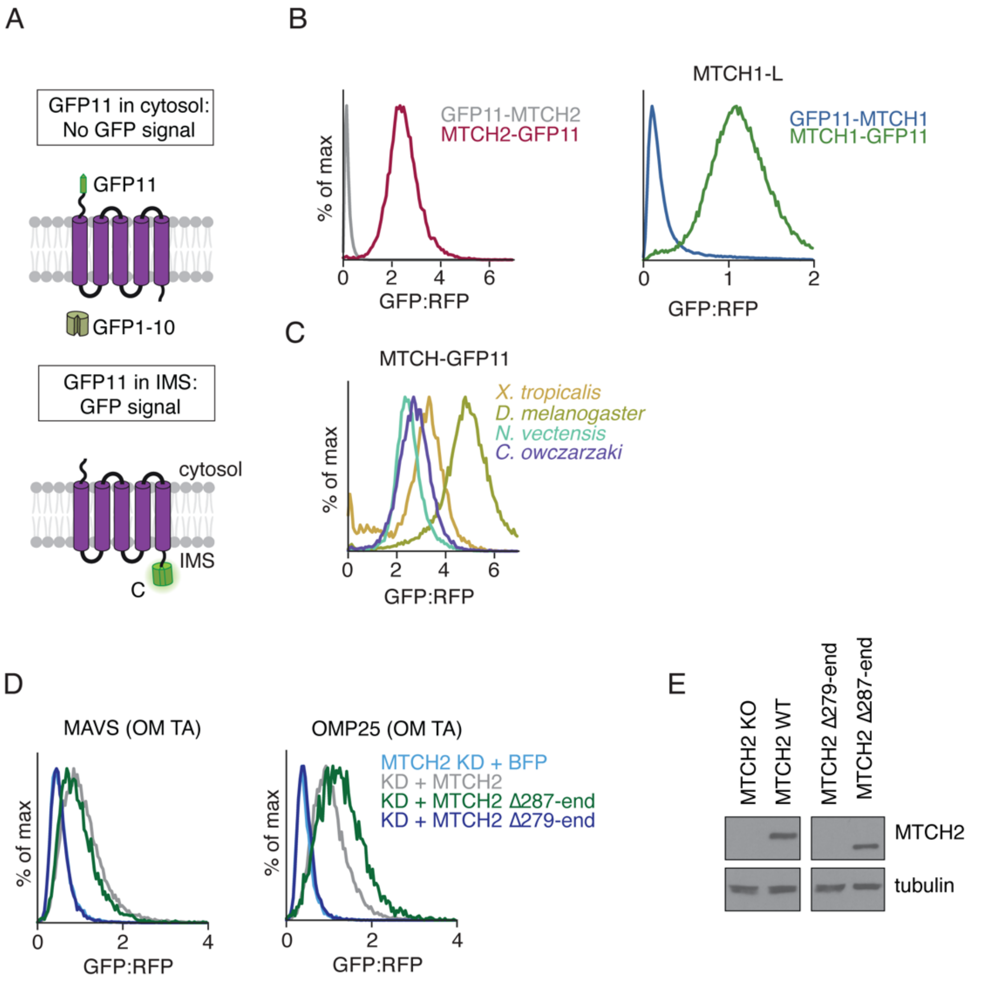


**Fig. S8. MTCH homologs adopt a conserved 5 TM topology, which is specifically stabilized by a portion of the C-terminus. A)** To experimentally confirm the topology of MTCH2 observed in the cryo-EM structure, we leveraged the split-GFP reporter system described in Fig. 1B. Here, GFP11 is fused to either the N- or C-terminus while GFP(1-10) is expressed in the IMS. If the GFP11-tagged terminus is in the IMS of the mitochondria, this will result in complementation and thus GFP fluorescence. Comparison of the GFP fluorescence of these two constructs, relative to a normalization marker (RFP), can be used determine protein topology. **B)** (top) Flow cytometry measurements of K562 IMS GFP(1-10) cells expressing RFP-P2A-tagged MTCH2 or MTCH1-L (long isoform, see fig. S2A) with GFP11 on either the N- (gray) or C-terminus (red). The C-terminal GFP11 tag has a much higher GFP:RFP for both proteins, consistent with positioning of their C-termini in the IMS, and their N-termini in the cytosol, as observed in the structure. Based on these data, together with the structure, we definitively concluded that MTCH2 contains five TMs. **C)** As in (B) for the indicated C-terminally GFP11 tagged MTCH homologs: *X. tropicalis*, *D. melanogaster, N. vectensis*, and *C. owczarzaki*. GFP complementation for all the homologs is similar to that for *H. sapiens* MTCH2-GFP11, suggesting the MTCH family all adopt same topology with 5 TMs and their C-termini in the IMS. **D)** Based on its position in the structure, we hypothesized that the C-terminal domain of MTCH2 could be important for stabilizing its remaining 5 TMs, and thereby the hydrophilic groove created by loss of the 6$^{th}$ TM. To test this, we generated two truncations to the C-terminus of MTCH2 and tested their ability to mediate insertion of OMP25 and MAVS in K562 CRISPRi cells expressing a MTCH2 targeting sgRNA (MTCH2 KD). Rescue constructs encoding either a BFP control, WT MTCH2, or MTCH2 with C-terminal truncations to position 287 or 279 were expressed to determine if they could rescue the MTCH2 KO phenotype. **E)** Immunoblotting analysis of MTCH2 was performed to verify the expression of the constructs from (C). To account for differences in transduction efficiency, samples are normalized to the % BFP positive cells. While the truncation at residue 287 retained insertase activity and expressed as normal, the truncation at residue 279 was destabilized, indicating an important role for residues 279-286 in stabilizing the fold of MTCH2.

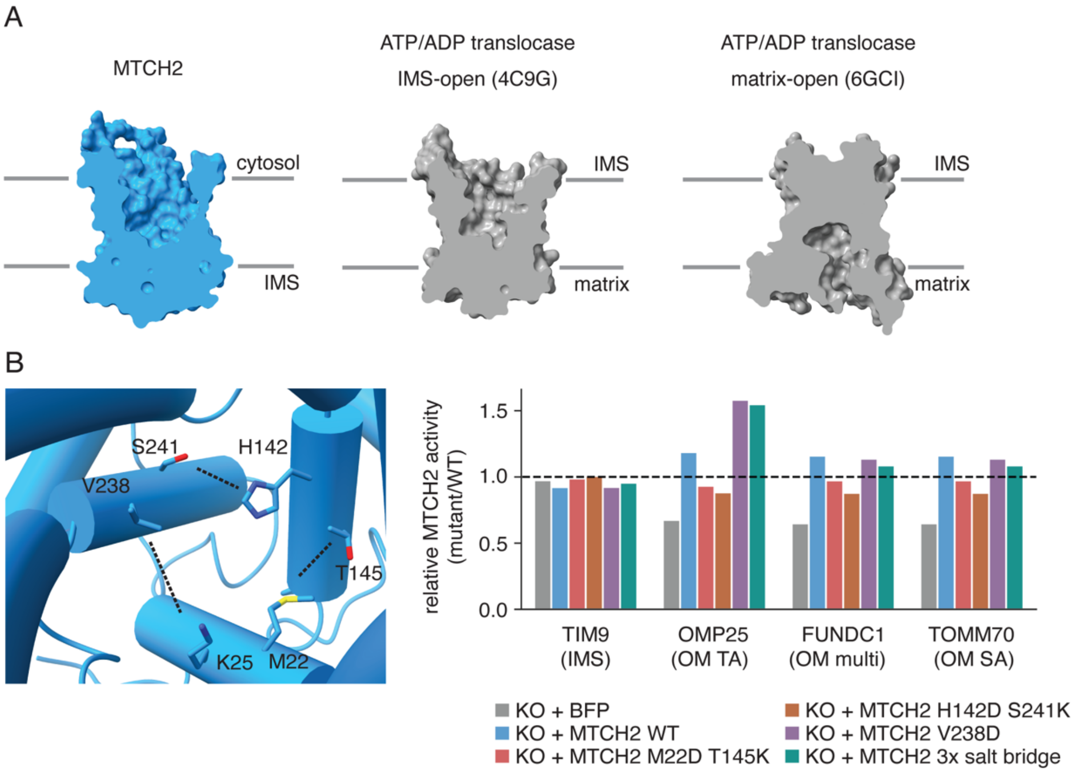


**Fig. S9. MTCH2 exhibits a cytoplasm-open conformation without stabilizing salt bridges. A)** Space-filling representation of the experimental MTCH2 structure and the SLC25 ATP/ADP translocase (AAC) in either an IMS-open (*26*) or matrix-open conformation (*25*) highlighting accessibility of the interior cavity from either side of the membrane. MTCH2 is stabilized in a conformation that is open towards the cytosol, consistent with its function in inserting OM proteins from a cytosolic substrate pool, despite the absence of canonical salt bridge-forming residues known to stabilize this conformation in other SLC25 carriers. **B)** Canonical SLC25s contain several conserved salt bridges between TM helices that enable the conformational changes required for the alternating access model of solute transport. The majority of these salt bridges are no longer found in the MTCH family of insertases, and we wondered if the loss of these interactions were important for MTCH2's insertase function. Therefore, we tested the effect of restoring these SLC25 salt bridges to MTCH2 on its insertion activity. (left) MTCH2 side chains are shown at positions where conserved salt-bridge forming residues are normally located. (right) Insertion of three MTCH2-dependent OM proteins, OMP25 and FUNDC1, and RHOT1, and the soluble IMS control TIM9 were measured in MTCH2 KO cells. Rescue constructs encoding either a BFP control, WT MTCH2, MTCH2 mutants with a single salt bridge restored at one of three positions, or a MTCH2 mutant with all three salt bridges restored were expressed to determine if they could rescue the MTCH2 KO phenotype. The activity of each MTCH2 mutant relative to WT MTCH2 (calculated as median $GFP:RFP_{mut}$/median $GFP:RFP_{wt}$) was determined. The presence of salt bridges either does not alter MTCH2 insertase function, or potentially enhances it in one position, V238. We therefore concluded that loss of these salt bridges was not a critical evolutionary adaptation of the MTCH family during their evolution from an SLC25 transporter.

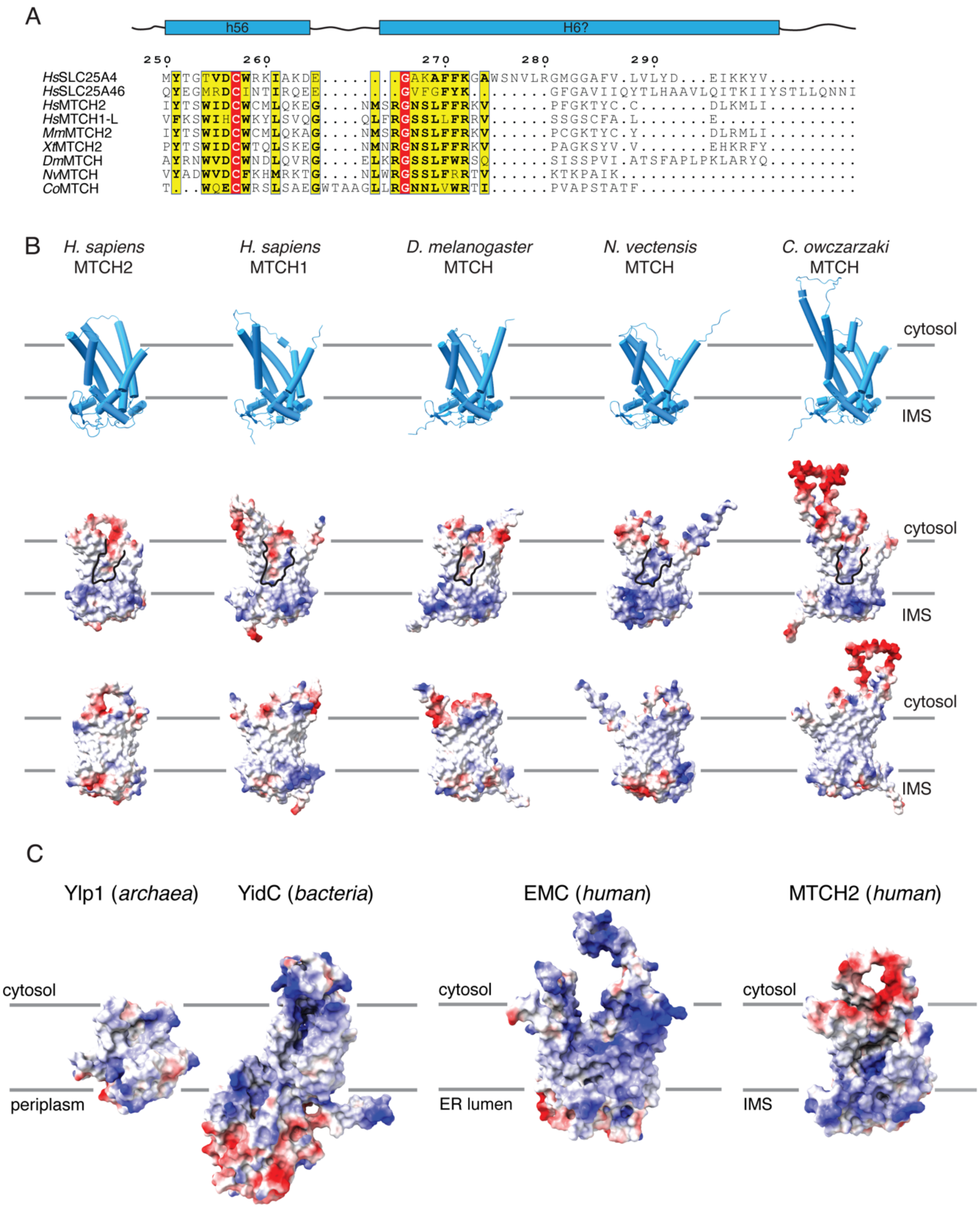


**Fig. S10. Identification of conserved features of the MTCH family of insertases. A)** MUSCLE-generated sequence alignment of the conserved C-terminal domain of the MTCH insertases that is not found in the SLC25 family. Residues 291-303 within this domain are modeled with very low confidence by AlphaFold (*23*, *24*), and its conservation and positioning suggested it may be functionally important. Labels above the alignment correspond to positions in canonical SLC25 transporters that typically encode the structural features h56 (a membrane-peripheral helix between H5 and H6) and H6. **B)** Comparison (from left to right) between our experimental MTCH2 model, and AlphaFold-predicted (*23*, *24*) models of human MTCH1, and MTCH homologs from *Drosophila melanogaster*, *Nematostella Vectensis*, and *Capsaspora Owczarzaki*. Cartoon rendered models are shown on top and surface-mapped coulombic electrostatic potential (calculated with ChimeraX) is shown in the middle (front-facing) and bottom rows (180° rotated). All homologs have a comparable fold. We have also experimentally confirmed that they each contain 5 TMs, can express in the OM of human cells (fig. S8A-C), and adopt the same topology as human MTCH2 with their C-termini in the IMS. The loss of TM6, when compared to an SLC25, results in a prominent, lipid-accessible hydrophilic groove, which is a conserved feature across the MTCH family of insertases. Structural modeling does not accurately predict the conformation of the C-terminal domain of human MTCH2, so it is not possible to interpret the structural conservation of this feature. However, sequence alignments in (A), suggest that this domain is also a conserved feature across the MTCH family of insertases that is not found in canonical SLC25s. Residues 1-70 of MTCH1, which are predicted with low confidence were omitted for simplicity. AlphaFold model confidence values are reported in table S6. **C)** Space-filling representation of human MTCH2 colored by electrostatic potential in comparison to other evolutionarily unrelated insertases. Highlighted is the presence of a conserved lipid-exposed hydrophilic groove, which is a unifying feature defining diverse insertase folds. Coulombic electrostatic potential was calculated using ChimeraX and mapped onto the surface of Ylp1 (PDB: 5C8J; (*92*)), YidC (PDB: 6Y86; (*93*)), EMC (PDB: 8S9S; (*46*)), and MTCH2.

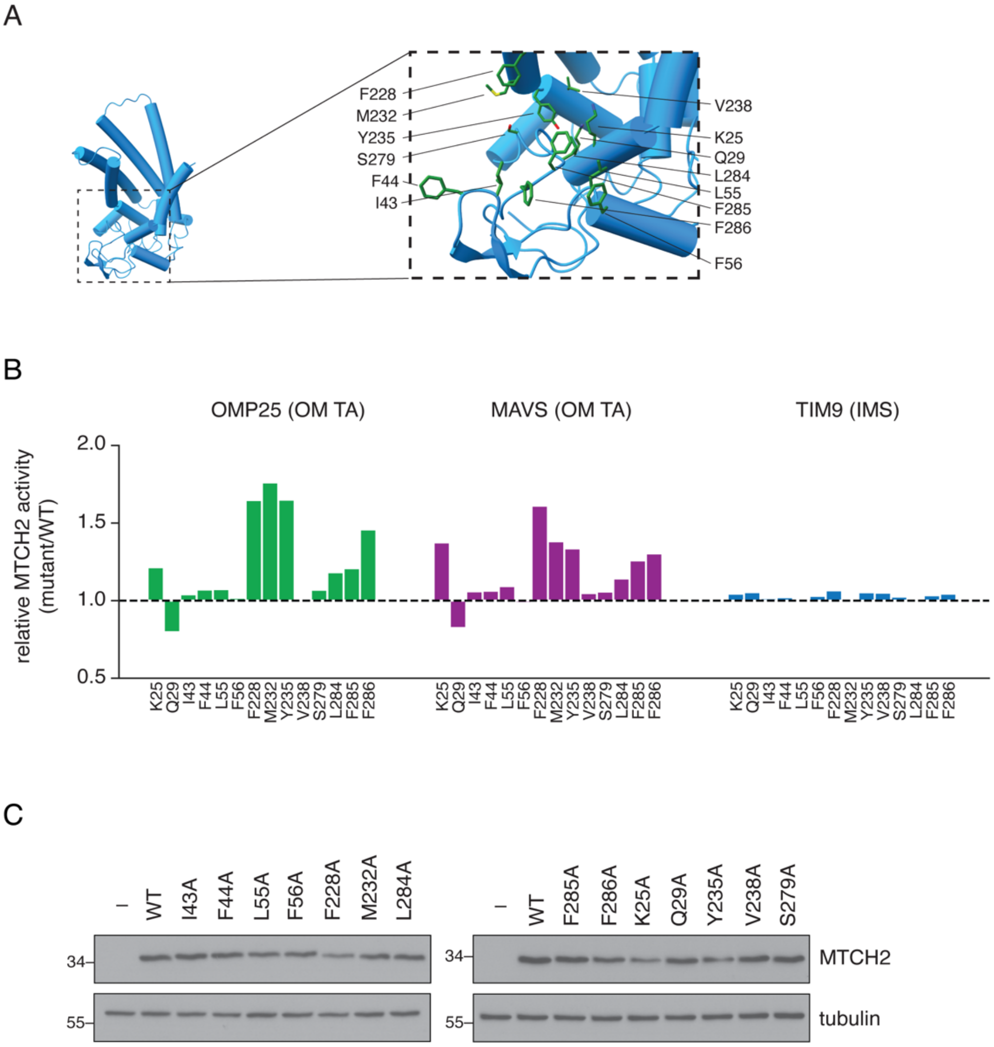


**Fig. S11. MTCH2 activity is attenuated by residues lining its hydrophilic groove. A)** The most distinctive feature of the MTCH homologs relative to other SLC25 family members is the absence of TM6, leaving a hydrophilic groove accessible to the bilayer. Given the functional significance of these types of hydrophilic grooves in other insertase families, we hypothesized that the amino acids that define this region would be important for MTCH2's insertase activity. To assess the importance of amino acids in this region, 14 sites (K25, Q29, I43, F44, L55, F56, F228, M232, Y235, V238, S279, L284, F285, F286) were chosen for initial focus in structure-function experiments. **B)** Alanine scanning mutagenesis was performed on MTCH2 at the indicated 14 sites in and adjacent to the base of the hydrophilic groove. In order to ensure that both activating and inhibiting mutations could be detected, we focused on substrates which had previously been shown depend directly on MTCH2 levels for their insertion. For example, the levels of some MTCH2 substrates, such as many multipass proteins, appear to be independently regulated and/or may be affected by the levels of factors other than MTCH2 (*8*, *22*). The ratiometric reporter system was used to measure the insertion of two OM proteins, OMP25 and MAVS, and the soluble IMS protein control, TIM9, in MTCH2 KO cells. Rescue constructs encoding WT MTCH2 and alanine mutations at each of the 14 positions were expressed to determine their effect on OM insertion. The activity of each MTCH2 mutant relative to WT

MTCH2 (calculated as median $GFP:RFP_{mut}$/median $GFP:RFP_{wt}$) was determined. Notably many of the mutations appear to increase insertion activity. This data is mapped onto the MTCH2 structure in Fig. 3B. **C)** Immunoblotting of the MTCH2 mutants in (B) was performed to exclude potential effects of protein expression and stability on activity. To control for differences in transduction efficiency, samples were normalized to % BFP positive cells.

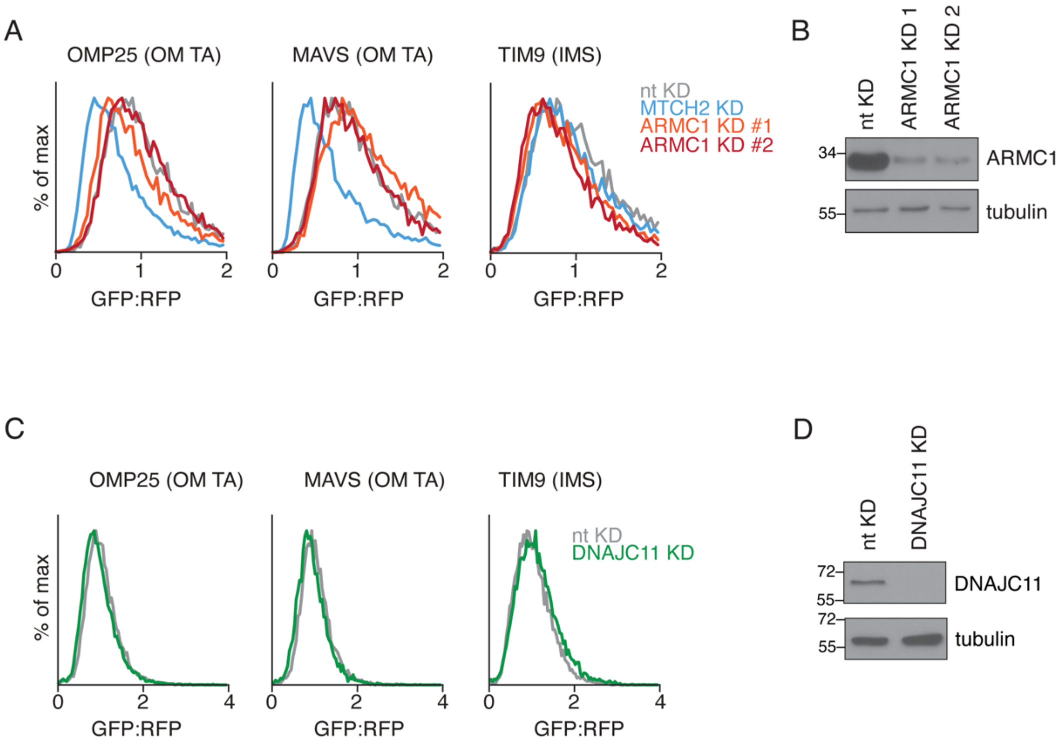


**Fig. S12. In K562 cells, MTCH2 activity is not attenuated by its two known binding partners, DNAJC11 and ARMC1. A)** One potential explanation for MTCH2's apparent attenuation, could be binding of an inhibitory regulatory molecule. We assessed whether this could be the case for one of the known binding partners of MTCH2, ARMC1 (*38*). To test whether depletion of ARMC1 in K562 cells would affect MTCH2's insertase function, we measured the insertion of the indicated OM proteins and soluble IMS control, in K562 CRISPRi cells expressing either a non-targeting (nt) or a ARMC1-targeting sgRNA. Results are normalized to nt, and displayed as a histogram. The results suggest that MTCH2 activity is not attenuated by endogenous ARMC1 in K562 cells. However, we cannot exclude the possibility that ARMC1 may regulate MTCH2 in other cell or tissue types where its relative expression may differ. **B)** Immunoblotting was performed to measure ARMC1 depletion in (A). **C,D)** As in (A, B), except for DNAJC11, since it is another MTCH2-interaction partner that could regulate its activity (*38*).

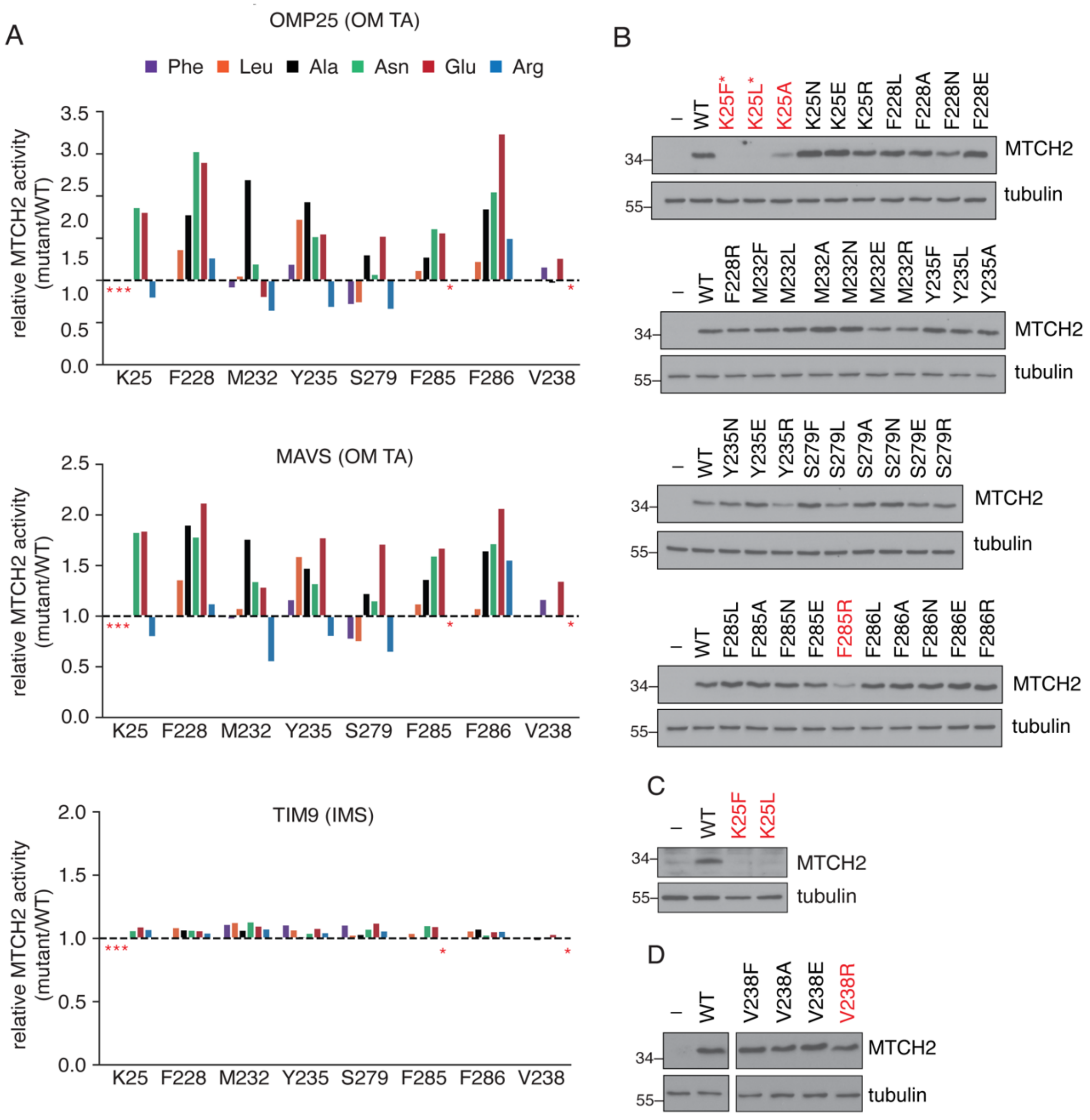


**Fig. S13. Systematic mutagenesis of MTCH2 to identify hyperactivity patterns. A)** A subset of 8 residues (K25, F228, M232, Y235, S279, F285, and F286, V238) from the mutants shown in fig. S11 were chosen for further interrogation. After observing hyperactivity from the alanine scanning analysis (fig. S11), we hypothesized that certain amino acids may have an inhibitory effect at these positions. To test this hypothesis, we mutated each position to a broader range of amino acids (Phe, Leu, Ala, Asn, Glu, and Arg) and tested the effects of these mutations on insertion of two OM proteins, OMP25 and MAVS, and a soluble IMS control, TIM9, in MTCH2 KO cells. Rescue constructs encoding WT MTCH2 and a total of 42 mutants were expressed to determine their effect on OM insertion. Relative activity was determined for each mutant compared to WT MTCH2. Mutations clearly affecting stability, as shown in B-D), were excluded from analysis at positions marked with a red asterisk. V238 mutational analysis was performed in a separate experiment where only Phe, Ala, Arg and Glu mutants were tested. **B)**

Immunoblotting of the MTCH2 mutants in (A) was performed, controlling for differences in transduction efficiency by normalizing to % BFP positive cells. Red font is used to denote cases where mutations are destabilizing, markedly decreasing MTCH2 expression (K25F, K25L, K25A, F285R). We concluded that we cannot interpret the effects of these mutations on MTCH2 activity. *K25F and K25L were not normalized for % BFP positivity in this set of immunoblots due to their toxicity during lentiviral production and thereby poor transduction efficiency. **C)** Immunoblotting with K25F and K25L mutants from (A) after normalizing to % BFP positive cells, confirming lack of expression. **D)** Immunoblotting of MTCH2 V238 mutants in (A) was performed, controlling for differences in transduction efficiency by normalizing to % BFP positive cells. Individual lanes were cropped from the same immunoblot at the same exposure level.

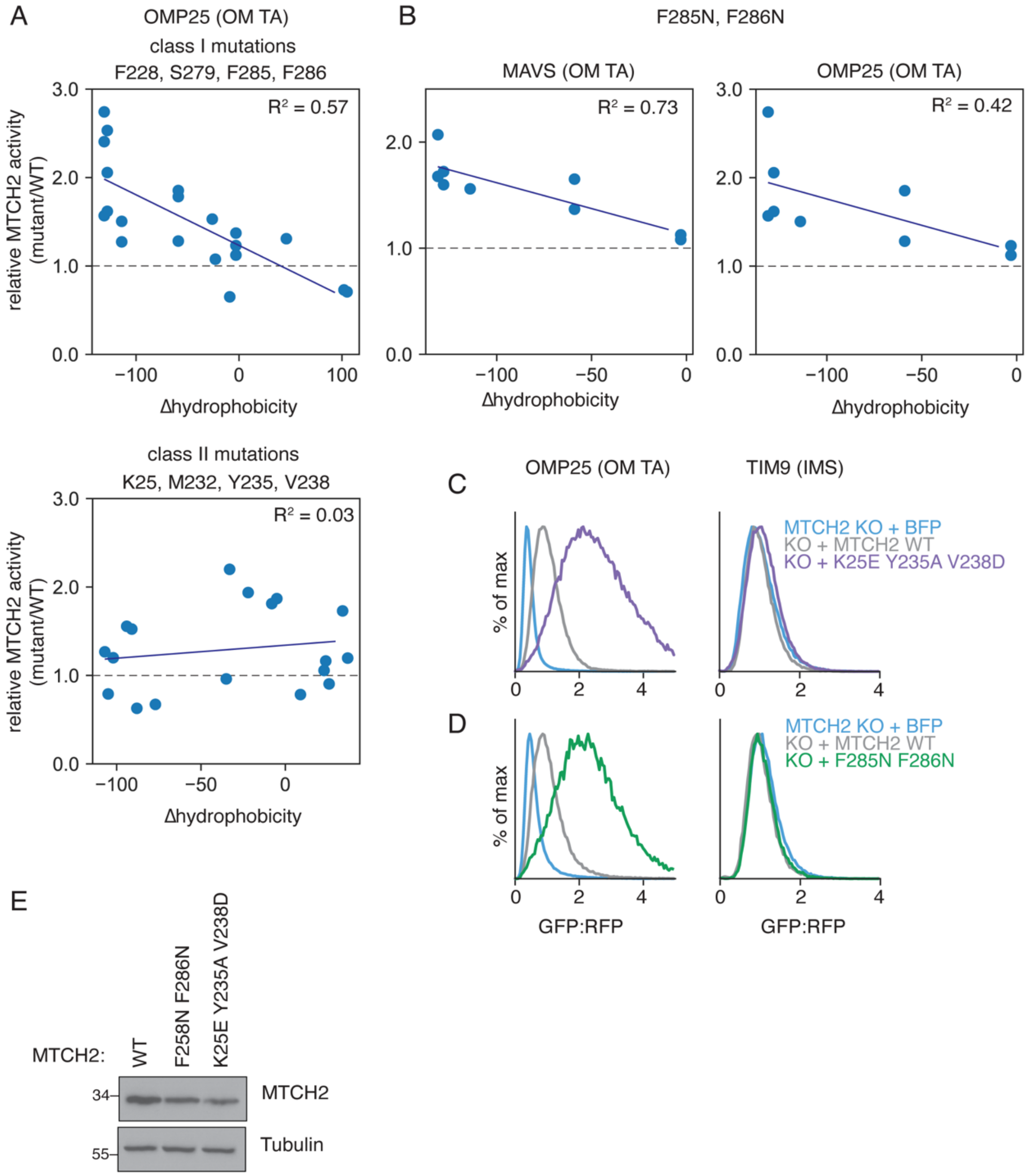


**Fig. S14. Two distinct classes of MTCH2 mutations result in hyperactivation. A)** Additional analysis related to Fig. 3D. (top) Data summarizing the effect of MTCH2 mutations (Phe, Leu, Ala,Gln, Glu, Arg) at 4 class I positions (F228, S279, F285, F286) on OMP25 insertion (full mutation panel shown in fig. S13A). The relative MTCH2 activity ($GFP:RFP_{mut}/GFP:RFP_{wt}$) plotted against the change in hydrophobicity of the mutated amino acid ($\Delta$Hydrophobicity=hydrophobicity$_{aa\ mut}$-hydrophobicity$_{aa\ wt}$), as determined using the indicated hydrophobicity scale (*77*). (bottom) data shown for 4 class II positions (K25, M232, Y235, V238). **B)** (left) Data summarizing the effect of mutations (Phe, Leu, Ala,Gln, Glu, Arg) at 2 positions (F285, F286) on MAVS insertion by MTCH2 mutants (full mutation panel shown in fig. S13A). The relative MTCH2 activity ($GFP:RFP_{mut}/GFP:RFP_{wt}$) plotted against the change in hydrophobicity of the mutated amino acid ($\Delta$Hydrophobicity=hydrophobicity$_{aa\ mut}$-

hydrophobicity$_{aa\ wt}$), as determined using a published hydrophobicity scale (*77*). (right) Effects for the same mutations on OMP25 insertion. **C-D)** Additional activity measurements for class I and class II combination mutants corresponding to data shown in Fig. 4A,C. We measured the insertion of indicated OM proteins and soluble IMS control, in MTCH2 KO cells. Rescue constructs encoding a BFP control, WT MTCH2, and either the class II combination mutant MTCH2$^{F285N,F286N}$ in (D) or the class I combination mutant MTCH2$^{F285N,F286N}$ in (E) were expressed to determine their effect on OM insertion. Results are normalized to MTCH2 WT and displayed as histograms. **E)** Immunoblotting of the MTCH2 mutants in (D-E) was performed, controlling for differences in transduction efficiency by normalizing to % BFP positive cells.

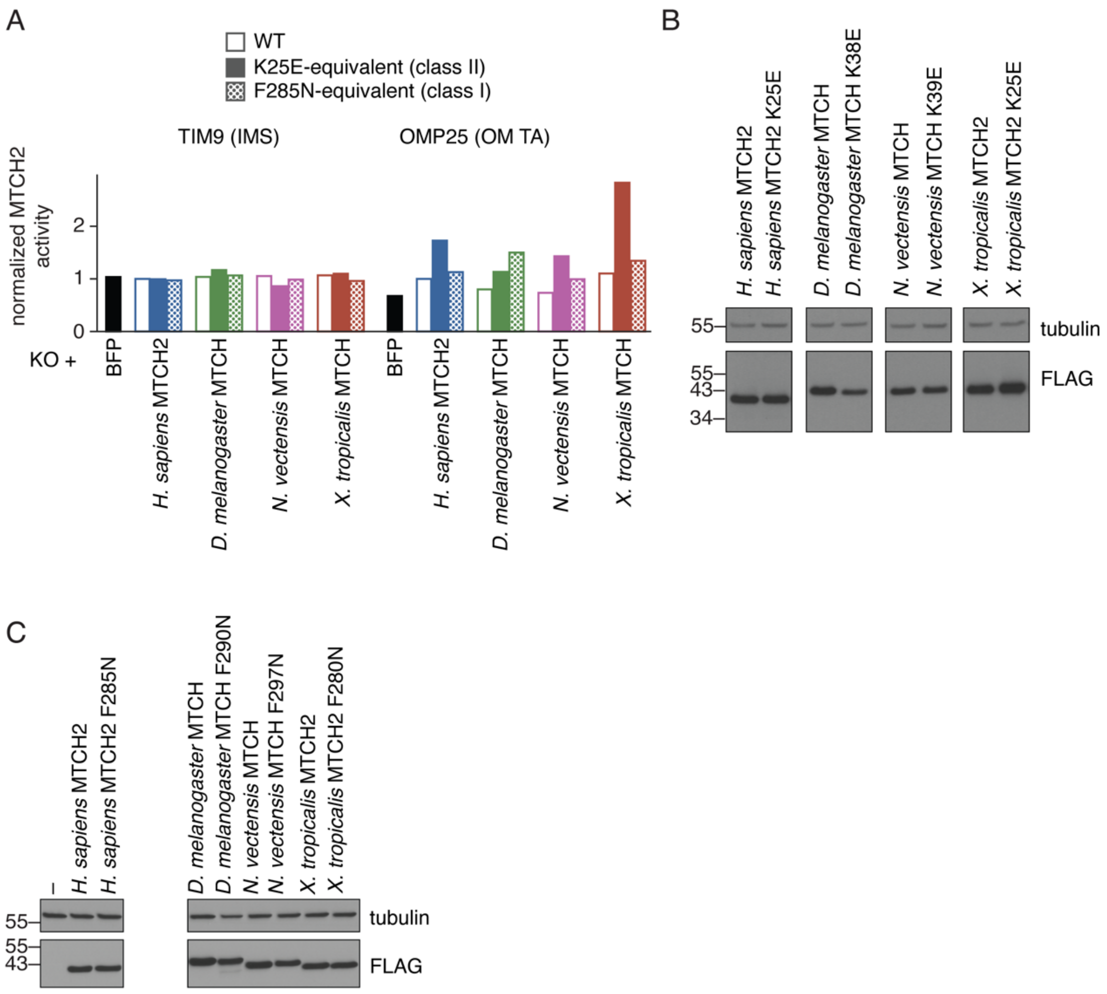


**Fig. S15. Activity of the MTCH homologs are all attenuated through a similar mechanism.** **A)** To test if the activity of the MTCH family of insertases relied on a similar mechanism to *H. sapiens* MTCH2, we introduced the equivalent mutation to the activating K25E to several MTCH homologs. The ratiometric reporter system was used to measure the insertion of indicated OM proteins and a soluble IMS control. Rescue constructs encoding a BFP control, WT MTCH2, and K25E-equivalent mutations and F285N-equivalent mutations to 3 MTCH homologs from *D. melanogaster, N. vectensis*, and *X. tropicalis*, all 3xFLAG-tagged, were expressed to determine their effect on OM insertion. Relative insertion of each construct was compared to WT MTCH2 (calculated as median GFP:RFP$_{mut}$/median GFP:RFP$_{wt}$). K25E-equivalent mutations were activating across all homologs tested, suggested a conserved mechanism of activity across all MTCH homologs. Additional data for this experiment is displayed in Fig. 3E. **B)** Immunoblot analysis was performed to confirm the expression of the MTCH homologs and the MTCH homolog K25E-equivalent mutants shown in (A). Individual lanes were cropped from the same immunoblot at the same exposure level. **C)** Immunoblot analysis was performed to confirm expression of the MTCH homologs and the MTCH homolog F285N-equivalent mutants shown in (A). Individual lanes were cropped from the same immunoblot at the same exposure level.

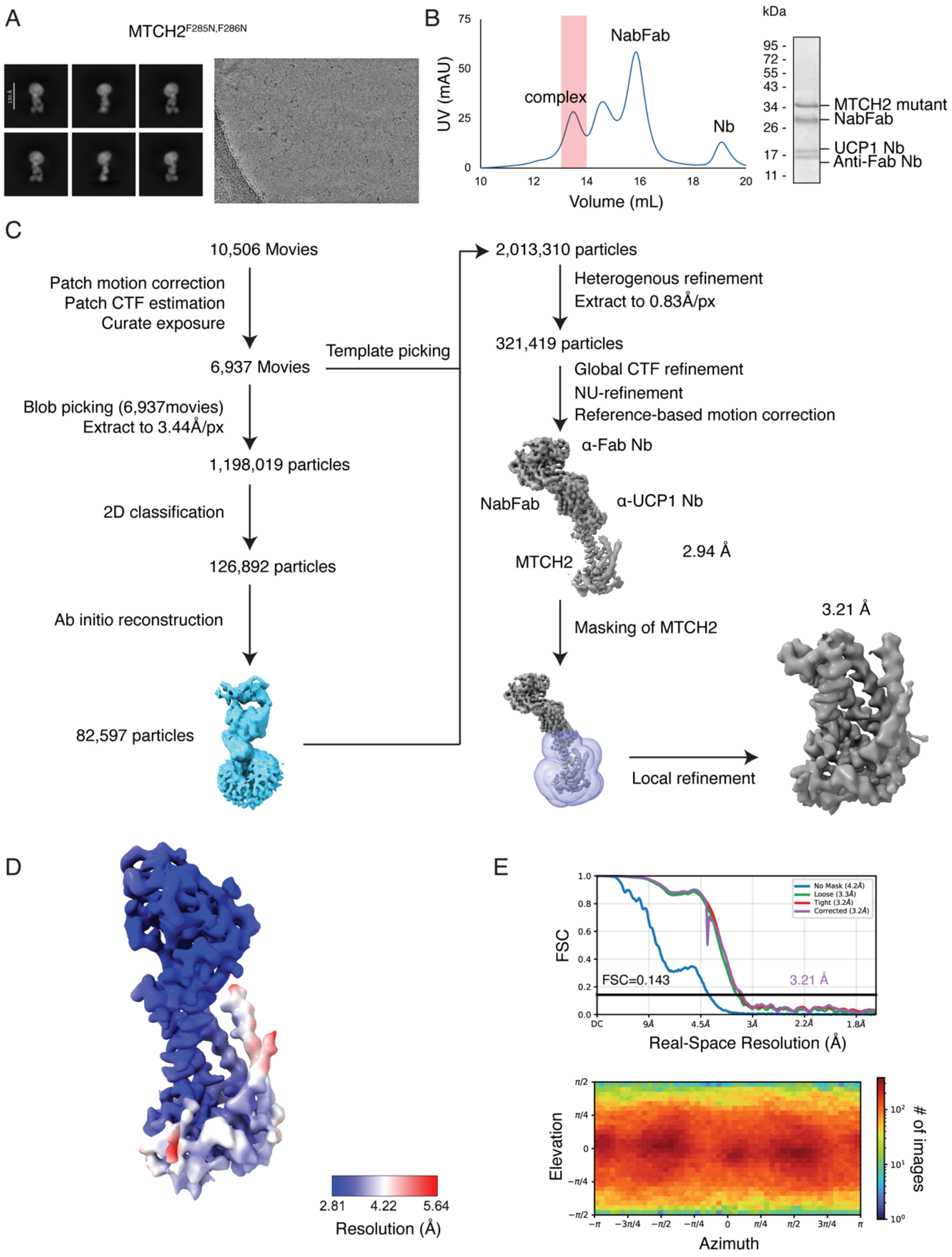


**Fig. S16. Cryo-EM data processing workflow for MTCH2$^{F285N,F286N}$. A)** (left) Representative 2D class averages of the MTCH2$^{F285N,F286N}$•α-UCP1 Nb•NabFab•α-Fab Nb complex, (right) and a micrograph from the data collection. **B)** (left) Size exclusion chromatography trace with the peak corresponding to the MTCH2$^{F285N,F286N}$•α-UCP1 Nb•NabFab•α-Fab Nb complex highlighted in red. (right) Pooled peak fractions were analyzed by SDS-PAGE and SYPRO Ruby stain (right). **C)** Data processing pipeline of MTCH2$^{F285N,F286N}$ using cryoSPARC v4.7.0. Micrographs were curated and a subset was used for blob picking. The resulting particle set was extracted at 3.44 Å/pixel and subject to iterative rounds of 2D classification. Particles from selected 2D classes were used in ab-initio reconstruction to generate 3D volumes. The volume representing a Fab attached to a detergent micelle was selected and used to generate 2D templates for template picker. After re-picking particles from the entire dataset, the resulting particle set was combined with the initial set and processed through iterative rounds of heterogenous refinement, yielding a final particle set of 321,419 particles. The particles were re-extracted at 0.83 Å/pixel and subject to global CTF refinement and reference-based motion correction. The final map was obtained from a non-uniform refinement job followed by the local refinement job using a mask on MTCH2 only. **D)** Cryo-EM density map of MTCH2$^{F285N,F286N}$•α-UCP1 Nb colored by local resolution in Å as calculated by cryoSPARC v4.7.0. **E)** Gold Standard Fourier Shell Correlation (GSFSC) curves (top) of the MTCH2 complex with a loose mask (green), tight mask (red), corrected mask (purple) or no mask (blue), and particle Euler angle distribution (bottom). A nominal resolution of 3.21 Å was determined for this map based off of an FSC cutoff of 0.143. Both plots were generated from the local refinement job in cryoSPARC v4.7.0.

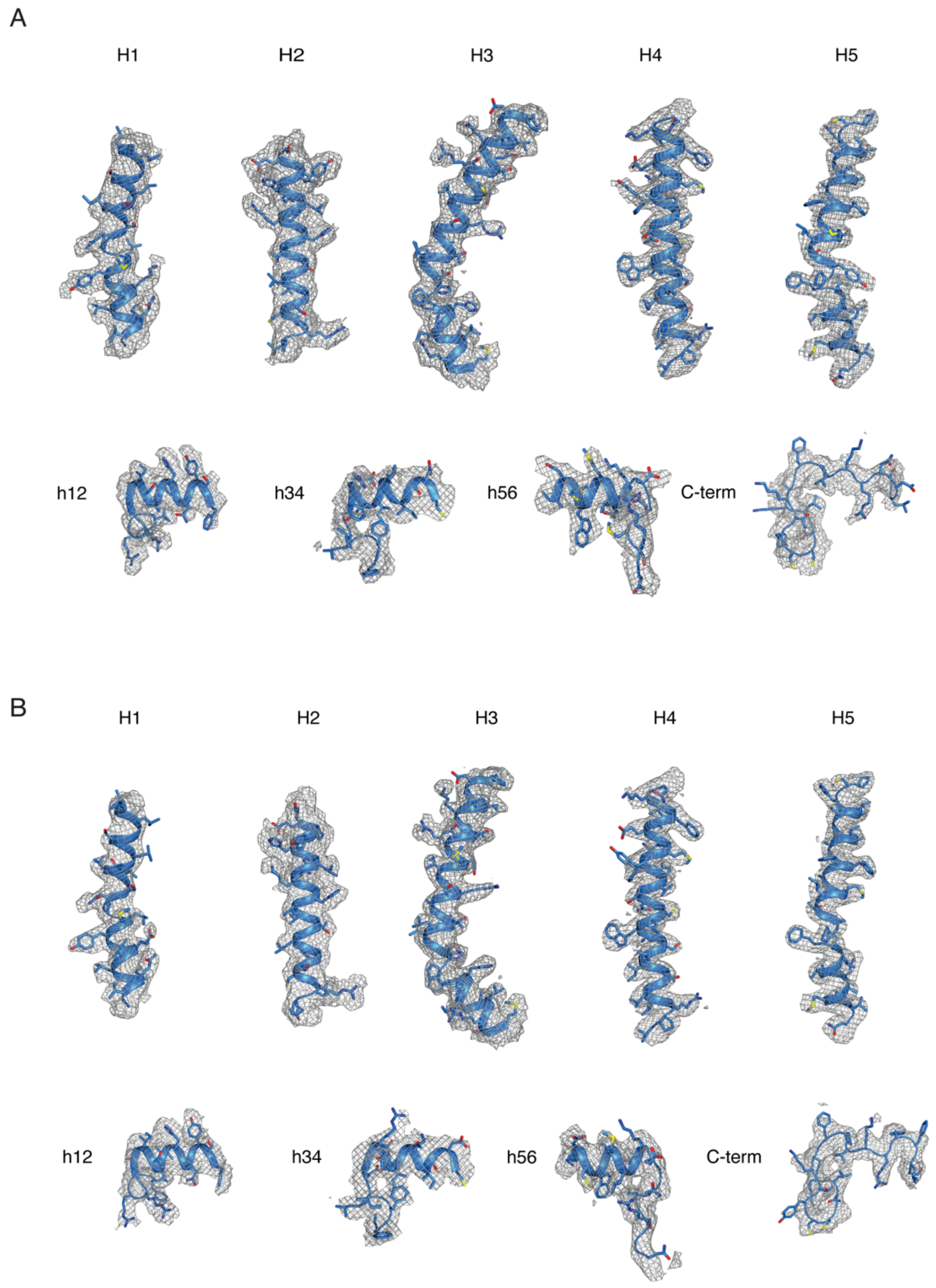


**Fig. S17. Representative density for the MTCH2$^{K25E,Y235A,V238D}$ and MTCH2$^{F285N,F286N}$ mutant structures. A)** Mesh density from the MTCH2$^{K25E,Y235A,V238D}$ cryo-EM map displayed around the 5 TM helices, helical linker h12, h34, h56, and the C-terminus, highlighting overall map quality, with density for at least one or two prominent side chains visible on each helix. **B)** Same as (A) for MTCH2$^{F285N,F286N}$.

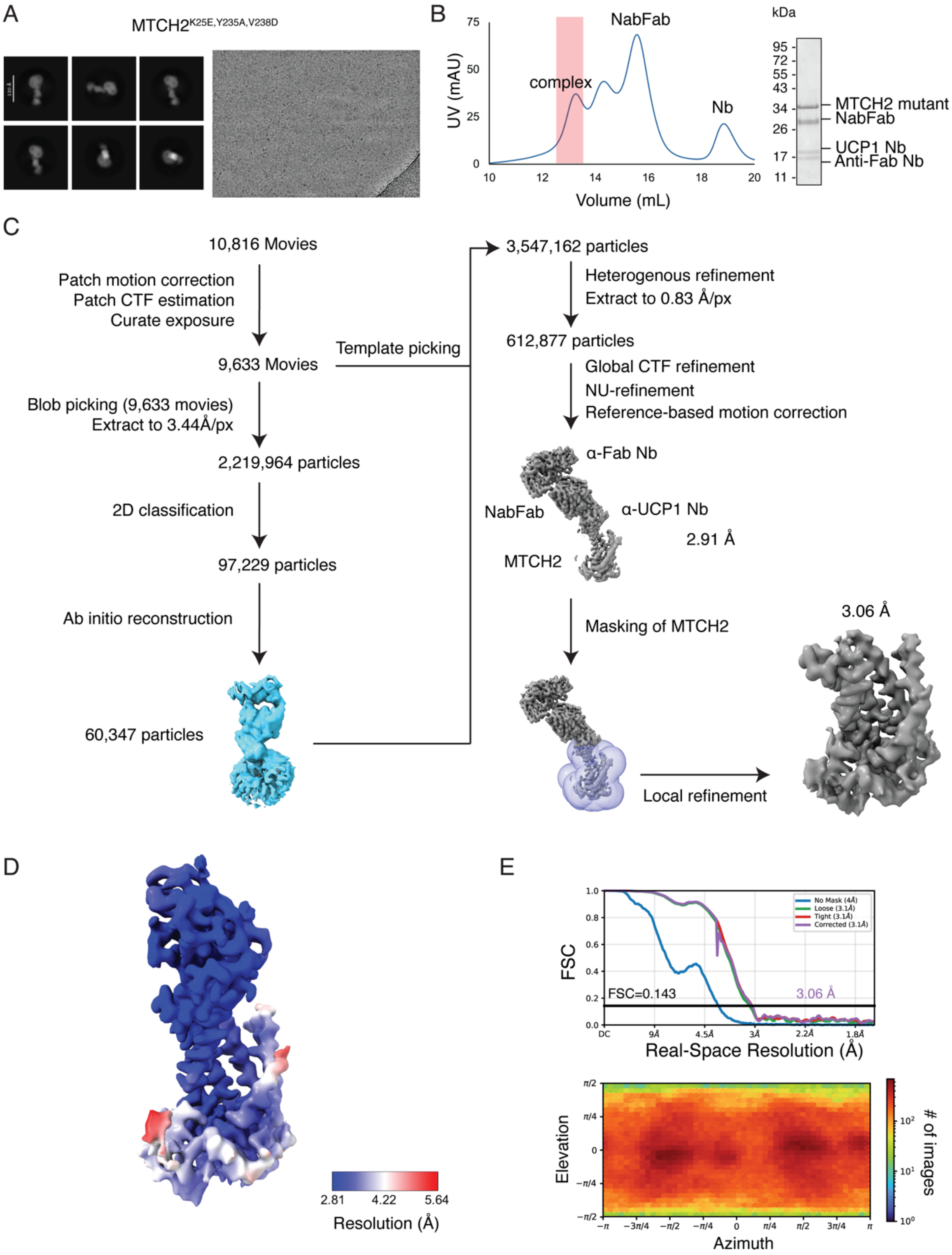


**Fig. S18**. **Cryo-EM data processing workflow for MTCH2$^{K25E,Y235A,V238D}$. A)** (left) Representative 2D class averages of the MTCH2$^{K25E,Y235A,V238D}$•α-UCP1 Nb•NabFab•α-Fab Nb complex, (right) and a micrograph from the data collection. **B)** (left) Size exclusion chromatography trace with the peak corresponding to the MTCH2$^{K25E,Y235A,V238D}$•α-UCP1 Nb•NabFab•α-Fab Nb complex highlighted in red. (right) Pooled peak fractions were analyzed by SDS-PAGE and SYPRO Ruby stain (right). **C)** Data processing pipeline of MTCH2$^{K25E,Y235A,V238D}$, same as that for MTCH2$^{F285N,F286N}$•α-UCP1 Nb•NabFab•α-Fab Nb complex described in S15C, resulting in a set of 612,877 particles. **D)** Cryo-EM density map of MTCH2$^{K25E,Y235A,V238D}$•α-UCP1 Nb colored by local resolution in Å as calculated by cryoSPARC v4.7.0. **E)** Gold Standard Fourier Shell Correlation (GSFSC) curves (top) of the MTCH2 complex with a loose mask (green), tight mask (red), corrected mask (purple) or no mask (blue), and particle Euler angle distribution (bottom). A nominal resolution of 3.06 Å was determined for this map based off of an FSC cutoff of 0.143. Both plots were generated from the local refinement job in cryoSPARC v4.7.0.

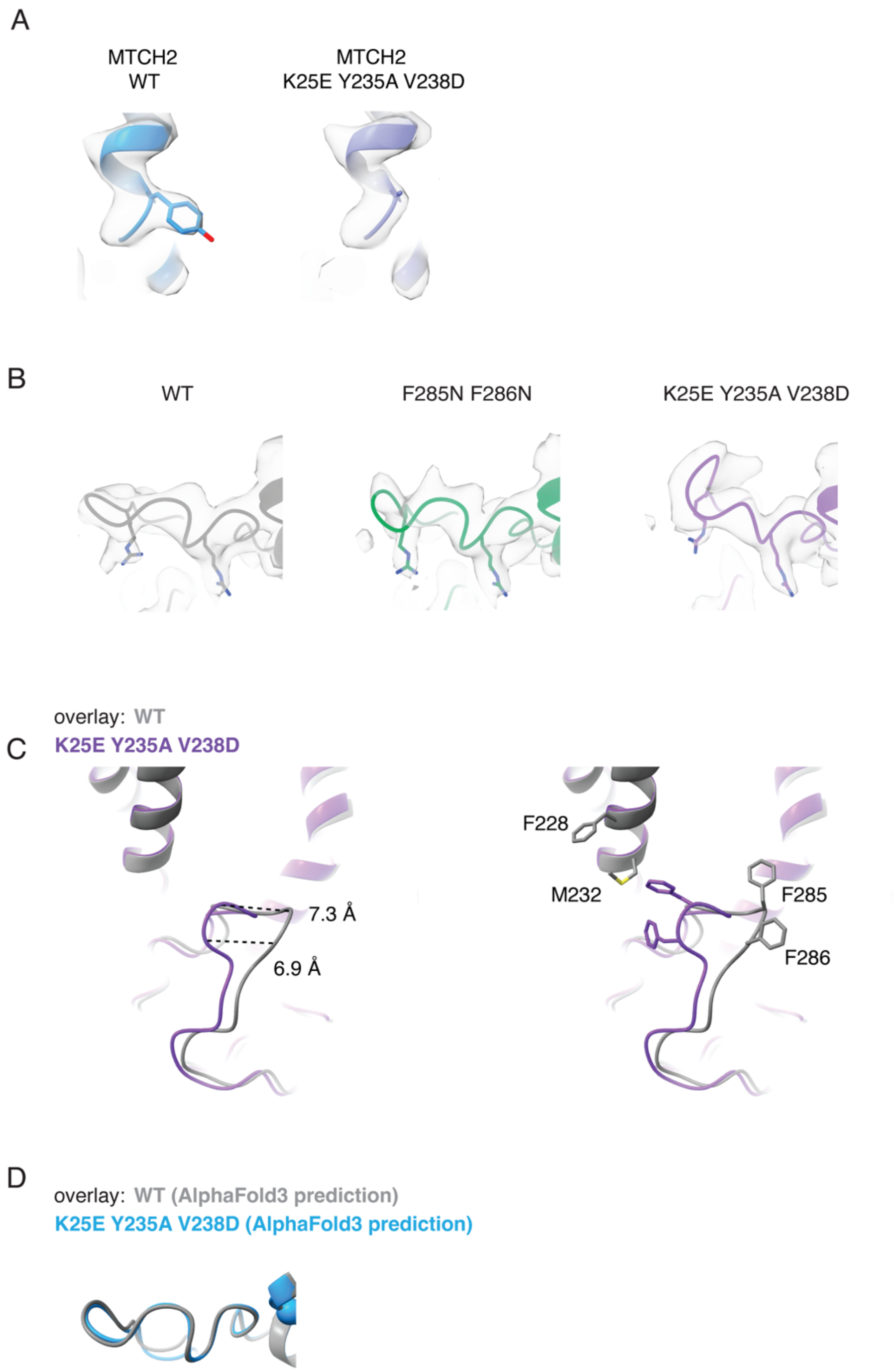


**Fig. S19. Cryo-EM density of the alternative C-terminal conformation in MTCH2$^{K25E,Y235A,V238D}$. A)** Comparison of the cryo-EM density surrounding Y235 in WT MTCH2 and MTCH2$^{K25E,Y235A,V238D}$ confirms the loss of density due to the alanine mutation. **B)** Comparison of cryo-EM density around the C-terminus for WT MTCH2, MTCH2$^{K25E,Y235A,V238D}$, and MTCH2$^{F285N,F286N}$. Prominent density for two arginine residues, R280 and R287, help to verify the correct sequence register and overall positioning of the peptide backbone for the region. **C)** (left) C-terminus centered view of MTCH2 WT (gray) overlayed with

MTCH2$^{K25E,Y235A,V238D}$ (purple), with measurements of the Cα displacement shown for F285 and F286. (right) The same overlay with side chains for F228, M232, F285, and F286 for WT and the altered F285 and F286 positions for MTCH2$^{K25E,Y235A,V238D}$. **D)** Overlay of the C-terminus of the AlphaFold-predicted models of WT MTCH2 and MTCH2$^{K25E,Y235A,V238D}$, showing no conformational change.

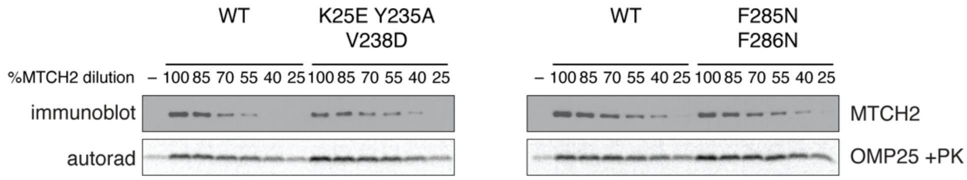


**Fig. S20. MTCH2 hyperactive mutants work by directly increasing MTCH2 insertase activity**. Corresponding data to Fig. 4F. Insertion of *in vitro* translated OMP25 in proteoliposomes containing either wiltype or mutant MTCH2 (F285N F286N or K25E Y235A V238D) was measured with a protease protection assay. To obtain matched wildtype and mutant MTCH2 levels, insertion was measured across a dilution series of proteoliposomes (100-25%).
(top) To quantify the relative amount of wildtype and mutant MTCH2 in each reaction, a dilution series of wildtype MTCH2 was measured by Western Blot. Using this dilution series, a linear regression was performed and applied to quantify the relative amount of each MTCH2 mutant utilized in each insertion reaction.
(bottom) SDS-PAGE and autoradiography was used to measure PK-protected OMP25 fragments in each sample. The % inserted for each wildtype and mutant MTCH2 was calculated relative to the maximum insertion observed for wildtype MTCH2 at its highest concentration (i.e. 100*[intensity of protease protected fragment MTCH2 mutant/intensity of protease protected fragment of wildtype MTCH2 at highest concentration]).

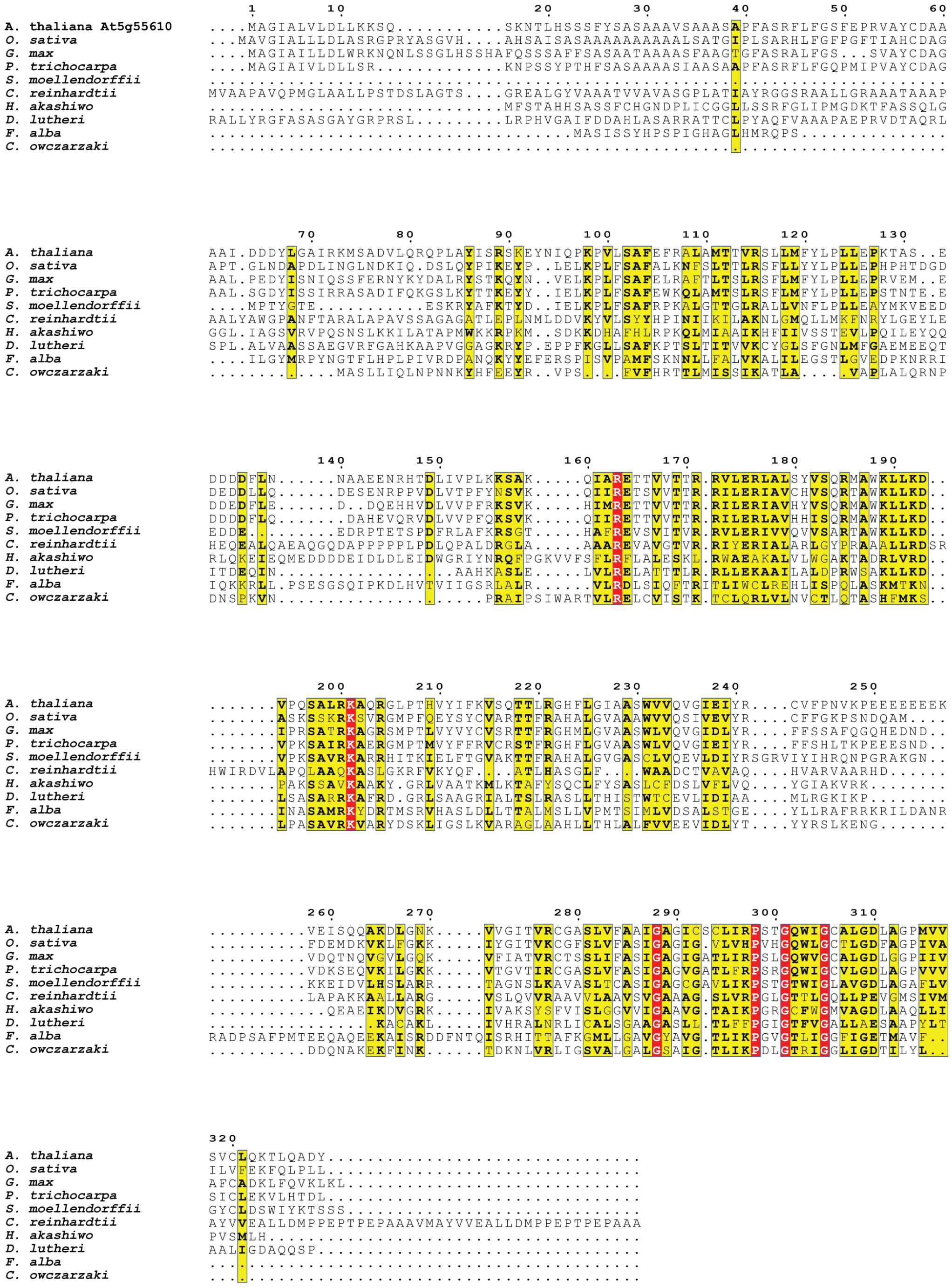


**Fig. S21. The At5g55610 sequence is conserved across plants and other eukaryotes**. Multiple sequence alignment of At5g55610 homologs from plants, *C. reinhardtii* (Chlorophyte), *H. akashiwo* (stramenopiles), *D. lutheri* (haptista), *C. owczarzaki* (holozoa), and *F. alba*

(holomycota) generated with MUSCLE (*74*) and visualized with the ESPript 3.2 server (*75*). The yellow regions correspond to positions where 70% or more of the residues have similar physicochemical properties, and the red regions correspond to positions where all residues have similar physicochemical properties.

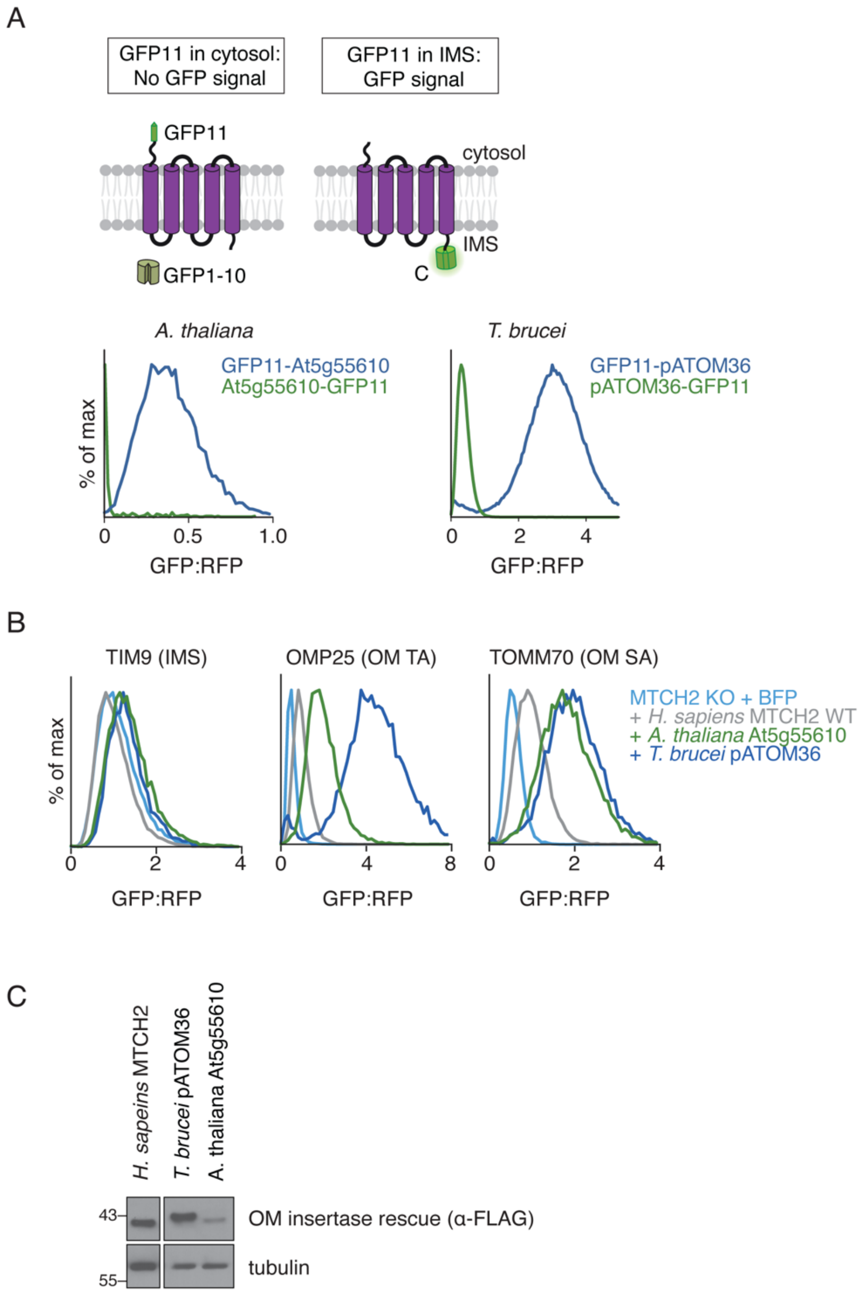


**Fig. S22. pATOM36 and At5g55610 localize to the OM and function as insertases in human cells. A)** Flow cytometry measurements of K562 IMS GFP(1-10) cells expressing RFP-P2A-tagged pATOM36 and At5g55610 with GFP11 on either the N- (blue) or C-terminus (green). N-terminal GFP11 tags have a much higher GFP:RFP ratio for both proteins, indicating that the N-terminus is in the IMS, while the C-terminus is in the cytosol. This is a reverse topology compared to MTCH homologs, and when analyzed together with the AlphaFold models (shown in Fig. 5B) suggests that pATOM36 and At5g55610 adopt an inverted fold compared to the MTCH homolog family. AlphaFold modeling suggested that these plant and protist OM insertases adopt an orientation that is more open towards the cytosol to facilitate their insertase function, similar to that observed for MTCH2. **B)** Additional data corresponding to Fig. 5C. We compared the insertase activity in human cells of MTCH2 to the known OM insertase from *T. brucei* pATOM36 and an OM protein from *A. thaliana* At5g55610 identified for its resemblance to MTCH2 and especially pATOM36. We used our ratiometric reporter assay to query indicated OM substrates and an IMS control. Rescue constructs encoding a BFP control, WT MTCH2, pATOM36, and At5g55610, all 3x-FLAG tagged, were expressed to determine their effect on OM insertion. Results are displayed as a histogram, normalized to WT MTCH2. **C)** Immunoblot analysis of was performed to compare expression levels of MTCH2 with At5g55610 and pATOM36 rescue constructs used in (A). Individual lanes were cropped from the same immunoblot at the same exposure level.

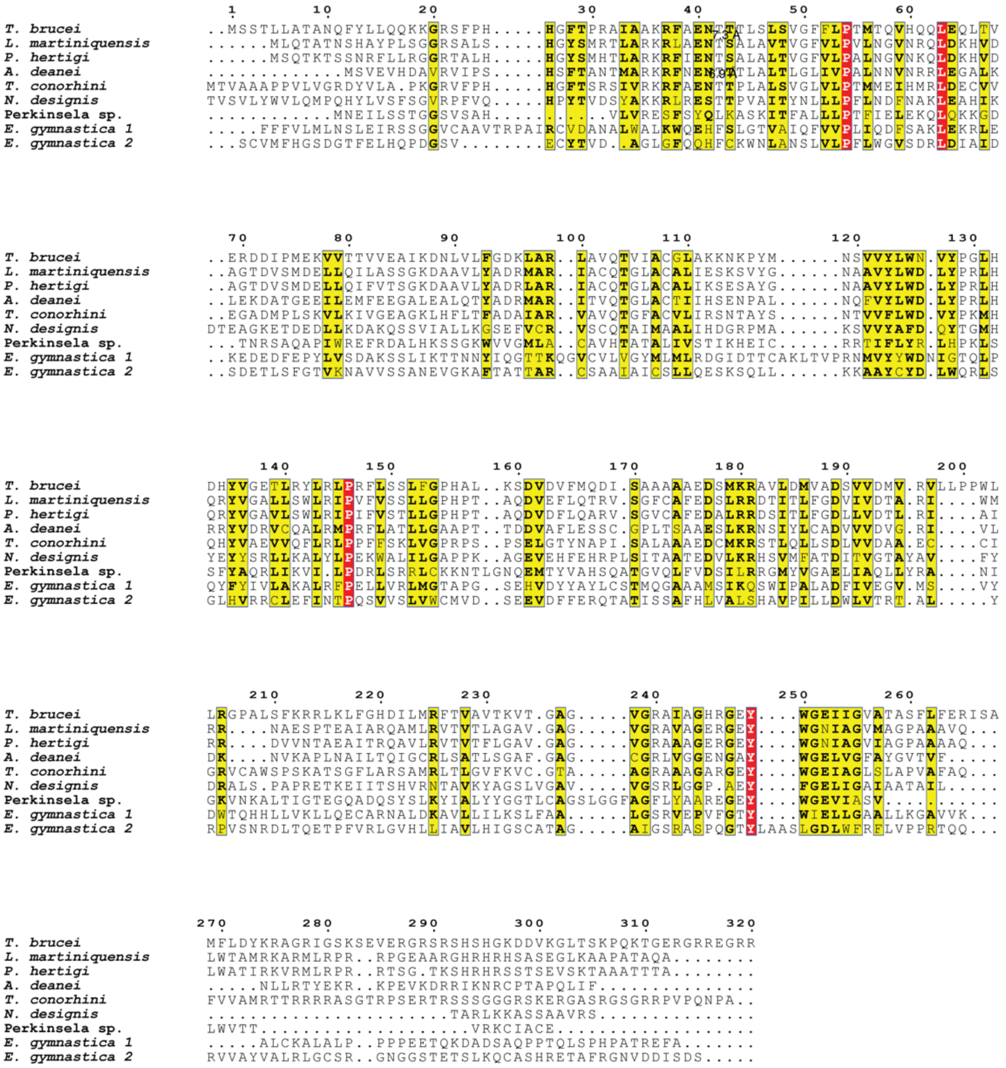


**Fig. S23. The pATOM36 sequence is conserved across Euglenozoa**. Multiple sequence alignment of pATOM36 homologs from Euglenozoa generated with MUSCLE (*74*) and visualized with the ESPript 3.2 server (*75*). The yellow regions correspond to positions where 70% or more of the residues have similar physicochemical properties, and the red regions correspond to positions where all residues have similar physicochemical properties.

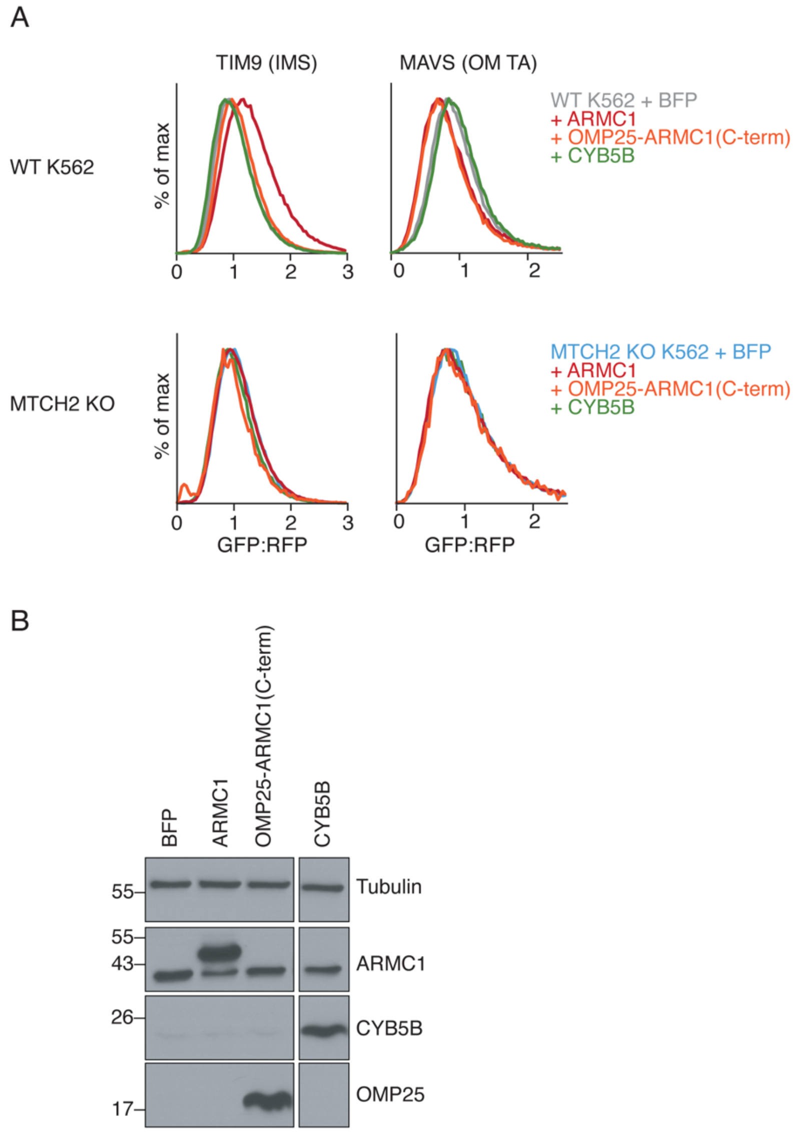


**Fig. S24. Overexpression of the ARMC1 C-terminus is both necessary and sufficient for inhibition of MTCH2 insertase activity**. **A)** The ratiometric reporter system was used to measure the insertion of the MTCH2-dependent OM protein MAVS and a soluble IMS control (TIM9). To test whether overexpression of ARMC1 (a known MTCH2 interaction partner) could inhibit MTCH2 insertase activity, the following constructs were transduced along with the each reporter: a BFP control; WT ARMC1; a chimera of OMP25 in which its C-terminal TM was replaced with the C-terminus of ARMC1 (OMP25-ARMC1[C-term]); or an unrelated MTCH2-dependent OM protein (CYB5B). All expression constructs included a BFP normalization marker to identify cells expressing the relevant fusions. Relative integration of MAVS and TIM9 into mitochondria was compared across each expression condition. The experiment was performed in both wildtype (WT) and MTCH2 knockout (KO) K562 IMS GFP(1-10) cells to determine whether the effect of each overexpressed protein was depended on the presence of MTCH2. Although inhibition of MTCH2 was not observed by endogenous ARMC1 in K562 cells (Fig. S12A), partial inhibition of MTCH2 by overexpression of the C-terminus of ARCM1

could be consistent with regulation of MTCH2 in cell or tissue types with higher ARMC1 levels.
**B)** Immunoblot analysis was performed to assess expression levels of the indicated proteins in (A).

**Table S1.** Cryo-EM data collection, refinement and validation statistics

| | MTCH2 (BRIL) (EMDB-76658) | MTCH2 (UCP1) (EMD-76655) (PDB 12OY) | MTCH2 F285N F286N (EMD-76656) (PDB 12OZ) | MTCH2 K25E Y235A V238D (EMD-76659) (PDB 12PB) |
|---|---|---|---|---|
| **Data collection and processing** | | | | |
| Magnification (nominal) | 105,000 | 105,000 | 105,000 | 105,000 |
| Voltage (kV) | 300 | 300 | 300 | 300 |
| Electron exposure (e–/Å$^2$) | 60 | 60 | 60 | 60 |
| Defocus range (µm) | -0.8 to -1.8 | -0.8 to -1.8 | -0.8 to -1.8 | -0.8 to -1.8 |
| Pixel size (Å) | 0.832 | 0.832 | 0.832 | 0.832 |
| Initial particle images (no.) | 7,786,491 | 4,791,546 | 3,547,162 | 2,013,310 |
| Final particle images (no.) | 199,397 | 485,622 | 612,877 | 321,419 |
| Map resolution (Å) | 3.55 | 3.57 | 3.06 | 3.21 |
| FSC threshold | 0.143 | 0.143 | 0.143 | 0.143 |
| Map resolution range (Å) | 3.0-9.8 | 2.9-7.4 | 2.5-8.5 | 2.7-7.4 |
| **Refinement** | | | | |
| Software (phenix.real_space_refine) | N/A | PHENIX 1.21.2-5419-000 | | |
| Initial model used (PDB code) | AlphaFold | AlphaFold | AlphaFold | AlphaFold |
| Correlation coefficient ($CC_{mask}$) | | 0.85 | 0.83 | 0.83 |
| Map sharpening *B* factor (Å$^2$) | | 0 | 0 | 0 |
| Model composition | | | | |
| Non-hydrogen atoms | | 3301 | 3295 | 3295 |
| Protein residues | | 426 | 426 | 426 |
| *B* factors (Å$^2$) | | | | |
| Protein (min/max/mean) | | 71/265/136 | 67/340/177 | 80/355/194 |
| R.m.s. deviations | | | | |
| Bond lengths (Å) | | 0.002 | 0.003 | 0.002 |
| Bond angles (°) | | 0.477 | 0.406 | 0.444 |
| Validation | | | | |
| MolProbity score | | 1.24 | 1.22 | 1.32 |
| Clashscore | | 4.25 | 2.74 | 3.80 |
| Poor rotamers (%) | | 0.56 | 0.85 | 0.56 |
| Cβ outliers (%) | | 0.00 | 0.00 | 0.00 |
| CaBLAM outliers (%) | | 1.44 | 2.15 | 1.67 |
| Ramachandran plot | | | | |
| Favored (%) | | 97.87 | 97.16 | 97.16 |
| Allowed (%) | | 2.13 | 2.84 | 2.84 |
| Disallowed (%) | | 0.00 | 0.00 | 0.00 |

**Table S2.** Plasmids used

| Plasmid | Backbone | Insert | Source |
|---|---|---|---|
| | | Split GFP-reporter plasmids | |
| pAG391 | UCOE EF-1α | RFP-P2A-MAVS-GFP11 | Guna et al. (*8*) |
| pAG397 | UCOE EF-1α | RFP-P2A-OMP25-GFP11 | Guna et al. (*8*) |
| pAG205 | UCOE EF-1α | RFP-P2A-MICU1(1-60)-GFP11-Sec61B | Guna et al. (*8*) |
| pAG206 | UCOE EF-1α | RFP-P2A-TIM9A-GFP11 | Guna et al. (*8*) |
| pMH409 | UCOE EF-1α | RFP-P2A-RHOT2-GFP11 | Guna et al. (*8*) |
| pAG435 | UCOE EF-1α | RFP-P2A-FUNDC1-GFP11 | Guna et al. (*8*) |
| pAG648 | UCOE EF-1α | RFP-P2A-GFP11-TOMM70 | Guna et al. (*8*) |
| pAG430 | UCOE EF-1α | RFP-P2A-RHOT1-GFP11 | Guna et al. (*8*) |
| pTS311 | UCOE EF-1α | RFP-P2A-GFP11-At5g55610-3xFLAG | this study |
| pTS702 | UCOE EF-1α | RFP-P2A-3xFLAG-At5g55610-GFP11 | this study |
| pTS331 | UCOE EF-1α | RFP-P2A-ATOM36-GFP11 | this study |
| pAG658 | UCOE EF-1α | RFP-P2A-GFP11-ATOM36-1xFlag | this study |
| pAG670 | UCOE EF-1α | RFP-P2A-C. owczarzaki MTCH2-GFP11 | this study |
| pTS686 | UCOE EF-1α | RFP-P2A-GFP11-C. owczarzaki MTCH2 | this study |
| pTS684 | UCOE EF-1α | RFP-P2A-MTCH1-L-GFP11 (Q9NZJ7-1) | this study |
| pTS685 | UCOE EF-1α | RFP-P2A-GFP11-MTCH1-L (Q9NZJ7-1) | this study |
| pAG553 | UCOE EF-1α | RFP-P2A-GFP11-MTCH2 | this study |
| pAG554 | UCOE EF-1α | RFP-P2A-MTCH2-GFP11 | this study |
| pAG708 | UCOE EF-1α | RFP-1xFLAG-M. musculus MTCH2-GFP11 | this study |
| pAG676 | UCOE EF-1α | RFP-1xFLAG-X. tropicalis MTCH2-GFP11 | this study |
| pAG673 | UCOE EF-1α | RFP-1xFLAG-D. melanogaster MTCH2-GFP11 | this study |
| pAG674 | UCOE EF-1α | RFP-1xFLAG-N. vectensis MTCH2-GFP11 | this study |
| | | Mammalian rescue plasmids | |
| pAG496 | UCOE EF-1α | BFP-P2A-MTCH2 | Guna et al. (*8*) |
| pAG301 | UCOE EF-1α | BFP | Guna et al. (*8*) |
| pTS682 | UCOE EF-1α | BFP-P2A-MTCH2 (UCP1) | this study |
| pTS683 | UCOE EF-1α | BFP-P2A-MTCH2 (BRIL) | this study |
| pTS244 | UCOE EF-1α | BFP-P2A-3xFlag-MTCH2 | this study |
| pAG718 | UCOE EF-1α | BFP-P2A-3xFlag-C. owczarzaki MTCH | this study |
| pAG719 | UCOE EF-1α | BFP-P2A-3xFlag-D. melanogaster MTCH | this study |
| pMH511 | UCOE EF-1α | BFP-P2A-3xFlag-D. melanogaster MTCH K38E | this study |
| pTS909 | UCOE EF-1α | BFP-P2A-3xFlag-D. melanogaster MTCH F290N | this study |
| pAG720 | UCOE EF-1α | BFP-P2A-3xFlag-N. vectensis MTCH | this study |
| pMH514 | UCOE EF-1α | BFP-P2A-3xFlag-N. vectensis MTCH K39E | this study |
| pTS912 | UCOE EF-1α | BFP-P2A-3xFlag-N. vectensis MTCH F297N | this study |
| pAG721 | UCOE EF-1α | BFP-P2A-3xFLAG-X. tropicalis MTCH2 | this study |
| pMH517 | UCOE EF-1α | BFP-P2A-3xFLAG-X. tropicalis MTCH2 K25E | this study |
| pTS906 | UCOE EF-1α | BFP-P2A-3xFlag-X. tropicalis MTCH2 F280N | this study |
| pAG710 | UCOE EF-1α | BFP-P2A-3xFLAG-M. musculus MTCH2 | this study |
| pAG507 | UCOE EF-1α | BFP-P2A-MTCH1-L (Q9NZJ7-1) | this study |
| pCL16 | UCOE EF-1α | BFP-P2A-MTCH1-S (Q9NZJ7-2) | this study |
| pAG569 | UCOE EF-1α | BFP-P2A-SLC25A46 | this study |

| | | | |
|---|---|---|---|
| pTS783 | UCOE EF-1α | BFP-P2A-CYB5B | this study |
| pTS788 | UCOE EF-1α | BFP-P2A-3xFlag-ARMC1 | this study |
| pTS790 | UCOE EF-1α | BFP-P2A-OMP25-ARMC1(C-term) | this study |
| Lentiviral packaging plasmids | | | |
| psPax2 | | | gift from Didier Trono (Addene #12260) |
| pMD2.G | | | gift from Didier Trono (Addene #12259) |
| Baculovirus generation plasmids | | | |
| pZL5 | pFastBac1 | GFP-SUMO-MTCH2(BRIL fusion) | this study |
| pZL10 | pFastBac1 | GFP-SUMO-MTCH2(UCP1 fusion) | this study |
| pZL15 | pFastBac1 | GFP-SUMO-MTCH2(UCP1 fusion) F285N F286N | this study |
| pZL16 | pFastBac1 | GFP-SUMO-MTCH2(UCP1 fusion) K25E Y235A V238D | this study |
| Mammalian expression plasmids | | | |
| pTS168 | pHAGE2 CMV | GFP-SUMO-MTCH2 | Guna et al. (*8*) |
| pMH474 | pHAGE2 CMV | GFP-SUMO-MTCH2 K25E Y235A V238D | this study |
| pTS859 | pHAGE2 CMV | GFP-SUMO-MTCH2 F285N F286N | this study |
| E. coli expression plasmids | | | |
| pTS117 | pQE | 14xHis-avi-GFP Nb | Stevens et al. (*67*) (Addgene #199370) |
| pAV286 | pDG | SENPEu | Vera Rodriguez et al. (*68*) (Addgene #149333) |
| pTS534 | pET Duet | Pelb-14xHis-TEV-anti-UCP1 Nb TC-NB4 scaffold | This study |
| pZL2 | pET Duet | Pelb-14xHis-TEV-anti-Fab Nb | Li et al. (*94*) |
| | pRH2.2 | NabFab | Bloch et al. (*30*) |
| | pRH2.2 | anti BRIL sAB BAK5 | Mukherjee et al. (*28*) |
| In vitro translation plasmids | | | |
| pAI351 | SP64 | OMP25 | Guna et al. (*8*) |
| pAI379 | SP64 | VHP-RHOT1 | Guna et al. (*8*) |
| gTS10 | SP64 | SU9-DHFR | Guna et al. (*8*) |

**Table S3.** CRISPRi sgRNA plasmids used.

| Plasmid | Backbone | sgRNA | sgRNA sequence | Source |
|---|---|---|---|---|
| pAG285 | pU6-sgRNA EF-1α-Puro | Non-targeting | GAACGACTAGTTAGGCGTGTA | Guna et al. (*8*) |
| pAG491 | pU6-sgRNA EF-1α-Puro | MTCH2 dual | GACCGGCTCACCGGGTCGCT, GGGCTCACCGGGTCGCTTGG | Guna et al. (*8*) |
| pAG692 | pU6-sgRNA EF-1α-Puro | MTCH1 dual | GCGGCACCGCCGCGAGCCCA, GAGCCCAGGGCGCCACTTCC | Guna et al. (*8*) |
| pAG59 | pU6-sgRNA EF-1α-Puro-T2A-BFP | Non-targeting | GAACGACTAGTTAGGCGTGTA | Guna et al. (*8*) |
| pAG440 | pU6-sgRNA EF-1α-Puro-T2A-BFP | MTCH2 single | GACGGAGCCACCAAGCGACC | Guna et al. (*8*) |
| pAG514 | pU6-sgRNA EF-1α-Puro-T2A-BFP | SLC25A46 dual | GCAGGGATGAGGGGTTACTG, GAGCACACACAGCTTCCCGT | this study |
| pAG515 | pU6-sgRNA EF-1α-Puro-T2A-BFP | MTCH1 dual | GCGGCACCGCCGCGAGCCCA, GAGCCCAGGGCGCCACTTCC | Guna et al. (*8*) |
| pMH458 | pU6-sgRNA EF-1α-Puro-T2A-BFP | ARMC1 #1 | GGGAAGCGGCCCCTGTACCG | this study |
| pMH459 | pU6-sgRNA EF-1α-Puro-T2A-BFP | ARMC1 #2 | GCAGAGGCGTCAGGTCACCT | this study |
| pAG452 | pU6-sgRNA EF-1α-Puro-T2A-BFP | DNAJC11 | GTGTCCCTGACGCGGATCAC, GGGGCCAGTGATCCGCGTCA | this study |

**Table S4.** UniProtKB-based MTCH homolog identification

| Protein family | PTHR10780:SF20 (MTCH2) | PTHR10780:SF3 (MTCH1) | PTHR10780:SF18 (MTCH) |
|---|---|---|---|
| Species | | # of sequences | |
| *H. sapiens* (human) | 4 | 23 | 0 |
| *M. musculus* (mouse) | 5 | 9 | 0 |
| *X. tropicalis* (frog) | 4 | 0 | 0 |
| *S. salar* (salmon) | 3 | 3 | 0 |
| *C. elegans* (nematode) | 0 | 0 | 2 |
| *D. melanogaster* (fly) | 0 | 0 | 14 |
| *N. vectensis* (anenome) | 0 | 0 | 3 |
| *S. rosetta* (choanoflagellate) | 0 | 0 | 1 |
| *C. owczarzaki* | 0 | 0 | 1 |
| *S. arctica* | 0 | 0 | 1 |

**Table S5.** UniProtKB-based OM insertase homolog identification

| Protein family | PTHR10780 (MTCH) | PTHR28241 (Mim1) | IPR043645 (pATOM36) | PTHR36074 (At5g55610) |
|---|---|---|---|---|
| Clade (taxid) | # of sequences | | | |
| Euglenozoa (33682) | 0 | 0 | 42 | 0 |
| Heterolobosea (5752) | 0 | 0 | 0 | 0 |
| Chlorophyta (3041) | 0 | 0 | 0 | 45 |
| Streptophyta (35493) | 0 | 0 | 0 | 682 |
| Rhodophyta (2763) | 0 | 0 | 0 | 0 |
| Alveolata (33630) | 0 | 0 | 0 | 0 |
| Stramenopila (33634) | 0 | 0 | 0 | 94 |
| Amoebozoa (554915) | 0 | 0 | 0 | 0 |
| Fungi (4751) | 0 | 1235 | 0 | 0 |
| Fonticula (691882) | 0 | 0 | 0 | 1 |
| Capsaspora (192874) | 0 | 0 | 0 | 1 |
| Choanoflagellates (28009) | 2 | 0 | 0 | 0 |
| Metazoans (33208) | 2836 | 0 | 0 | 0 |

**Table S6.** AlphaFold model confidence metrics.

| | **figures** | **source** | **accession** | **<pLDDT>** | **<PAE>** | **pTM** |
|---|---|---|---|---|---|---|
| *H. sapiens* MTCH2 | S1, S10 | Uniprot | Q9Y6C9 | 80.2 | 7.01 | 0.85 |
| *D. melanogaster* MTCH | S1, S10 | Uniprot | Q9V3Y4 | 82.1 | 7.63 | 0.84 |
| *C. owczarzaki* MTCH | S1, S10 | Uniprot | A0A0D2WXZ5 | 74.9 | 11.0 | 0.77 |
| *N. vectensis* MTCH | S10 | NCBI | XP_032228681 | 73.2 | 9.55 | 0.78 |
| *H. sapiens* MTCH1 | S10 | Uniprot | Q9NZJ7 | 70.5 | 15.9 | 0.69 |
| *T. brucei* pATOM36 | 5 | Uniprot | Q582I5 | 80.2 | 7.01 | 0.62 |
| *A. thaliana* At5g55610 | 5 | Uniprot | Q9FM77 | 66.5 | 18.03 | 0.59 |
| *H. sapiens* SLC25A4 | S1 | Uniprot | P12235 | 87.0 | 4.15 | 0.9 |
| *H. sapiens* UCP1 | S1 | Uniprot | P25874 | 81.5 | 6.53 | 0.84 |

**Movie S1.** Morph between WT MTCH2 and $MTCH2^{K25E,Y235A,V238D}$ conformation shown in space-filling representation.
